# Glacier retreat decreases plant-pollinator network robustness over space-time

**DOI:** 10.1101/2024.06.21.600080

**Authors:** Matteo Conti, Pierfilippo Cerretti, Andrea Ferrari, Paolo Gabrieli, Francesco Paone, Carlo Polidori, Gianalberto Losapio

## Abstract

Glaciers are retreating worldwide at an ever-increasing rate, exposing new ice-free areas to ecological succession. This process leads to changes in biodiversity and potentially to species interactions. However, we still have a limited understanding of how glacier retreat influences species interaction networks, particularly the structure and robustness of mutualistic networks. After reconstructing plant-pollinator networks along a glacier foreland, we address the effects of glacier retreat on pollination network structure and robustness. Our results show that the prevalence of different network motifs changes over space-time. With glacier retreat, pollination networks shift from being highly connected with specialist interactions to loosely connected with generalist interactions. Furthermore, network robustness decreased with glacier retreat. Despite the turnover of plant species, we find that structural roles played by different plant species stay constant over space-time. Our findings suggest that glacier retreat pushes pollination networks towards a loss of specialist interactions and low robustness, leading to increased fragility in the long term. Monitoring network motifs may provide valuable insights into the ability of novel pollination networks to withstand disturbances and preserve functionality in the face of glacier extinction.

## 1. Introduction

Glacier retreat is one of the most emblematic symptoms of anthropogenic climate change (Marzeion et al. **2014**, Roe et al. **2017**). According to recent projections (Rounce et al. **2023**), 49±9% of the world’s glaciers may disappear at the current global warming rate by 2100. Glacier retreat has a wide range of hydrological and geomorphological consequences on mountain ecosystems, impacting water quality and availability in high-altitude environments (Moore et al. **2009**) and increasing the frequency and magnitude of natural hazards (Keiler et al. **2010**). However, the effects of glacier retreat on species interactions are still poorly understood and difficult to predict.

As glaciers retreat, new ice-free areas are progressively exposed to colonisation by microbes, plants, and animals (Whittaker **1993**, Chapin et al. **1994**, Körner and Spehn **2002**, Bradley et al. **2014**, Tampucci et al. **2015**, Eichel **2019**). This process leads to a primary succession, with the communities closest to the glacier front being the youngest and those further away the oldest (Matthews **1992**). Therefore, given the possibility of working with a space-time substitution approach, studying pollination networks along a glacier foreland means not only assessing differences in space but also estimating how the system will evolve over time (Losapio et al. **2015**). Following glacier retreat, there is an initial increase in plant and pollinator diversity, leading to a local increase in biodiversity (Cauvy-Fraunié and Dangles **2019**, Losapio et al. **2021a**, Tu et al. **2024**). However, this is a just a temporary phase. When the glacier completely recedes from the landscape, the entire glacier foreland reflects the final stage of succession, leading to a significant reduction in local and regional biodiversity (Stibal et al. **2020**; Losapio et al. **2021a**, Tu et al. **2024**). Yet, we still know very little about the implications these changes entail for the structure of pollination networks and their ability to withstand disturbances.

The structure of pollination networks is typically analysed using one or more indices that capture various aspects of network architecture (Simmons et al. **2019a**). Network motifs provide a more sensitive approach to characterizing network structure, particularly in response to changes in local interactions (Simmons et al. **2019a**, **2019b**). Network motifs are sub-graphs defined by recurring patterns of interactions between small sets of species that are over-represented in a network as compared to random expectation (Milo et al. **2002**, Alon **2007**, Simmons et al. **2019a**, **2019b**, Lanuza et al. **2023**). Motifs are the building blocks upon which the network is assembled, and they may represent a framework in which specific functions are achieved efficiently within the network (Milo et al. **2002**, Alon **2007**, Simmons et al. **2019a**). A broad range of network configurations may exhibit similar values for indices like nestedness or modularity. However, far fewer configurations share similar motif compositions (Simmons et al. **2019a**).

Network motifs are used in two main ways: to calculate the frequency of occurrence of different motifs in a network and to quantify the structural role of species within motifs (Simmons et al. **2019b**, Lanuza et al. **2023**, Cirtwill et al. **2024**). We used both approaches to investigate pollination network structure following glacier retreat.

Regarding the first approach, we are interested in seeing how motifs’ prevalence changes as a result of glacier retreat. For instance, we expect to observe a shift from motifs rich in specialist interactions to motifs dominated by generalist interactions (both on the part of plants and pollinators), as glacier retreat has been shown to favour plant and pollinator generalisation (Roberts et al. **2011**, Cauvy-Fraunié and Dangles **2019**, Bogusch et al. **2020**) and push mutualistic networks from facilitation to competition (Losapio et al. **2021a**). Glacier retreat is facilitating the shrinkage of grassland environments and the upward shift of the treeline (Körner and Hiltbrunner **2021**). This process forces grassland plant species to shift their local range, resulting in competition for resources and potentially altering the ecosystem’s composition (Parolo and Rossi **2008**, Albrecht et al. **2014**, Alexander et al. **2015**, Zimmer et al. **2018**, Inouye **2020**, Losapio et al. **2021a**). Upward colonisation of plant species results in specialist pioneer species – living close to the glacier – to compete with generalist species adapted to later succession al stages (Losapio et al. **2021a**). The most immediate consequence of this process is the loss of suitable habitat for pioneer species (Körner and Hiltbrunner **2021**, Losapio et al. **2021a**), which are forced to go locally extinct due to increased competition (Pertoldi and Bach **2007**, Losapio et al. **2021a**).

Moreover, the frequencies of motifs in ecological networks have been associated with community stability, with some motifs being more frequently observed in stable networks than in unstable ones (Borrelli et al. **2015**). One of our objectives is therefore to understand the connection between network robustness and motifs’ prevalence. We investigate ‘robustness’ as the ability of pollination networks to resist extinction cascades (Dunne et al. **2002**, Landi et al. **2018**). This choice was made because robustness is a fundamental feature of complex biological systems (Kitano **2004**), such as pollination networks (Olesen et al. **2006**), and because the core element that characterises it, i.e. the system’s resistance to species loss (Dunne et al. **2002**), fits well with the actual scenario that climate change is imposing on alpine ecosystems.

Regarding the second approach, species roles within the network can be defined by tracking the frequency with which each species occupies each position within each motif (Cirtwill et al. **2018**). Some motifs have been shown to influence the size and population dynamics of focal species, indicating that a species’ involvement in a specific set of motifs may correspond to a distinct functional role within the community (Cirtwill et al. **2018**). One of our objectives is to understand whether plant species structural roles within motifs are altered by glacier retreat (environmental component) or are only driven by differences among species (evolutionary component). To the best of our knowledge, there has been no prior investigation into the effects of glacier retreat on pollination network robustness and its relationship with network motifs. Therefore, this study aims to provide the first insights to fill this knowledge gap. Here, we addressed how the retreat of Mont Miné glacier in the Swiss Alps impacts pollination network structure and robustness, testing the following hypotheses: (I) glacier retreat alters network motifs’ patterns pushing pollination networks towards a loss of specialist interactions, (II) robustness, in the long term, decreases following glacier retreat and is linked to motifs’ prevalence, and (III) motif roles of plant species are determined both by environmental conditions and evolutionary histories.

## 2. Materials and methods

### 2.1 Study area

Fieldwork was carried out along Mont Miné glacier foreland in the Vallon de Ferpècle, Val d’Hérens, Switzerland (centre of the study area: 46°3’33.646’N/7°32’54.550’E). Mont Miné glacier covers an area of 9.67 km², has a length of 5.28 km and an altitudinal range from 2122 to 3804 m a.s.l. (2015 glacier state) (Paul et al. **2020**, Nicolussi et al. **2022**). At the peak of the Little Ice Age (LIA, c 1860), Mont Miné and the adjacent Ferpècle glacier were connected, covering an area of 26.9 km². The two glaciers remained connected until 1956. Subsequently, Mont Miné retreated by 842 m up to 2019, marking a total retreat of approximately 2.53 km since the LIA maximum (Bezinge **2000**, Curry et al. **2006**, Nicolussi et al. **2022**).

Through the analysis of geochronology reports, it is possible to reconstruct the different stages of glacier retreat in a ‘chronosequence’ by calculating an age (years since the glacier retreated) for the terrains and their associated communities (Walker et al. **2010**, Losapio et al. **2021a**, Ficetola et al. **2021**), performing a so called ‘space-for-time substitution’ (Foster and Tilman **2000**, Walker et al. **2010**, Losapio et al. **2015**). Time since deglaciation is the primary driver of community richness and structure, particularly over large spatial scales (Foster and Tilman **2000**, Ficetola et al. **2021**). Glacier forelands represent therefore a unique natural model system for studying the impacts of climate change on alpine communities (Albrecht et al. **2010**, Losapio et al. **2021a**). However, some caution must be exercised since time is not the only driver in shaping communities along successions (Raffl et al. **2006**, Johnson and Miyanishi **2008**, Garibotti et al. **2011**, Ficetola et al. **2021**). For instance, plant species richness and diversity are strong predictors of plant-pollinator interaction diversity (Tu et al. **2024**).

Based on existing geochronology of the Mont Miné glacier (Curry et al. **2006**, Lambiel et al. **2016**, Nicolussi et al. **2022**) and our reconstruction of historical cartography (‘Journey through time’ tool at https://map.geo.admin.ch), we divided the Mont Miné glacier foreland in four stages: Stage 1, Stage 2, Stage 3 and Stage 4, representing terrain deglaciated since 1989, 1925, 1900 and 1864, respectively (Figure 1) (Curry et al. **2006**, Lambiel et al. **2016**, Nicolussi et al. **2022**).

**Figure 1.**
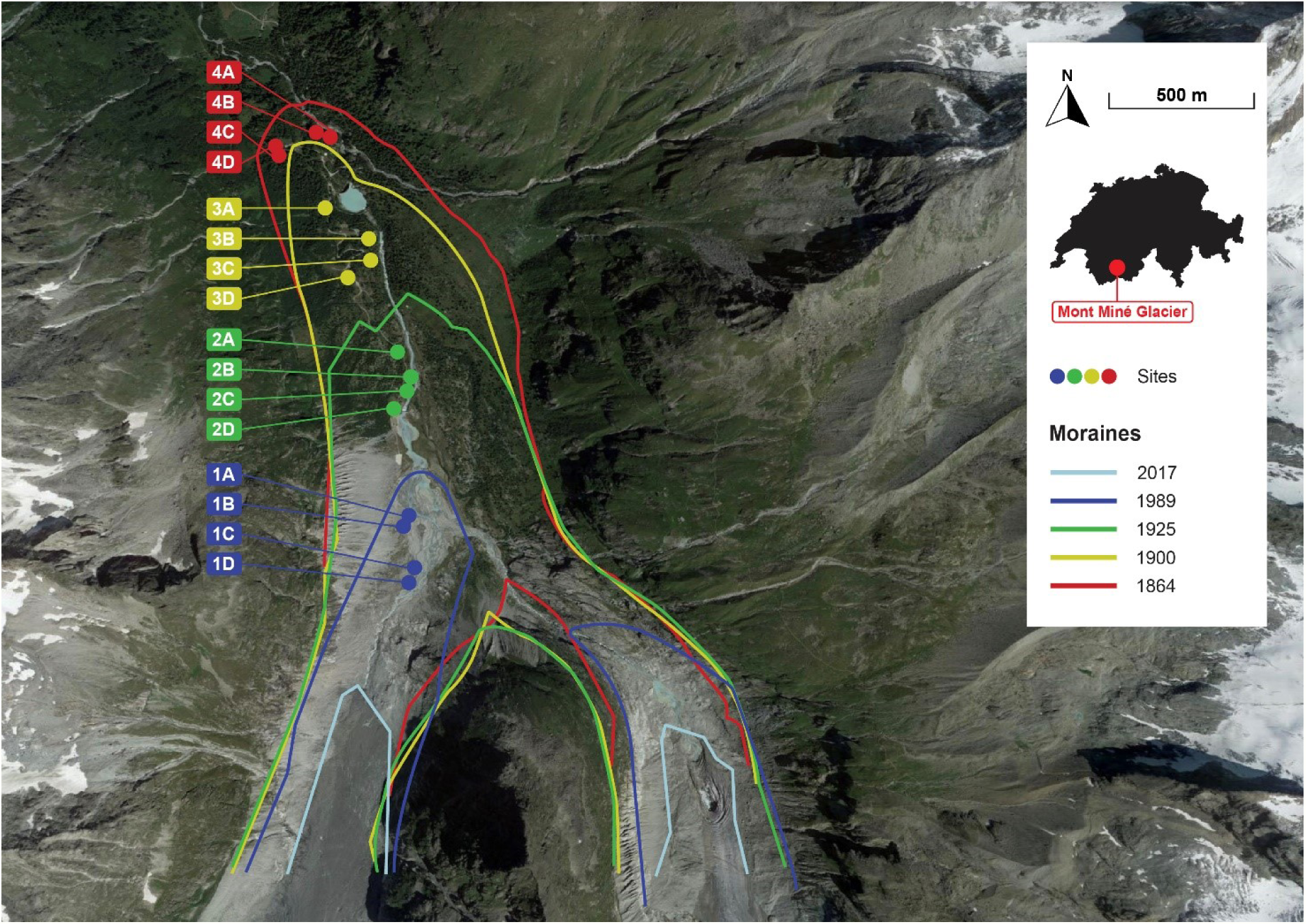
Satellite image of Vallon de Ferpècle (Google Earth 10.48.0.2, 08/07/2016) with geochronological reconstruction of Mont Miné (left) and Ferpècle (right) glaciers’ moraines.

We defined the time since deglaciation (x) of each stage (i) as 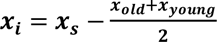, where 𝒙_s_ is the year of sampling and 𝒙_𝒐𝒍𝒅_ and 𝒙_𝒚𝒐𝒖𝒏𝒈_ are the ages of the moraines delimiting the i-th stage (Losapio et al. **2021a**, Tu et al. **2024**). Time since deglaciation is 17 years for Stage 1, 66 years for Stage 2, 111 years for Stage 3 and 141 years for Stage 4. We randomly selected 4 sampling sites per stage, for a total of 16 sites (Figure 1). Sites were chosen only on the west side of the river, ensuring minimal variation in elevation gradient (1961 to 2000 m a.s.l) and avoiding as much as possible the impact of human construction activities related to the Barrage de Ferpècle (Tu et al. **2024**). In each site, we installed two orthogonal transects measuring 25 m in length and 1 m in width, corresponding to 50 m² per site (Gibson et al. **2011**, Grange et al. **2021**, Martínez-Núñez et al. **2022**, Tu et al. **2024**) (supplementary information, Figure S2b).

### 2.2 Sampling

Sampling took place during subalpine flowering season, from June the 15^th^ to July the 15^th^ 2023.

Within each transect, we surveyed blooming plant species and recorded plant-pollinator visitations. Visitations in which flower visitors came into direct contact with flower anthers or stigmas were considered as pollination interactions (Gibson et al. **2011**). We recorded interactions by collecting flower-visiting insects using an entomological aspirator or by sweep netting, depending on the size of the specimen. We then placed collected pollinators in labelled tubes with 70% ethanol for subsequent identification. We limited the collection of specimens to those necessary for species identification and proceeded with recording visitations by direct observation.

To standardise observations over time, we walked transects at a slow and steady pace for 15 minutes per orthogonal line, for a total of 30 minutes per transect (Martínez-Núñez et al. **2022**). Sampling sessions were carried out on sunny and low-windy days from 8:30 a.m. to 5:30 p.m. (Fijen et al. **2018**, Martínez-Núñez et al. **2022**, Mahon and Hodge **2022**). We sampled each site eight times, for a total of *n*=128 temporal replicates. To minimise the impact of sampling time on pollinator activity, we randomised the sampling order of sites within a day and within a round of replicates such that every site was sampled at all possible times of the day (Martínez-Núñez et al. **2022**).

We performed plant identification during fieldwork at species level according to Flora Helvetica (Lauber and Wagner **1998**, updated version on https://www.infoflora.ch). Collected insect specimens were first identified at genus- or species-level whenever possible using taxonomic keys (Rognes **1991**, Cerretti **2010**, Cerretti et al. **2012**, Gregor et al. **2016**, Cappellari et al. **2018**, Falk **2019**, Michez et al. **2019**, Rasmont et al. **2021**, Museum of Zoology - Sapienza University of Rome), and then photographed using a stereomicroscope.

We performed all statistical analyses considering pollinators up to family level, because this was the lowest possible level for which high confidence in identification was maintained. We created a citizen science project on iNaturalist platform to gather further taxonomic information, for data sharing and for public outreach (https://www.inaturalist.org/projects/pollinator-diversity-at-ferpecle-glacier-ecosystems).

We observed a total of 108 blooming plant species belonging to 28 families. Regarding pollinators, we identified 65 families belonging to 8 orders. In total, we recorded 2180 plant-pollinator interactions.

### 2.3 Data analysis

All statistical analyses were performed in ‘R’ software (version 4.3.1) (R Core Team **2023**). When addressing network motifs properties and network robustness, we adopted a null model approach. This allowed us (i) to ensure that that the measures considered in the network analysis were informative properties of the networks and did not simply emerge by chance, and (ii) to compare network properties among networks differing in other properties than target ones, such as matrix size and filling (Vázquez and Aizen **2003**, **2006**, Dormann et al. **2009**). To address network motifs analyses, we randomised the networks 100 times under four different null model approaches (varying in their level of conservativism). Here, we’ll present considerations for the main reference model, but details regarding the other three models can be found in the supplementary information. Results did not qualitatively change with different null model approaches, except for the most conservative one, which gave a much lower total percentage of above-threshold motifs.

#### 2.3.1 Motifs’ prevalence

To understand how glacier retreat affects network motifs, we pooled interactions at the site level and calculated the frequency of occurrence of all 44 possible motifs up to 6 nodes (supplementary information, Figure S1) within sites’ networks. Motifs’ occurrences were converted to relative frequencies by expressing counts as a proportion of the total number of motifs in the network (Mora et al. **2018**, Simmons et al. **2019b**).

For each null model iteration, motifs’ frequency of occurrence was calculated as well. To test whether observed motifs’ frequency significantly differ from that of null model networks, we performed a one-sample Z-test on each motif within each site. For motifs above threshold, we can reject the null hypothesis that frequencies calculated on observed networks do not differ from chance expectation. The prevalence of these motifs is informative of specific patterns of interactions that are not expected by chance alone. The frequency of the remaining motifs could just arise by chance as a product of sampling effort or network size.

On the obtained set of motifs’ frequencies in observed networks, we performed a Non-metric Multidimensional Scaling (NMDS). We had *N* = 44 variables (i.e. motifs’ frequencies) associated to *n* = 16 objects (i.e. sites). To construct the distance matrix, we considered Bray-Curtis dissimilarities (Bray and Curtis **1957**). On the resulting ordination plot, NMDS1 (i.e. the coordinates of the axis that explains most of the variance) represent the gradient of connection of motifs: more negative NMDS1 values are associated with highly connected motifs where specialist pollinators have greater complementarity in the specialist plants they visit, while more positive NMDS1 values are associated with loosely connected motifs where generalist pollinators compete for generalist plants (Simmons et al. **2019b**).

To test whether stages were significantly different in terms of sets of motifs, we performed a Permutational Multivariate Analysis of Variance (PERMANOVA), also checking for multivariate homogeneity of groups’ dispersion.

#### 2.3.2 Network robustness

To calculate network robustness, we built a function in ‘R’ software that: (i) performs a secondary extinction cascade on a pollination network, (ii) interpolates an extinction function based on this cascade, and (iii) calculates robustness as the area under the extinction curve. The secondary extinction cascade occurs by removing plant species from the network and counting how many pollinator families are left with at least one link. This process is iterated until all plants and pollinators are left with no links. Coordinates of the extinction cascade (number of extinct plants and number of remaining pollinators at each iteration) are scaled to 1 and a function is interpolated to fit the resulting curve. This extinction curve is the ATC (Attack Tolerance Curve) (supplementary information, Figure S3) as devised by Memmot et al. (**2004**) and Burgos et al. (**2007**). We defined robustness (*R*) as the area under the ATC, i.e. its definite integral from 0 to 1 (Burgos et al. **2007**, Sheykhali et al. **2020**). A robustness value close to 1 is associated with a very robust network and corresponds to an ATC that decreases mildly. On the other hand, a robustness value close to 0 is associated with a very fragile network and corresponds to an ATC that decreases abruptly (Burgos et al. **2007**). Although this model relies on static assumptions such as lack of rewiring, it provides a solid estimation of community resilience.

We built two robustness models: the test model and the null model. In the test model, plant species go extinct with a probabilistic approach. Plant species have different extinction probabilities based on their distribution, which accounts both for coverage and habitat specificity along the glacier foreland. The narrower and more centred around the early stages a species’ distribution, and the lower its mean coverage, the higher the chances that the robustness function extracts the corresponding species. In the null model (or random model), all plant species have the same probability of becoming extinct (and thus being extracted by the robustness function) (Memmot et al. **2004**, Burgos et al. **2007**).

In both the test model and the null model, we pooled interactions at the site level (*n* = 16) and the final value of robustness was calculated as the mean value resulting from the extinction cascade iterated 1000 times per site.

To test whether the test model was significantly different from the null model, for each site we performed two-sample Z-tests. Z-score calculation was impossible for sites 4A and 4B, as the averages of the two models are identical and the standard deviation for both is zero. That’s because these sites comprise solely *Rhododendron ferrugineum* as plant species and their extinction cascade always repeats itself across iterations.

To test the effects of glacier retreat on robustness and to test whether differences in motifs’ prevalence between stages could predict robustness, we fitted a beta regression model. This choice was made because beta regression is a very effective and flexible approach in modelling response variables between 0 and 1. We included robustness as response variable, and quadratic time since deglaciation and linear NMDS1 coordinates of motifs’ ordination as predictors. We then performed Type-II ANOVA to assess the significance of the contribution of the individual predictors to the observed variance.

To find out whether the assumptions of the test model affected in some way the relationship between robustness and the predictors, we fitted a linear model (LM) using Z-scores as response variable, and time since deglaciation and NMDS1 coordinates as predictors. The relationship between test model robustness or null model robustness and the predictors gives us an ‘absolute’ trend of network resistance following glacier retreat, while the relationship between Z-scores and the predictors gives us an indication of how relevant the assumptions of the test model are in the way robustness relates to the predictors.

Finally, we tested the collinearity of the predictors using a linear model (LM) with time since deglaciation as response variable and NMDS1 coordinates as predictor.

#### 2.3.3 Plant species’ roles

To investigate the role of plant species within network motifs, we pooled interactions at the stage level and calculated plant species contributions to motifs’ node positions. For a given species in a given position in a motif, contribution was defined as the sum of focal species’ interactions divided by the sum of all interactions in that motif (Simmons et al. **2019b**). We considered all possible positions within motifs up to 5 species (supplementary information, Figure S1).

For each null model iteration, plant species contributions were calculated as well. To test whether contributions calculated on observed networks significantly differ from null models, we performed one sample Z-tests calculating Z-scores for each possible node position up to 5 species in each stage. Above-threshold node positions within motifs are informative of specific patterns of interactions that are not expected by chance alone.

Along the same lines of what we did for motifs’ frequencies, we performed an NMDS on plant species’ contributions (in this case *N* = 23, i.e. positions’ contributions, and *n* = 133, i.e. plant species with duplicates across different stages).

To test whether contributions of different stages (i.e., environmental component) or species (i.e., evolutionary component) were significantly different, we performed PERMANOVA and checked for multivariate homogeneity of groups’ dispersion. We also calculated plant species’ Shannon index of interactions (*H’*) as 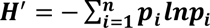, where 𝒑_𝒊_ is the proportional abundance of the i-th interaction (Bersier et al. **2002**, Dušek and Popelková **2017**), and performed PERMANOVA and dispersion analyses using plant species’ interaction diversity as a grouping factor.

To test whether differences in contributions to node positions between species could predict their interaction diversity, we fitted a Generalised Linear Model (GLM) using gamma as family, Shannon index of interactions (calculated for each plant species) as response variable, and quadratic NMDS1 coordinates of contributions’ ordination as predictor (Bingham and Fry **2010**). We chose gamma family because it is suitable for modelling strictly positive right-skewed continuous data.

We calculated motifs’ frequencies and plant species’ contributions using ‘bmotif’ package (Mora et al. **2018**, Simmons et al. **2019b**). We used ‘vegan’ package (Oksanen et al. **2022**, Oksanen **2023**) to build null models, to perform the NMDS, to perform PERMANOVA and to test the multivariate homogeneity of groups’ dispersions. One additional null model approach was implemented using ‘bipartite’ package (Dormann et al. **2023**). We built beta regression models using ‘betareg’ package (Cribari-Neto and Zeileis **2010**, Zeileis et al. **2022**). Finally, we performed Type-II ANOVA using ‘car’ package (Fox et al. **2023**).

## 3. Results

In Stage 1, which is the harsh and open environment close to the glacier, ants (Formicidae, ≈ 21% of visits) and bees (Apoidea families, ≈ 20% of visits) play a key role in the pollination of many plant species, including some pioneer plants such as *Epilobium fleischeri*, *Hieracium staticifolium* and *Linaria alpina*. In Stage 2, characterised mainly by blooming meadows, the relevance of ants (≈ 28% of visits) and bees (≈ 9% of visits) persists, while the importance of muscoid Diptera (≈ 26% of visits) and hoverflies (Syrphidae, ≈ 12% of visits) starts to increase. Mid-successional plants, such as *Saxifraga aizoides* and *Potentilla aurea*, acquire a predominant role, while typical late-stage species, such as *Rhododendron ferrugineum*, start appearing within the network. Moving to Stage 3, where the environment is a mix of grassland and woodland, muscoid Diptera (≈49% of visits) and hoverflies (≈ 13% of visits) take over and dominate interactions. Mid-successional plants such as *Potentilla aurea* and *Ranunculus spp.* are predominant and *Rhododendron ferrugineum* takes a large share of the observed interactions. Finally, with Stage 4, which is a closed woodland of *Larix decidua* with an undergrowth dominated by very few species of Ericaceae, such as *Rhododendron ferrugineum*, *Vaccinium spp*. and *Empetrum nigrum*, pollination is dominated by muscoid Diptera (≈ 51% of visits) and bees (Apidae, ≈ 28% of visits), and most interactions are taken by *Rhododendron ferrugineum*. Details regarding pollinator families and their abundances can be found in the supplementary information (Figure S5 and Table S5).

### 3.1 Motifs’ prevalence

We found that approximately 45% of network motifs across all stages and sites were significantly over-represented in observed networks compared to null model networks (supplementary information, Table S1).

Following the NMDS ordination (Figure 2), PERMANOVA revealed that different stages are significantly different in terms of prevalence of motifs (Table 1). Stages were also different in terms of homogeneity of variances (Table 1). Following glacier retreat, we observed a shift from a prevalence of motifs’ rich in specialist interactions (associated with the left part of the ordination plot) to ones dominated by generalist interactions (associated with the right part) (Figure 2).

**Figure 2.**
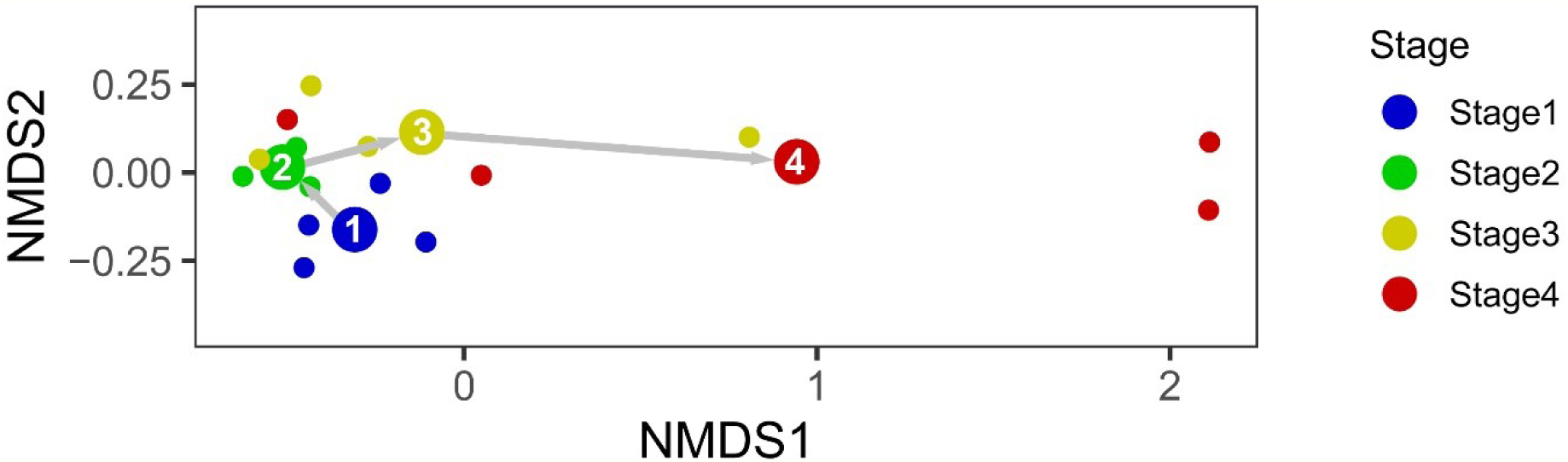
Non-metric Multidimensional Scaling (NMDS) ordination plot of normalised frequency of motifs. Small dots represent different sites within stages; big dots with numbers are stages’ centroids. Grey arrows show the observed shift along ordination axes.

**Table 1.** Summary statistics for PERMANOVA and homogeneity of variances analyses. Further details and groups’ pairwise comparisons can be found in the supplementary information.

| Analysis | Variables | Objects | Grouping factor | df | F | p |
| --- | --- | --- | --- | --- | --- | --- |
| PERMANOVA | Motifs' frequencies | Sites | Stage | 3 | 2.444 | 0.039 |
| Homog. Var. |  |  | Stage | 3 | 4.282 | 0.026 |
| PERMANOVA | Contr. to node positions | Plant species | Stage | 3 | 0.887 | 0.455 |
| Homog. Var. |  |  | Stage | 3 | 0.888 | 0.434 |
| PERMANOVA |  |  | Species | 73 | 0.839 | < 0.001 |
| Homog. Var. |  |  | Species | 73 | 1.493 | 0.236 |
| PERMANOVA |  |  | Interaction diversity | 2 | 182.650 | < 0.001 |
| Homog. Var. |  |  | Interaction diversity | 2 | 17.597 | 0.001 |

This is true from the perspective of both plants and pollinators, but the trend seems to be stronger for plants. Top 3 most representative motifs for each stage (the ones with highest mean frequency of occurrence across sites) are shown in Figure 3. Stage 4 is strongly characterised by loosely connected motifs with plant species’ generalist interactions (in which the protagonist is *Rhododendron ferrugineum*), while Stage 1, Stage 2 and Stage 3 are characterised by similar motifs that are more interconnected. These motifs also proved to be significantly over-represented in one or more sites within the corresponding stages.

**Figure 3.**
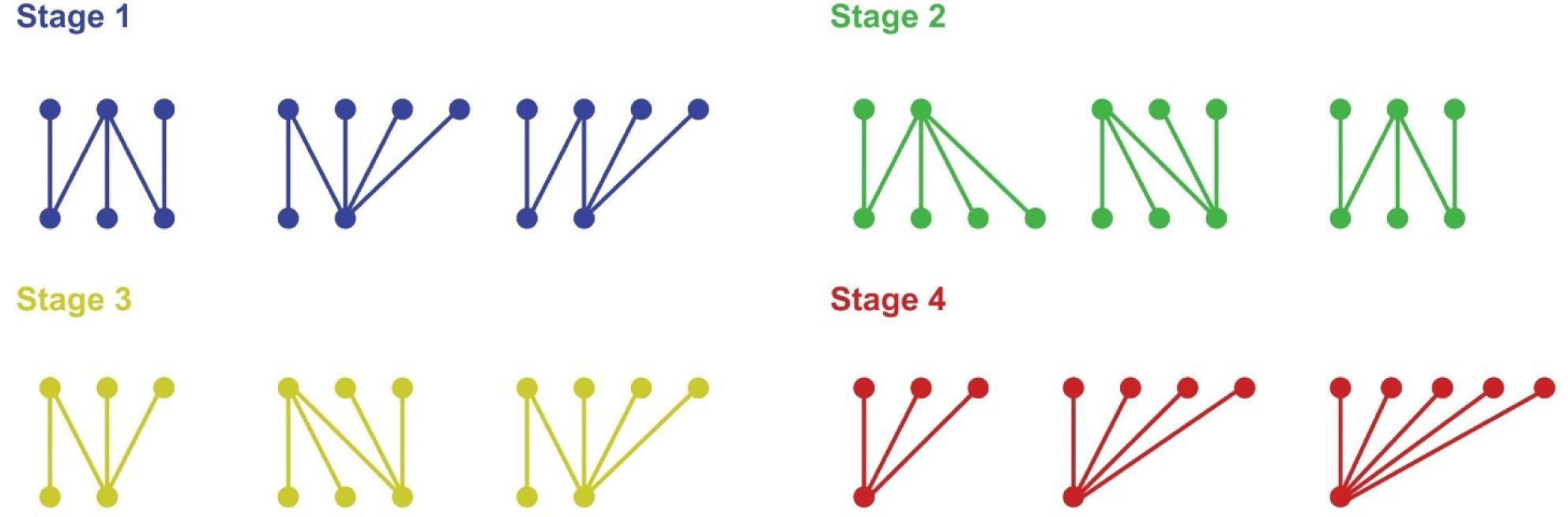
Most representative motifs. Lower nodes represent plant species, upper nodes insect families.

### 3.2 Network robustness

For each site, two-sample Z-tests revealed significant differences between observed and random networks in terms of network robustness (Z-test: |Z| > 1.96, *p* < 0.05; supplementary information, Table S3). Specifically, robustness measured under the extinction scenario of the test model was lower than that calculated under the null model in sites 3C and 4C (Z < −1.96, *p* < 0.05), while greater in the remaining 12 sites (Z > 1.96, *p* < 0.05).

Robustness was significantly negatively impacted by both time since deglaciation and NMD1 coordinates (Table 2). Robustness initially increases following glacier retreat, reaching its maximum in Stage 2, but in the long term it decreases, reaching its minimum in Stage 4 (Figure 4a). We observed a negative relationship between robustness and NMDS1 coordinates (Figure 4b). Negative values of NMDS1 are associated with higher robustness, while positive NMDS1 values are associated with lower robustness.

**Figure 4.**
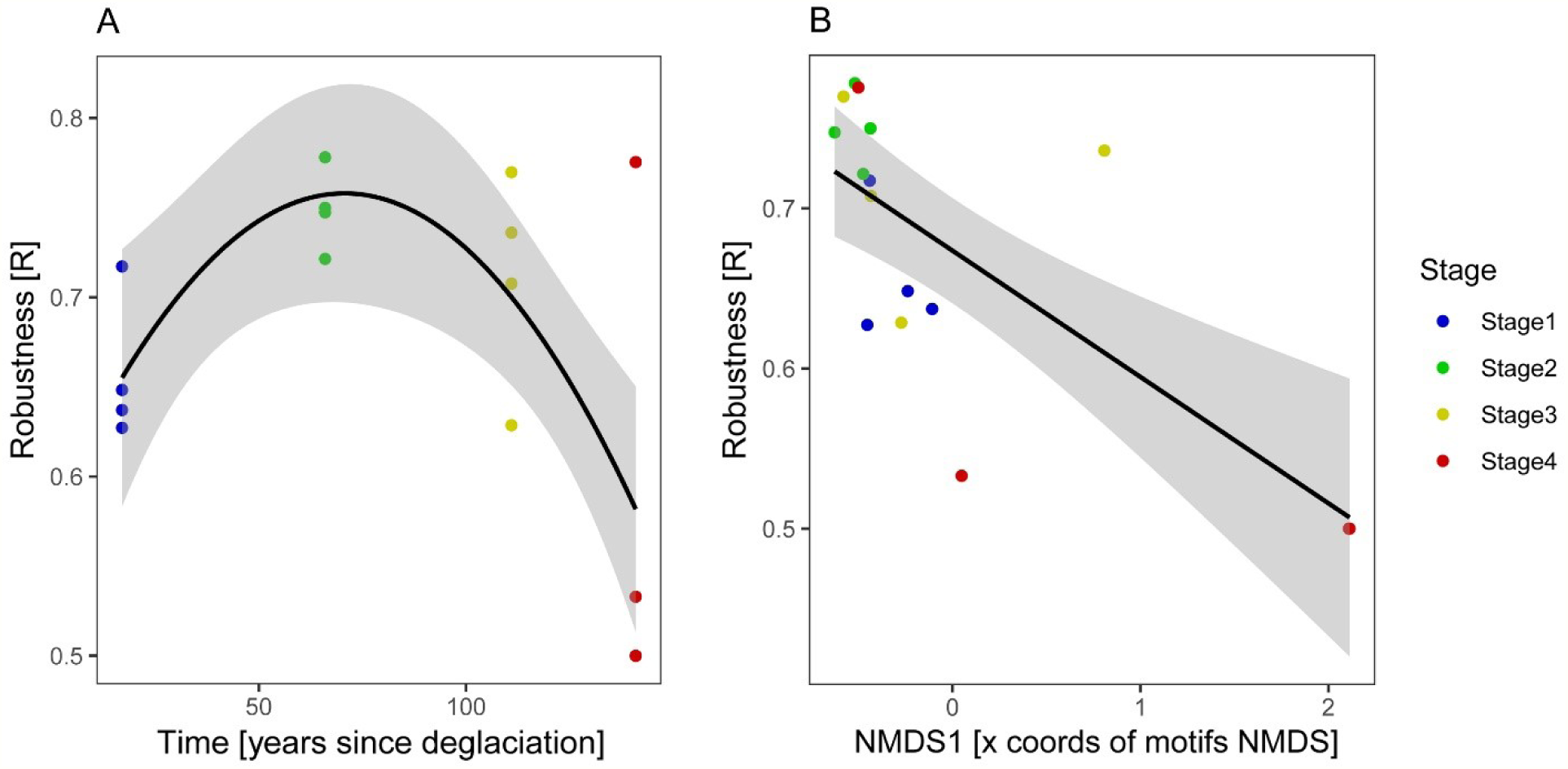
A) Relationship between test model robustness and time since deglaciation. B) Relationship between test model robustness and NMDS1 coordinates of motifs’ occurrences ordination. NMDS1 coordinates represent the degree of specialisation and connection of motifs: more negative values are associated with highly connected motifs with both specialist and generalist interactions, while more positive values are associated with loosely connected motifs dominated by generalist interactions. Fitted models are represented with black lines, while the 95% confidence interval is represented in grey.

**Table 2.** Summary statistics for fitted models. Note that the “NMDS1” of robustness models refers to motifs’ prevalence ordination, while the “NMDS1” of Shannon model refers to that of node positions. Further details can be found in the supplementary information.

| Response variable | Family | Predictors | Est | SE | Stat | p |
| --- | --- | --- | --- | --- | --- | --- |
| Robustness | Beta (link = logit) | poly(Time, 2) 1 | 0.066 | 0.285 | $z = 0.230$ | 0.818 |
| | | Poly(Time, 2) 2 | -0.671 | 0.276 | $z = -2.427$ | 0.015 |
| | | NMDS1 | -0.271 | 0.088 | $z = -3.074$ | 0.002 |
| Robustness Z-score | Gaussian (link = identity) | poly(Time, 2) 1 | -4.208 | 21.904 | $t = -0.192$ | 0.851 |
| | | Poly(Time, 2) 2 | -21.691 | 20.841 | $t = -1.041$ | 0.323 |
| | | NMDS1 | -15.919 | 14.371 | $t = -1.108$ | 0.294 |
| Shannon | Gamma (link = inverse) | poly(NMDS1, 2) 1 | 29.130 | 6.006 | $t = 4.850$ | < 0.001 |
| | | poly(NMDS1, 2) 2 | 9.281 | 2.565 | $t = 3.618$ | < 0.001 |

It’s important to underline here that network robustness decreases with time since deglaciation and NMDS1 coordinates regardless of how secondary extinction occurs, because Z-scores do not show any particular trend in response to the predictors (Table 2).

Type-II ANOVA revealed that both predictors are significant contributors to the variance of the response variable (𝑝_𝑡𝑖𝑚𝑒_ = 0.035, 𝑝_𝑁𝑀𝐷𝑆1_ = 0.002).

Time since deglaciation and NMD1 coordinates did not show significantly relevant collinearity, but only marginal (*p* = 0.050).

### 3.3 Plant species’ roles

We found that approximately 18% of node positions across all stages and species had a significantly over-represented contribution in observed networks compared to null model networks (supplementary information, Table S4).

PERMANOVA analyses indicated no significant differences in plant species’ contributions to node positions among stages (Table 1). Instead, we observed significant differences among species (Table 1). In this instance, stage did not exhibit the same predictive strength as observed with motifs’ occurrences, since a strong overlap between groups is observed (supplementary information, Figure S4a). Here, it is the species that differ from one another, regardless of the habitat they occupy (supplementary information, Figure S4b).

Interaction diversity proved to be a strongly significant grouping factor (Table 1) (Figure 5a). Modelling Shannon index of interactions (*H’*) in relation to NMDS1 coordinates, we observed a strongly significant inverse relationship (Table 2) (Figure 5b). More negative NMDS1 values are associated with higher interaction diversity, while more positive NMDS1 values are associated with lower interaction diversity. Differences in species structural roles within motifs can be explained in terms of interaction diversity.

**Figure 5.**
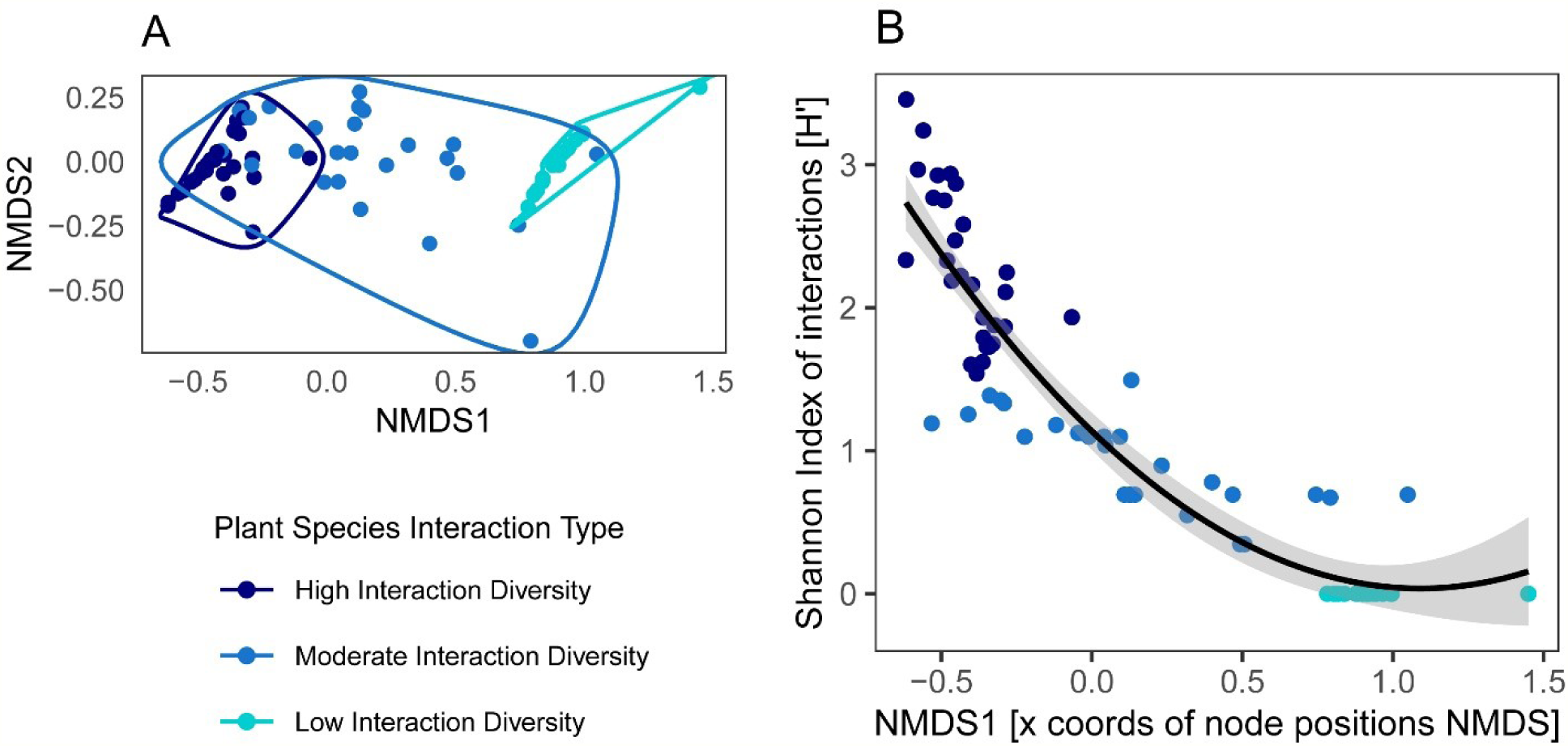
A) Non-metric Multidimensional Scaling (NMDS) ordination plot of plant species’ contribution to node positions within network motifs, grouped by plant species interaction diversity. Dots are plant species’ centroids. B) Relationship between Shannon index of plant species interactions and NMDS1 coordinates of node positions ordination. Fitted model is represented with a black line, while the 95% confidence interval is represented in grey.

## 4. Discussion

The impact of glacier retreat on biodiversity is increasingly documented, but still poorly investigated. Understanding how ecological networks develop and break down after glacier retreat is key for biodiversity conservation and management of novel, post-glacial ecosystems.

### 4.1 The impact of glacier retreat on network motifs

The prevalence of different network motifs changes with glacier retreat. We report a shift from more specialist motifs to generalist ones, supporting our hypothesis that the development of insect pollination networks in novel ecosystems is initially driven by specialisation and then by generalisation.

The observed specialist-to-generalist shift in motifs’ prevalence aligns with previous results on biogeography and species distribution highlighting that generalisation is favoured by glacier retreat, while specialisation can be disfavoured in light of current global warming trends (Erschbamer **2007**, Clavel et al. **2010**, Roberts et al. **2011**, Miller-Struttmann et al. **2015**, Cauvy-Fraunié and Dangles **2019**, Bogush et al. **2020**). Glacier retreat results in some alpine plant species being favoured (‘winners’), while others are disadvantaged (‘losers’) (Erschbamer **2007**, Cauvy-Fraunié and Dangles **2019**, Losapio et al. **2021a**). Most of the ‘losers’ are specialist species, adapted to proglacial environments, whereas the winners are generalist species moving upward from later stages of the succession (Cauvy-Fraunié and Dangles **2019**). It is therefore reasonable to assume that specialised plants (such as *Linaria alpina*) might face a higher risk compared to generalists (such as *Rhododendron ferrugineum*) (Losapio et al. **2021a**).

Differences between specialist and generalist plant species could impact pollinators too. For instance, in Europe, it has been argued that specialist bee and butterfly species are more vulnerable to the impacts of climate change compared to their generalist counterparts (Roberts et al. **2011**, Defra **2019**, Bogush et al. **2020**). Similar trends have been found for insects involved in other trophic interactions, such as parasitoids (Di Marco et al. **2023**).

Our results align with these considerations, suggesting that if a pollinator species manages to resist the species turnover that benefits generalists, it is inevitably forced to adapt by changing its altitudinal distribution or foraging behaviour (Miller-Struttmann et al. **2015**, Inouye **2020**).

However, given our lower taxonomic resolution, it is not immediate that our results would lead to such considerations at the species level and caution must therefore be exercised.

### 4.2 The impact of glacier retreat on network robustness

Network robustness initially increases after glacier retreat, but in the long term it sharply decreases, confirming our hypothesis. It has been shown that glacier retreat is a process that occurs in two separate phases: initially, the disappearance of the ice gives way to open grasslands, favouring pollination, but in the long term, once the closed forest rises in altitude and the grasslands disappear, it has a strong negative impact (Tu et al. **2024**). This trend has been shown with regard to plant diversity, pollinator diversity and plant-pollinator interaction diversity (Tu et al. **2024**), here we show that the same applies to robustness.

However, since the test model and the null model follow the same trend in relation to time since deglaciation and given that Z-scores do not follow any significant trend, we can conclude that robustness declines following glacier retreat regardless of how the extinction cascade occurs.

Moreover, our results suggest that differences in network robustness among stages can be explained in terms of motifs’ prevalence. Networks rich in motifs with more specialist interactions are associated with higher network robustness, while networks rich in motifs dominated by generalist interactions are associated with lower robustness. This may sound counterintuitive, since it has been shown that generalisation improves network robustness (Mello et al. **2011**, Astegiano et al. **2015**, Brosi **2016**). However, the observed shift from specialisation to generalisation following glacier retreat is not an increase in generalist interactions, but rather the progressive loss of specialist interactions. With a loss of specialist interactions due to glacier retreat, we expect to observe a corresponding loss of robustness, since many important bridges connecting network modules are lost.

Since motifs can predict network robustness, we suggest that studying motif patterns in pollination networks may provide valuable insights into their ability to withstand disturbances.

### 4.3 Plant species’ roles

Notably, we found evidence that roles plant species play within network motifs are mostly driven by their evolutionary history rather than environmental conditions. These differences are correlated with plant species interaction diversity. Based on these results, we expect to find: (i) similar node positions within motifs for the same species when comparing different specimens coming from different stages along the glacier foreland, (ii) similar node positions when comparing species with similar interaction diversity, and (iii) different node positions when comparing species with different degrees of interaction diversity.

For example, *Epilobium fleischeri* is a pioneer species that plays a crucial role in early-stage pollination and is characterised by high interaction diversity. *Linaria alpina* is another pioneer species, found only in Stage 1, characterised by low interaction diversity. *Rhododendron ferrugineum* is the dominant species in the latest stage and is characterised by high interaction diversity. The five most representative node positions within motifs (those with highest contribution) for these three species are shown in Figure 6. As can be seen, *E. fleischeri* and *R. ferrugineum* have the same pattern of node positions, despite belonging to opposite environments, while both differ from *L. alpina*.

**Figure 6.**
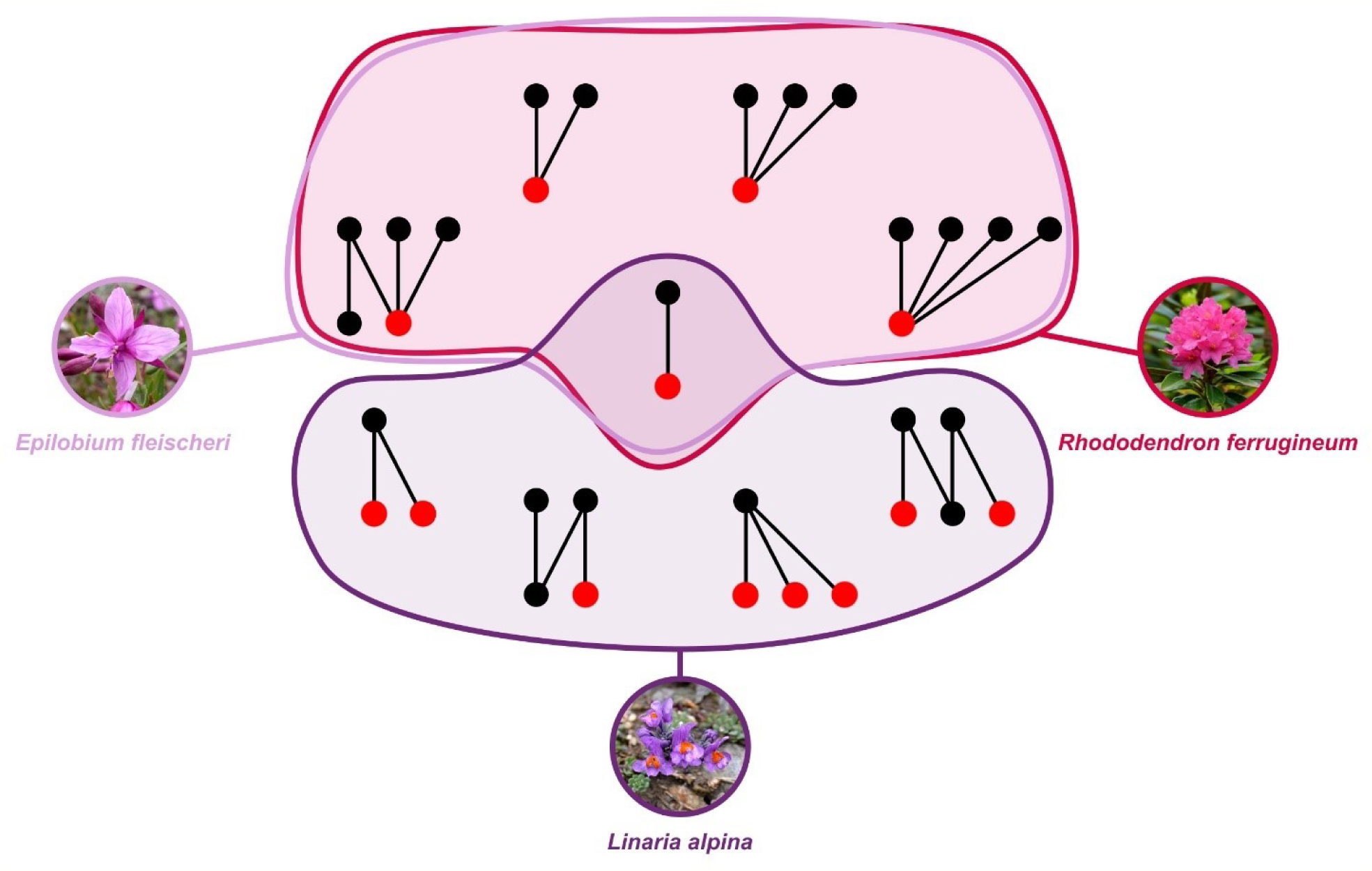
Most representative node positions for *E. fleischeri*, *L. alpina* and *R. ferrugineum*. Lower nodes represent plant species, upper nodes insect families. Red dots represent the positions occupied by plant species within motifs. If there is more than one highlighted position within a motif, it means that they are topologically the same position. Pictures rights belong to Barbara Studer (*E. fleischeri*), Bernard Dupont (*R. ferrugineum*, and Muriel Bendel (*L. alpina*) (©https://creativecommons.org/licenses/by-sa/4.0/).

The lack of differences in species’ roles across stages suggests that, along with taxonomic turnover, there should not be a corresponding ‘structural turnover’, since key roles are consistently filled by certain species. Although pollination networks are dynamic entities in both space and time in terms of species composition and interactions (Dupont et al. **2009**, Valdovinos et al. **2009**, Valverde et al. **2015**, Zografou et al. **2020**), it seems that many structural parameters remain relatively constant (Alarcón et al. **2008**, Dupont et al. **2009**), suggesting that species may be replaced by others with topologically similar roles (Dupont et al. **2009**). While climate change leads to a turnover of species and a loss of diversity, it may not necessarily imply that key roles and functions in pollination networks are lost as well. Early-stage species with high interaction diversity such as *Epilobium fleischeri* and *Hieracium staticifolium* show very similar roles as late-stage species such as *Rhododendron ferrugineum* and *Potentilla aurea*. The latter may replace the former due to climate change while maintaining unaltered pollination networks’ basic topological properties. However, given our taxonomic resolution, it is unclear whether and to what extent these species can interact with the same pollinator species, maintaining the network stable over time. Moreover, it’s important to underline that these considerations are driven by the sampling method we choose (visitation data), since flower-visitor data and pollen-load data provide complementary insights (Cirtwill et al. **2024**). We suggest implementing this study with pollen data, as plant species’ roles within motifs tend to be influenced by the sampling method (Cirtwill et al. **2024**).

In addition, we suggest that studying species roles within motifs can complement species-level network indices, such as centrality, in identifying key species in pollination networks. Those species with key roles and functions shall be the priority target of biodiversity conservation actions.

In conclusion, glacier retreat is a process that reshuffles plant-pollinator interactions, pushing network motifs towards a loss of specialist interactions, a dominance of generalist interactions, and lower robustness.

Motifs analysis could provide a means of prioritising species for conservation or restoration plans based on their roles and functions in the stability of biological communities (Cirtwill et al. **2018**). As motifs mediate the impacts of glacier retreat on pollination network robustness, we argue that understanding the assembly and breaking down of pollination network motifs is key for assessing the development of novel glacier ecosystems. The results of this study can therefore contribute to informing future conservation plans for mountain ecosystems facing ongoing climate change.

