## Supplementary Information for "Glacier retreat decreases plant-pollinator network robustness over space-time"

**This document includes:**
- Figures S1 to S5
- Tables S1 to S5
- 'R' code

#### Supplementary figures

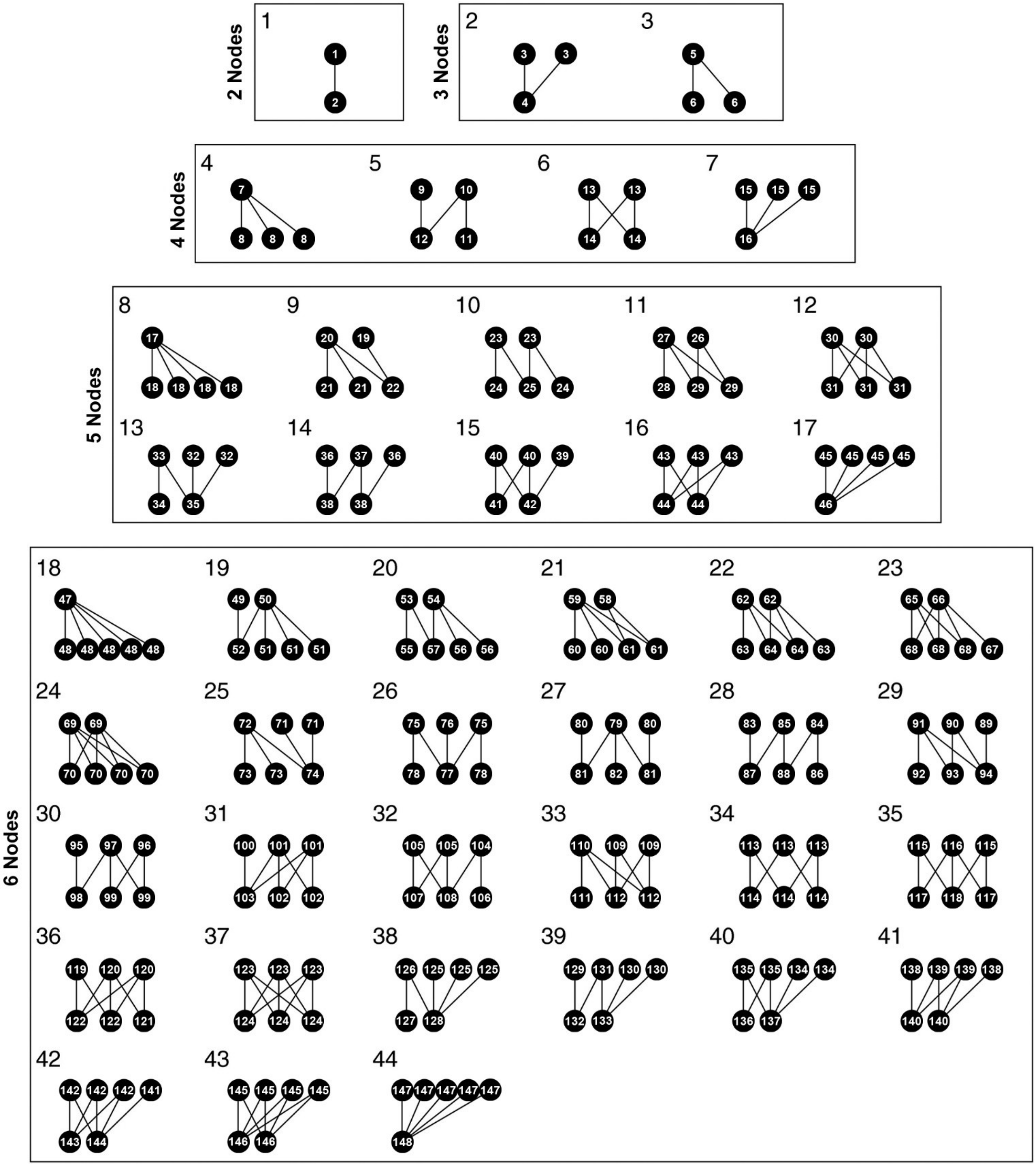

Figure S1. Bipartite network motifs up to 6 nodes. *Large numbers identify each motif. Small numbers represent the unique positions species can occupy within motifs. Lines between small numbers indicate undirected species interactions. This figure is adapted from Simmons et al. (2019a). Motifs are mirrored vertically with respect to Simmons et al. (2019a), since in our representation plants occupy the bottom nodes, while Simmons' algorithm places the lower level (in our case plants) in the top nodes when calculating motifs' occurrences and node positions.*

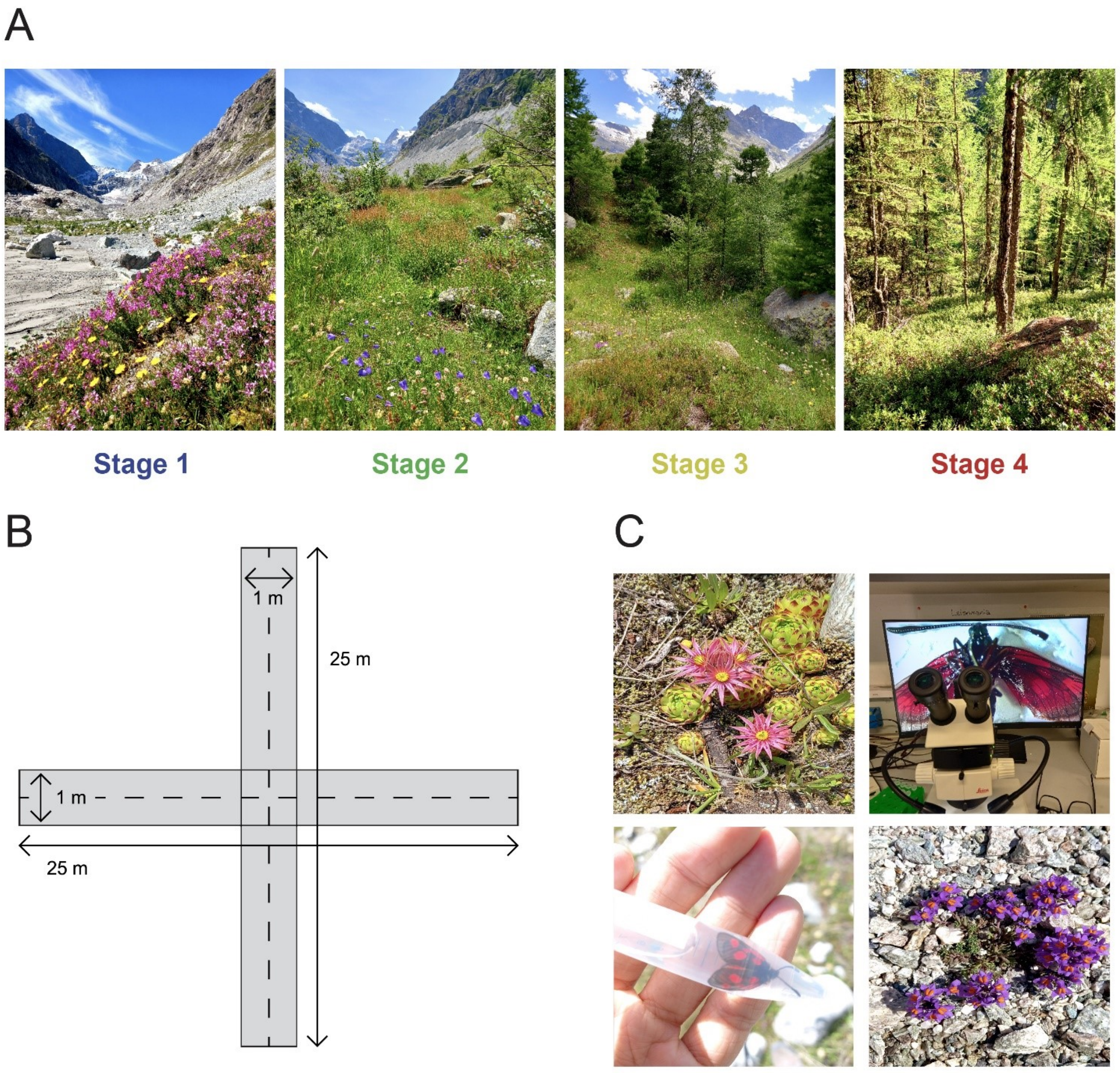

Figure S2. Stages, sampling method and activities. A) *A picture for each stage of the succession to show macroscopic environmental differences (pictures taken by Anonymous in July 2023).* B) *Schematic representation of sampling transect.* C*) Fieldwork and lab activities. Clockwise: a blooming Sempervivum montanum, Zygaena exulans seen through a stereomicroscope during labwork for identification, a blooming Linaria alpina, the same Zygaena specimen caught in a falcon tube during fieldwork (pictures taken by Anonymous between June, July and September 2023).*

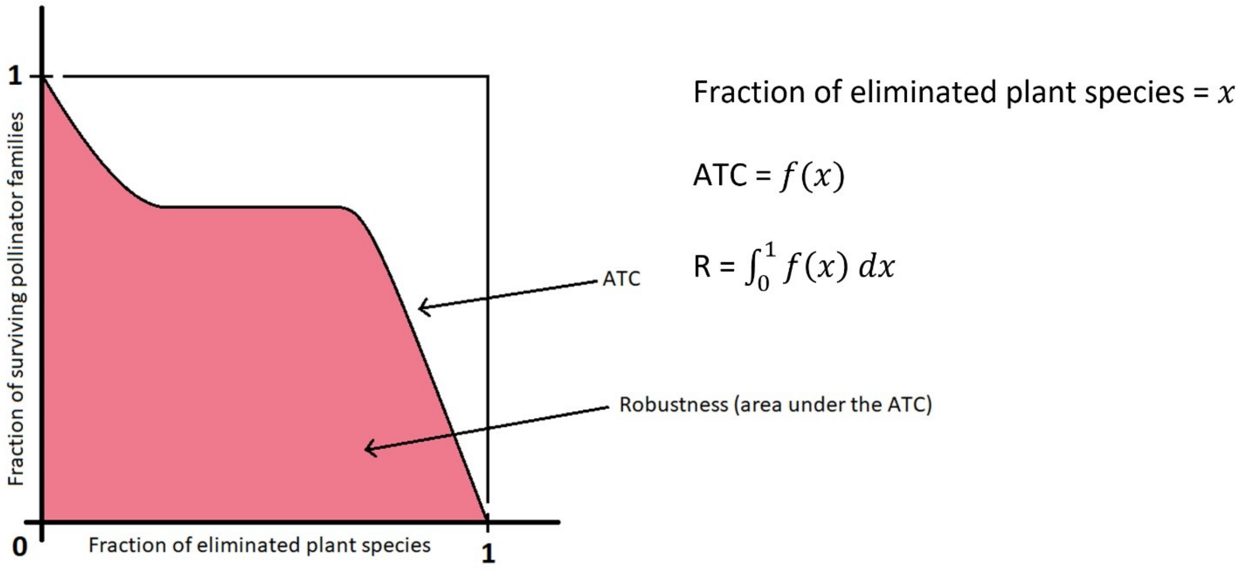

Figure S3. ATC. *An example of a possible Attack Tolerance Curve. Robustness is defined as the integral of the ATC in the range 0–1. Robustness tends to 1 in a perfectly nested system in which the extinction sequence goes in increasing order of link density (- → + scenario, least connected species are attacked first), while it tends to 0 in a perfectly nested system in which the extinction sequence goes in decreasing order of link density (+ → - scenario, most connected species are attacked first) (Burgos et al., 2007). Robustness takes on values between 0 and 1 in all situations in between these two extremes.*

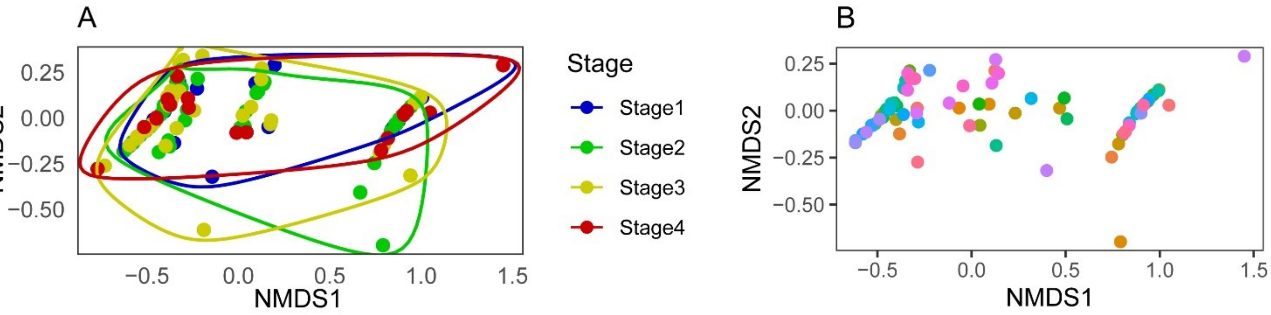

Figure S4. Plant species' roles within network motifs. A) *Non-metric Multidimensional Scaling (NMDS) ordination plot of plant species' contribution to node positions within network motifs, grouped by stage. Dots represent plant species involved in interactions.* B) *The same NMDS ordination but grouped by plant species. Dots represent plant species centroids.*

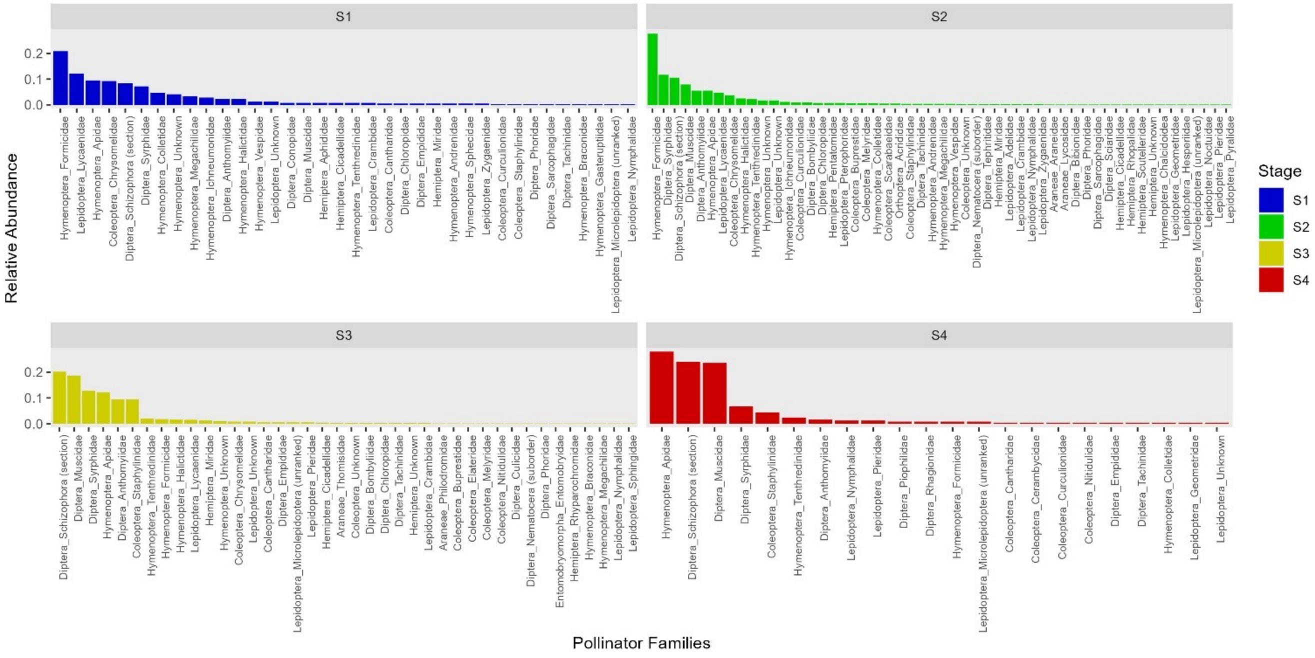

Figure S5. Pollinator families' relative abundances. *A list of pollinator families involved in plant–pollinator interactions within each stage, with relative abundance (n° of visits / n° tot of visits).*

### Supplementary tables

| Motifs' prevalence | | | | | | |
|---|---|---|---|---|---|---|
| **Most representative motifs** | | | | | | |
| | **Motif** | **Freq. Site A** | **Freq. Site B** | **Freq. Site C** | **Freq. Site D** | **Mean freq.** |
| **Stage 1** | 39 | 0.1549 | 0.1035 | 0.1094 | 0.0907 | 0.1146 |
| | 27 | 0.0717 | 0.1389 | 0.0958 | 0.0663 | 0.0932 |
| | 38 | 0.0793 | 0.0213 | 0.0475 | 0.1192 | 0.0668 |
| **Stage 2** | 19 | 0.0854 | 0.0781 | 0.1097 | 0.0860 | 0.0898 |
| | 25 | 0.0905 | 0.0574 | 0.1042 | 0.0983 | 0.0876 |
| | 27 | 0.1065 | 0.0716 | 0.0882 | 0.0628 | 0.0822 |
| **Stage 3** | 38 | 0.0831 | 0.1744 | 0.1181 | 0.0506 | 0.1065 |
| | 25 | 0.1144 | 0.0581 | 0.0946 | 0.0691 | 0.0841 |
| | 13 | 0.0363 | 0.1744 | 0.0542 | 0.0337 | 0.0747 |
| **Stage 4** | 17 | 0.3307 | 0.2941 | 0.0444 | 0.0072 | 0.1691 |
| | 7 | 0.2205 | 0.2941 | 0.0494 | 0.0075 | 0.1429 |
| | 44 | 0.3307 | 0.1765 | 0.0259 | 0.0050 | 0.1345 |

| Entries above threshold across all sites and stages | | | | |
|---|---|---|---|---|
| | **'r00_ind'** | **'r0_ind'** | **'c0_ind'** | **'vaznull'** |
| **n° entries with \|Z\|>1.96** | 460 | 451 | 438 | 131 |
| **% of entries with \|Z\|>1.96** | 74% | 75.92593% | 71.1039% | 22.16582% |
| **n° entries with Z>1.96** | 279 | 261 | 222 | 117 |
| **% of entries with Z>1.96** | 44.92754% | 43.93939% | 36.03896% | 19.79695% |
| **n° entries with Z<-1.96** | 181 | 190 | 216 | 14 |
| **% of entries with Z<-1.96** | 29.14654% | 31.98653% | 35.06494% | 2.368866% |
| **Most over-respresented motifs** | | | | |
| | **'r00_ind'** | **'r0_ind'** | **'c0_ind'** | **'vaznull'** |
| **Stage 1** | 1 (Z=+18.22) | 3 (Z=+19.08) | 2 (Z=+14.81) | 11 (Z=+1.96) |
| **Stage 2** | 4 (Z=+30.05) | 18 (Z=+79.25) | 2 (Z=+35.95) | 30 (Z=+2.93) |
| **Stage 3** | 1 (Z=+48.71) | 18 (Z=+72.62) | 2 (Z=+33.72) | 24 (Z=+2.19) |
| **Stage 4** | 25 (Z=+14.01) | 19 (Z=+16.53) | 7 (Z=+14.77) | 36 (Z=+5.55) |
| **Most under-respresented motifs** | | | | |
| | **'r00_ind'** | **'r0_ind'** | **'c0_ind'** | **'vaznull'** |
| **Stage 1** | 34 (Z=-4.98) | 17 (Z=-5.00) | 11 (Z=-3.92) | 38 (Z=-1.10) |
| **Stage 2** | 34 (Z=-12.31) | 40 (Z=-10.33) | 8 (Z=-8.20) | 38 (Z=-0.81) |
| **Stage 3** | 41 (Z=-9.72) | 40 (Z=-12.17) | 8 (Z=-7.82) | 29 (Z=-1.42) |
| **Stage 4** | 8 (Z=-62.01) | 8 (Z=-56.34) | 23 (Z=-4.46) | 26 (Z=-1.28) |

Table S1. Motifs' prevalence. *<u>Most representative motifs</u>: for each stage, the table shows the top 3 most frequent motifs within observed networks, with their respective normalised occurrences and mean across sites. <u>Entries above threshold across all sites and stages</u>: the table shows for each null model approach the number and the percentage of tested entries above threshold, over-represented entries and under-represented entries. <u>Most over-represented and under-represented motifs</u>: the table shows for each stage in each null model approach the motifs with highest and lowest mean Z-score across sites.*

| Plant species successional status | | | | | | | | |
|---|---|---|---|---|---|---|---|---|
| **Species** | **HS1** | **HS2** | **HS3** | **HS4** | **Label** | **HSp** | **$\bar{C}$** | **Ep** |
| *Achillea erba-rotta* | 0,375 | 0,875 | 0,750 | 0,250 | U | 0,1 | 12,56 | 0,008 |
| *Achillea millefolium* | 0,000 | 0,250 | 0,250 | 0,000 | M | 0,6 | 3,00 | 0,200 |
| *Adenostyles alpina* | 0,000 | 0,250 | 0,000 | 0,000 | M | 0,6 | 1,00 | 0,600 |
| *Ajuga pyramidalis* | 0,000 | 0,250 | 0,125 | 0,000 | M | 0,6 | 1,00 | 0,600 |
| *Alchemilla alpina* | 0,250 | 0,000 | 0,000 | 0,000 | E | 0,8 | 1,00 | 0,800 |
| *Alchemilla monticola* | 0,000 | 0,750 | 0,500 | 0,000 | M | 0,6 | 1,00 | 0,600 |
| *Antennaria dioica* | 0,125 | 0,375 | 0,250 | 0,000 | M | 0,6 | 1,20 | 0,500 |
| *Anthoxanthum odoratum* | 0,375 | 0,625 | 0,625 | 0,125 | EU | 0,5 | 8,67 | 0,058 |
| *Anthriscus sylvestris* | 0,000 | 0,000 | 0,250 | 0,000 | M | 0,6 | 1,00 | 0,600 |
| *Anthyllis vulneraria* | 1,000 | 1,000 | 0,625 | 0,125 | EU | 0,5 | 10,70 | 0,047 |
| *Bartsia alpina* | 0,125 | 0,375 | 0,000 | 0,000 | M | 0,6 | 3,00 | 0,200 |
| *Biscutella laevigata* | 0,000 | 0,125 | 0,250 | 0,000 | M | 0,6 | 11,00 | 0,055 |
| *Campanula barbata* | 0,500 | 0,500 | 0,750 | 0,000 | EU | 0,5 | 5,00 | 0,100 |
| *Campanula cochleariifolia* | 0,500 | 0,000 | 0,000 | 0,000 | E | 0,8 | 1,00 | 0,800 |
| *Campanula rhomboidalis* | 0,000 | 0,000 | 0,000 | 0,250 | L | 0,4 | 1,00 | 0,400 |
| *Campanula scheuchzeri* | 0,125 | 0,625 | 0,375 | 0,000 | M | 0,6 | 3,00 | 0,200 |
| *Cardamine resedifolia* | 0,125 | 0,000 | 0,000 | 0,125 | R | 0,9 | 1,00 | 0,900 |
| *Carduus defloratus* | 0,000 | 0,750 | 0,250 | 0,000 | M | 0,6 | 12,50 | 0,048 |
| *Centaurea nervosa* | 0,000 | 0,250 | 0,250 | 0,000 | M | 0,6 | 15,00 | 0,040 |
| *Cerastium alpinum* | 0,500 | 0,000 | 0,000 | 0,000 | E | 0,8 | 1,00 | 0,800 |
| *Cerastium arvense* | 0,125 | 0,750 | 0,500 | 0,000 | M | 0,6 | 5,40 | 0,111 |
| *Cerastium latifolium* | 0,375 | 0,375 | 0,000 | 0,000 | EM | 0,7 | 3,20 | 0,219 |
| *Chaerophyllum villarsii* | 0,000 | 0,500 | 0,000 | 0,000 | M | 0,6 | 1,00 | 0,600 |
| *Chenopodium bonus-henricus* | 0,000 | 0,000 | 0,250 | 0,000 | M | 0,6 | 1,00 | 0,600 |
| *Crepis pontana* | 0,000 | 0,000 | 0,250 | 0,000 | M | 0,6 | 1,00 | 0,600 |
| *Dactylorhiza maculata* | 0,000 | 0,000 | 0,000 | 0,250 | L | 0,4 | 1,00 | 0,400 |
| *Dactylorhiza majalis* | 0,000 | 0,250 | 0,000 | 0,000 | M | 0,6 | 1,00 | 0,600 |
| *Dryas octopetala* | 0,000 | 0,125 | 0,625 | 0,000 | M | 0,6 | 4,33 | 0,138 |
| *Epilobium anagallidifolium* | 0,250 | 0,000 | 0,000 | 0,000 | E | 0,8 | 1,00 | 0,800 |
| *Epilobium angustifolium* | 0,000 | 0,500 | 0,375 | 0,125 | M | 0,6 | 5,00 | 0,120 |
| *Epilobium fleischeri* | 0,625 | 0,750 | 0,250 | 0,000 | EU | 0,5 | 18,71 | 0,027 |
| *Erigeron alpinus* | 0,125 | 0,500 | 0,250 | 0,000 | M | 0,6 | 1,00 | 0,600 |

| | | | | | | | | |
|---|---|---|---|---|---|---|---|---|
| *Euphrasia minima* | 0,500 | 0,250 | 0,125 | 0,000 | EM | 0,7 | 1,67 | 0,420 |
| *Galium anisophyllon* | 0,000 | 0,750 | 0,250 | 0,000 | M | 0,6 | 3,50 | 0,171 |
| *Gentiana nivalis* | 0,000 | 0,750 | 0,375 | 0,000 | M | 0,6 | 2,50 | 0,240 |
| *Geum montanum* | 0,000 | 0,250 | 0,000 | 0,000 | M | 0,6 | 1,00 | 0,600 |
| *Gymnadenia conopsea* | 0,000 | 0,750 | 0,000 | 0,000 | M | 0,6 | 2,00 | 0,300 |
| *Gypsofila repens* | 0,500 | 0,000 | 0,250 | 0,000 | EM | 0,7 | 1,00 | 0,700 |
| *Hieracium angustifolium* | 0,250 | 0,250 | 0,000 | 0,000 | EM | 0,7 | 1,00 | 0,700 |
| *Hieracium murorum* | 0,125 | 0,375 | 0,250 | 0,500 | LU | 0,2 | 13,00 | 0,015 |
| *Hieracium staticifolium* | 1,000 | 0,625 | 0,250 | 0,000 | EU | 0,5 | 10,43 | 0,048 |
| *Hippocrepis comosa* | 0,000 | 0,250 | 0,250 | 0,000 | M | 0,6 | 2,00 | 0,300 |
| *Homogyne alpina* | 0,000 | 0,125 | 0,375 | 0,000 | M | 0,6 | 1,33 | 0,450 |
| *Huguenina tanacetifolia* | 0,000 | 0,000 | 0,500 | 0,000 | M | 0,6 | 1,00 | 0,600 |
| *Leontodon helveticus* | 0,000 | 0,500 | 0,250 | 0,000 | M | 0,6 | 5,57 | 0,108 |
| *Leontodon hispidus* | 0,500 | 0,875 | 0,875 | 0,125 | EU | 0,5 | 9,86 | 0,051 |
| *Leucanthemopsis alpina* | 0,875 | 0,500 | 0,000 | 0,125 | EM | 0,7 | 1,83 | 0,382 |
| *Leucanthemum adustum* | 0,250 | 0,000 | 0,000 | 0,000 | E | 0,8 | 1,00 | 0,800 |
| *Leucanthemum vulgare* | 0,250 | 0,375 | 0,125 | 0,000 | EM | 0,7 | 3,00 | 0,233 |
| *Linaria alpina* | 0,500 | 0,000 | 0,000 | 0,000 | E | 0,8 | 1,00 | 0,800 |
| *Lotus corniculatus* | 1,000 | 1,000 | 1,000 | 0,625 | U | 0,1 | 11,00 | 0,009 |
| *Melampyrum sylvaticum* | 0,000 | 0,500 | 0,000 | 0,500 | ML | 0,3 | 1,00 | 0,300 |
| *Minuartia verna* | 0,375 | 0,000 | 0,000 | 0,000 | E | 0,8 | 1,00 | 0,800 |
| *Myosotis alpestris* | 0,250 | 0,625 | 0,750 | 0,375 | U | 0,1 | 1,86 | 0,054 |
| *Myosotis arvensis* | 0,000 | 0,250 | 0,000 | 0,250 | ML | 0,3 | 1,00 | 0,300 |
| *Nigritella rhellicani* | 0,000 | 0,125 | 0,250 | 0,000 | M | 0,6 | 1,00 | 0,600 |
| *Orchis mascula* | 0,250 | 0,250 | 0,000 | 0,000 | EM | 0,7 | 1,00 | 0,700 |
| *Oxyria digyna* | 0,250 | 0,000 | 0,000 | 0,000 | E | 0,8 | 1,00 | 0,800 |
| *Parnassia palustre* | 0,000 | 0,250 | 0,000 | 0,000 | M | 0,6 | 1,00 | 0,600 |
| *Pedicularis kerneri* | 0,250 | 0,000 | 0,000 | 0,000 | E | 0,8 | 1,00 | 0,800 |
| *Pedicularis tuberosa* | 0,250 | 0,000 | 0,000 | 0,000 | E | 0,8 | 1,00 | 0,800 |
| *Peucedanum ostruthium* | 0,000 | 0,375 | 0,375 | 0,500 | LU | 0,2 | 4,50 | 0,044 |
| *Phyteuma betonicifolium* | 0,000 | 0,500 | 0,875 | 0,000 | M | 0,6 | 10,50 | 0,057 |
| *Phyteuma hemisphaericum* | 0,250 | 0,125 | 0,250 | 0,000 | EM | 0,7 | 1,00 | 0,700 |
| *Pilosella cymosa* | 0,500 | 0,250 | 0,000 | 0,000 | EM | 0,7 | 1,00 | 0,700 |
| *Pilosella officinarum* | 0,500 | 0,750 | 0,500 | 0,000 | EU | 0,5 | 9,00 | 0,056 |
| *Polygala amarella* | 0,000 | 0,250 | 0,000 | 0,000 | M | 0,6 | 1,00 | 0,600 |
| *Polygonum viviparum* | 0,000 | 0,500 | 0,125 | 0,000 | M | 0,6 | 3,00 | 0,200 |
| *Potentilla aurea* | 0,125 | 0,750 | 0,625 | 0,125 | M | 0,6 | 15,00 | 0,040 |
| *Primula farinosa* | 0,000 | 0,250 | 0,000 | 0,000 | M | 0,6 | 1,00 | 0,600 |
| *Pyrola minor* | 0,250 | 0,375 | 0,250 | 0,000 | EU | 0,5 | 1,67 | 0,300 |
| *Ranunculus acris* | 0,000 | 0,125 | 0,250 | 0,000 | M | 0,6 | 3,00 | 0,200 |
| *Ranunculus montanus* | 0,000 | 0,875 | 0,750 | 0,125 | M | 0,6 | 13,33 | 0,045 |
| *Ranunculus platanifolius* | 0,000 | 0,000 | 0,000 | 0,250 | L | 0,4 | 1,00 | 0,400 |
| *Ranunculus villarsii* | 0,000 | 1,000 | 0,500 | 0,250 | LU | 0,2 | 13,33 | 0,015 |
| *Rhinanthus major* | 0,000 | 0,250 | 0,000 | 0,000 | M | 0,6 | 1,00 | 0,600 |
| *Rhinanthus minor* | 0,000 | 0,500 | 0,000 | 0,000 | M | 0,6 | 1,00 | 0,600 |

| | | | | | | | | |
|---|---|---|---|---|---|---|---|---|
| *Rhododendron ferrugineum* | 0,000 | 1,000 | 0,750 | 1,000 | LU | 0,2 | 31,55 | 0,006 |
| *Rumex acetosa* | 0,000 | 0,250 | 0,250 | 0,000 | M | 0,6 | 1,00 | 0,600 |
| *Rumex acetosella* | 0,000 | 0,250 | 0,000 | 0,000 | M | 0,6 | 1,00 | 0,600 |
| *Rumex alpinus* | 0,000 | 0,000 | 0,250 | 0,125 | M | 0,6 | 8,00 | 0,075 |
| *Rumex scutatus* | 0,375 | 0,500 | 0,000 | 0,125 | EM | 0,7 | 11,50 | 0,061 |
| *Saxifraga aizoides* | 0,125 | 0,375 | 0,000 | 0,000 | M | 0,6 | 1,00 | 0,600 |
| *Saxifraga bryoides* | 0,625 | 0,250 | 0,000 | 0,000 | EM | 0,7 | 3,75 | 0,187 |
| *Saxifraga paniculata* | 0,750 | 0,750 | 0,000 | 0,000 | EM | 0,7 | 3,67 | 0,191 |
| *Sempervivum arachnoideum* | 0,250 | 0,125 | 0,125 | 0,000 | E | 0,8 | 1,00 | 0,800 |
| *Sempervivum montanum* | 0,375 | 0,875 | 0,125 | 0,375 | U | 0,1 | 1,00 | 0,100 |
| *Silene acaulis* | 0,375 | 0,000 | 0,000 | 0,000 | E | 0,8 | 1,00 | 0,800 |
| *Silene dioica* | 0,000 | 0,000 | 0,250 | 0,000 | M | 0,6 | 1,00 | 0,600 |
| *Silene rupestris* | 0,375 | 0,375 | 0,250 | 0,125 | EU | 0,5 | 2,00 | 0,250 |
| *Silene vulgaris* | 0,125 | 0,625 | 0,625 | 0,125 | M | 0,6 | 7,00 | 0,086 |
| *Solidago virgaurea* | 0,125 | 0,250 | 0,000 | 0,125 | M | 0,6 | 1,00 | 0,600 |
| *Taraxacum officinale* | 0,250 | 0,000 | 0,000 | 0,000 | E | 0,8 | 3,50 | 0,229 |
| *Telephium imperati* | 0,000 | 0,000 | 0,250 | 0,000 | M | 0,6 | 1,00 | 0,600 |
| *Thesium alpinum* | 0,000 | 0,125 | 0,375 | 0,250 | ML | 0,3 | 1,50 | 0,200 |
| *Thymus praecox* | 0,500 | 0,500 | 0,250 | 0,000 | EU | 0,5 | 5,00 | 0,100 |
| *Tofieldia calyculata* | 0,000 | 0,250 | 0,000 | 0,000 | M | 0,6 | 1,00 | 0,600 |
| *Trifolium badium* | 0,000 | 0,750 | 0,250 | 0,000 | M | 0,6 | 1,00 | 0,600 |
| *Trifolium hybridum* | 0,000 | 0,250 | 0,250 | 0,000 | M | 0,6 | 1,00 | 0,600 |
| *Trifolium ochroleucon* | 0,250 | 0,500 | 0,750 | 0,000 | EU | 0,5 | 1,00 | 0,500 |
| *Trifolium pallescens* | 1,000 | 0,750 | 0,375 | 0,125 | EU | 0,5 | 9,71 | 0,051 |
| *Trifolium pratense* | 0,000 | 0,875 | 0,500 | 0,250 | LU | 0,2 | 6,50 | 0,031 |
| *Trifolium repens* | 0,000 | 0,000 | 0,500 | 0,000 | M | 0,6 | 1,00 | 0,600 |
| *Vaccinium vitis-idaea* | 0,000 | 0,125 | 0,625 | 0,875 | ML | 0,3 | 15,33 | 0,020 |
| *Valeriana tripteris* | 0,000 | 0,000 | 0,000 | 0,250 | L | 0,4 | 16,00 | 0,025 |
| *Veronica chamaedrys* | 0,000 | 0,000 | 0,375 | 0,125 | M | 0,6 | 2,00 | 0,300 |
| *Veronica fruticans* | 0,000 | 0,625 | 0,500 | 0,000 | M | 0,6 | 2,75 | 0,218 |
| *Vicia sepium* | 0,000 | 0,000 | 0,000 | 0,250 | L | 0,4 | 1,00 | 0,400 |

**Rules for labels and probabilities assignment**

HS1 < 0.25 & HS2 < 0.25 & HS3 < 0.25 & HS4 < 0.25 → **Rare (R)** → HSp=0.9
HS1 >= 0.25 & HS2 < 0.25 & HS3 < 0.25 & HS4 < 0.25 → **Early-successional (E)** → HSp=0.8
(HS1 >= 0.25 & HS2 >= 0.25 & HS3 < 0.25 & HS4 < 0.25) |
(HS1 >= 0.25 & HS3 >= 0.25 & HS2 < 0.25 & HS4 < 0.25) → **Early to mid-successional (EM)** → HSp=0.7
(HS2 >= 0.25 & HS1 < 0.25 & HS3 < 0.25 & HS4 < 0.25) |
(HS3 >= 0.25 & HS1 < 0.25 & HS2 < 0.25 & HS4 < 0.25) |
(HS2 >= 0.25 & HS3 >= 0.25 & HS1 < 0.25 & HS4 < 0.25) → **Mid-successional (M)** → HSp=0.6
HS1 >= 0.25 & HS2 >= 0.25 & HS3 >= 0.25 & HS4 < 0.25 → **Early-ubiquitous (EU)** → HSp=0.5
HS4 >= 0.25 & HS1 < 0.25 & HS2 < 0.25 & HS3 < 0.25 → **Late-successional (L)** → HSp=0.4
(HS2 >= 0.25 & HS4 >= 0.25 & HS1 < 0.25 & HS3 < 0.25) |
(HS3 >= 0.25 & HS4 >= 0.25 & HS1 < 0.25 & HS2 < 0.25) → **Mid to late-successional (ML)** → HSp=0.3
HS2 >= 0.25 & HS3 >= 0.25 & HS4 >= 0.25 & HS1 < 0.25 → **Late-ubiquitous (LU)** → HSp=0.2
(HS1 >= 0.25 & HS2 >= 0.25 & HS3 >= 0.25 & HS4 >= 0.25) |
(HS1 >= 0.25 & HS2 >= 0.25 & HS4 >= 0.25 & HS3 < 0.25) |
(HS1 >= 0.25 & HS3 >= 0.25 & HS4 >= 0.25 & HS2 < 0.25) → **Ubiquitous (U)** → HSp=0.1

Table S2. Plant species successional status. *A list of surveyed blooming plant species with corresponding successional status, mean coverage and extinction probability. HS stands for 'Habitat Specificity'; HS1, HS2, HS3 and HS4 are habitat specificity values of Stage 1, Stage 2, Stage 3, and Stage 4. Each HS value is the proportion of occupied sites within the corresponding stage, calculated as the average between our data and data collected by Anonymous and Anonymous in 2022 in the same sites. 'Label' is the successional status, identified according to the rules set out in the final section of the table. 'HSp' stands for 'Habitat Specificity Probability', which is an assigned probability belonging to each specific successional label. '$\bar{C}$' is the mean coverage calculated on data collected by Anonymous and Anonymous in 2022 in the same sites. 'Ep' stands for 'Extinction Probability' and is calculated as $E_p = {HS_p}/{\bar{C}}$. The higher the extinction probability, the higher the chance that the species is extracted during the extinction cascades. Extinction probability values are not indicative of the actual absolute likelihood of species extinction in the future; they only serve as relative values to determine the extinction sequence during the robustness calculation.*

| Network robusteness | | | | |
|---|---|---|---|---|
| **Site** | **Test model** | **Null model** | **Z-score** | **p (Wilcox)** |
| 1A | 0.6371357 | 0.6277974 | 3.942493 | 2.324963e-03 |
| 1B | 0.6272041 | 0.5922027 | 19.735111 | < 2.2e-16 |
| 1C | 0.7172250 | 0.6399113 | 33.179814 | < 2.2e-16 |
| 1D | 0.6483238 | 0.6270074 | 7.284407 | 2.402826e-12 |
| 2A | 0.7499226 | 0.6468331 | 37.398032 | < 2.2e-16 |
| 2B | 0.7473954 | 0.6766542 | 32.103407 | < 2.2e-16 |
| 2C | 0.7214505 | 0.6096866 | 48.016415 | < 2.2e-16 |
| 2D | 0.7780932 | 0.6868170 | 29.029148 | < 2.2e-16 |
| 3A | 0.7076826 | 0.6568768 | 15.846525 | < 2.2e-16 |
| 3B | 0.7360568 | 0.5780000 | 23.523647 | < 2.2e-16 |
| 3C | 0.6285985 | 0.6491748 | -5.866536 | 2.883235e-07 |
| 3D | 0.7697943 | 0.6989495 | 24.801532 | < 2.2e-16 |
| 4A | 0.5000000 | 0.5000000 | ---- | ---- |
| 4B | 0.5000000 | 0.5000000 | ---- | ---- |
| 4C | 0.5329652 | 0.5728809 | -10.883420 | < 2.2e-16 |
| 4D | 0.7754336 | 0.6281913 | 48.704895 | < 2.2e-16 |

Table S3. Network robustness. *The table shows robustness values for each site in both the test model and the null model, the corresponding Z-scores and the associated Mann-Whitney/Wilcox test p-values. Each robustness value is the average of 1000 iterations of the extinction cascade.*

| Plant species' roles | | | | | |
|---|---|---|---|---|---|
| Entries above threshold | | | | | |
| **Overall** | | **'r00_ind'** | **'r0_ind'** | **'c0_ind'** | **'vaznull'** |
| | **n° entries with \|Z\|>1.96** | 2159 | 1460 | 1742 | 412 |
| | **% of entries with \|Z\|>1.96** | 85.26856% | 65.7954% | 67.70307% | 14.6933% |
| | **n° entries with Z>1.96** | 461 | 612 | 312 | 302 |
| | **% of entries with Z>1.96** | 18.20695% | 27.57999% | 12.12592% | 10.77033% |
| | **n° entries with Z<-1.96** | 1698 | 848 | 1430 | 110 |
| | **% of entries with Z<-1.96** | 67.06161% | 38.21541% | 55.57715% | 3.922967% |
| **Stage 1** | | **'r00_ind'** | **'r0_ind'** | **'c0_ind'** | **'vaznull'** |
| | **n° entries with \|Z\|>1.96** | 437 | 309 | 384 | 68 |
| | **% of entries with \|Z\|>1.96** | 84.68992% | 71.36259% | 73.56322% | 12.01413% |
| | **n° entries with Z>1.96** | 97 | 116 | 63 | 47 |
| | **% of entries with Z>1.96** | 18.79845% | 26.78984% | 12.06897% | 8.303887% |
| | **n° entries with Z<-1.96** | 340 | 193 | 321 | 21 |
| | **% of entries with Z<-1.96** | 65.89147% | 44.57275% | 61.49425% | 3.710247% |
| **Stage 2** | | **'r00_ind'** | **'r0_ind'** | **'c0_ind'** | **'vaznull'** |
| | **n° entries with \|Z\|>1.96** | 819 | 593 | 637 | 175 |
| | **% of entries with \|Z\|>1.96** | 86.11987% | 70.01181% | 66.28512% | 16.68255% |
| | **n° entries with Z>1.96** | 184 | 237 | 123 | 128 |
| | **% of entries with Z>1.96** | 19.34805% | 27.98111% | 12.79917% | 12.2021% |
| | **n° entries with Z<-1.96** | 635 | 356 | 514 | 47 |
| | **% of entries with Z<-1.96** | 66.77182% | 42.0307% | 53.48595% | 4.480458% |
| **Stage 3** | | **'r00_ind'** | **'r0_ind'** | **'c0_ind'** | **'vaznull'** |
| | **n° entries with \|Z\|>1.96** | 637 | 438 | 495 | 134 |
| | **% of entries with \|Z\|>1.96** | 88.22715% | 66.87023% | 68.46473% | 16.68742% |
| | **n° entries with Z>1.96** | 150 | 195 | 105 | 106 |
| | **% of entries with Z>1.96** | 20.77562% | 29.77099% | 14.52282% | 13.2005% |
| | **n° entries with Z<-1.96** | 487 | 243 | 390 | 28 |
| | **% of entries with Z<-1.96** | 67.45152% | 37.09924% | 53.94191% | 3.486924% |

| Stage 4 | | 'r00_ind' | 'r0_ind' | 'c0_ind' | 'vaznull' |
|---|---|---|---|---|---|
| | n° entries with \|Z\|>1.96 | 266 | 120 | 226 | 35 |
| | % of entries with \|Z\|>1.96 | 77.55102% | 42.25352% | 61.58038% | 9.067358% |
| | n° entries with Z>1.96 | 30 | 64 | 21 | 21 |
| | % of entries with Z>1.96 | 8.746356% | 22.53521% | 5.722071% | 5.440415% |
| | n° entries with Z<-1.96 | 236 | 56 | 205 | 14 |
| | % of entries with Z<-1.96 | 68.80466% | 19.71831% | 55.85831% | 3.626943% |

| Most over-respresented node positions | | | | |
|---|---|---|---|---|
| | 'r00_ind' | 'r0_ind' | 'c0_ind' | 'vaznull' |
| Stage 1 | 18 (Z=-2.56) | 6 (Z=-0.08) | 18 (Z=-1.31) | 41 (Z=+0.73) |
| Stage 2 | 6 (Z=-2.12) | 34 (Z=+1.59) | 6 (Z=-0.90) | 34 (Z=+1.67) |
| Stage 3 | 6 (Z=-1.64) | 34 (Z=+2.72) | 6 (Z=-0.72) | 34 (Z=+2.24) |
| Stage 4 | 18 (Z=-1.54) | 6 (Z=+0.67) | 25 (Z=-0.39) | 34 (Z=+0.89) |
| **Most under-respresented node positions** | | | | |
| | 'r00_ind' | 'r0_ind' | 'c0_ind' | 'vaznull' |
| Stage 1 | 35 (Z=-17.45) | 24 (Z=-1.71) | 35 (Z=-7.74) | 25 (Z=-0.21) |
| Stage 2 | 35 (Z=-18.15) | 28 (Z=-1.11) | 35 (Z=-6.93) | 25 (Z=-0.08) |
| Stage 3 | 35 (Z=-15-65) | 24 (Z=+0.85) | 44 (Z=-5.07) | 25 (Z=-0.15) |
| Stage 4 | 35 (Z=-13.61) | 18 (Z=+0.50) | 41 (Z=-3.98) | 22 (Z=-0.36) |

Table S4. Plant species' roles. *Entries above threshold*: *the table shows for each null model approach the number and the percentage of tested entries above threshold, over-represented entries and under-represented entries (overall and for each stage). Most over-represented and under-represented node positions: the table shows for each stage in each null model approach the node positions with highest and lowest mean Z-score across plant species. Some Z-scores are negative in the over-represented section and positive in the under-represented one, but this is due to the fact that the value is the average across species (some species may have a positive and significant Z-score, others a negative one).*

| Pollinator families' abundance | | | |
|---|---|---|---|
| **Stage** | **Family** | **Abundance** | **Relative Abundance** |
| S1 | Hymenoptera_Formicidae | 81 | 0,20822622 |
| S1 | Lepidoptera_Lycaenidae | 47 | 0,12082262 |
| S1 | Hymenoptera_Apidae | 37 | 0,09511568 |
| S1 | Coleoptera_Chrysomelidae | 36 | 0,09254499 |
| S1 | Diptera_Schizophora (section) | 33 | 0,0848329 |
| S1 | Diptera_Syrphidae | 28 | 0,07197943 |
| S1 | Hymenoptera_Colletidae | 18 | 0,04627249 |
| S1 | Hymenoptera_Unknown | 16 | 0,04113111 |
| S1 | Hymenoptera_Megachilidae | 13 | 0,03341902 |
| S1 | Hymenoptera_Ichneumonidae | 11 | 0,02827763 |
| S1 | Diptera_Anthomyiidae | 9 | 0,02313625 |
| S1 | Hymenoptera_Halictidae | 9 | 0,02313625 |
| S1 | Lepidoptera_Unknown | 5 | 0,01285347 |
| S1 | Hymenoptera_Vespidae | 5 | 0,01285347 |
| S1 | Lepidoptera_Crambidae | 3 | 0,00771208 |
| S1 | Hemiptera_Aphididae | 3 | 0,00771208 |
| S1 | Hemiptera_Cicadellidae | 3 | 0,00771208 |
| S1 | Hymenoptera_Tenthredinidae | 3 | 0,00771208 |
| S1 | Diptera_Conopidae | 3 | 0,00771208 |
| S1 | Diptera_Muscidae | 3 | 0,00771208 |
| S1 | Hemiptera_Miridae | 2 | 0,00514139 |
| S1 | Diptera_Empididae | 2 | 0,00514139 |
| S1 | Diptera_Chloropidae | 2 | 0,00514139 |
| S1 | Hymenoptera_Andrenidae | 2 | 0,00514139 |
| S1 | Coleoptera_Cantharidae | 2 | 0,00514139 |
| S1 | Lepidoptera_Zygaenidae | 2 | 0,00514139 |
| S1 | Hymenoptera_Sphecidae | 2 | 0,00514139 |
| S1 | Hymenoptera_Braconidae | 1 | 0,00257069 |
| S1 | Diptera_Tachinidae | 1 | 0,00257069 |
| S1 | Coleoptera_Curculionidae | 1 | 0,00257069 |
| S1 | Diptera_Phoridae | 1 | 0,00257069 |
| S1 | Coleoptera_Staphylinidae | 1 | 0,00257069 |
| S1 | Hymenoptera_Gasteruptiidae | 1 | 0,00257069 |
| S1 | Lepidoptera_Microlepidoptera (unranked) | 1 | 0,00257069 |
| S1 | Diptera_Sarcophagidae | 1 | 0,00257069 |
| S1 | Lepidoptera_Nymphalidae | 1 | 0,00257069 |
| S2 | Hymenoptera_Formicidae | 220 | 0,27638191 |
| S2 | Diptera_Syrphidae | 93 | 0,11683417 |
| S2 | Diptera_Schizophora (section) | 84 | 0,10552764 |
| S2 | Diptera_Muscidae | 63 | 0,07914573 |
| S2 | Diptera_Anthomyiidae | 44 | 0,05527638 |
| S2 | Hymenoptera_Apidae | 44 | 0,05527638 |
| S2 | Lepidoptera_Lycaenidae | 38 | 0,04773869 |

| | | | |
|---|---|---|---|
| S2 | Coleoptera_Chrysomelidae | 30 | 0,03768844 |
| S2 | Hymenoptera_Halictidae | 20 | 0,02512563 |
| S2 | Hymenoptera_Tenthredinidae | 18 | 0,02261307 |
| S2 | Lepidoptera_Unknown | 14 | 0,01758794 |
| S2 | Hymenoptera_Unknown | 14 | 0,01758794 |
| S2 | Hymenoptera_Ichneumonidae | 9 | 0,01130653 |
| S2 | Coleoptera_Curculionidae | 8 | 0,01005025 |
| S2 | Diptera_Bombyliidae | 8 | 0,01005025 |
| S2 | Diptera_Chloropidae | 6 | 0,00753769 |
| S2 | Hemiptera_Pentatomidae | 6 | 0,00753769 |
| S2 | Lepidoptera_Pterophoridae | 6 | 0,00753769 |
| S2 | Coleoptera_Buprestidae | 5 | 0,00628141 |
| S2 | Coleoptera_Melyridae | 5 | 0,00628141 |
| S2 | Hymenoptera_Colletidae | 5 | 0,00628141 |
| S2 | Coleoptera_Scarabaeidae | 4 | 0,00502513 |
| S2 | Orthoptera_Acrididae | 4 | 0,00502513 |
| S2 | Hymenoptera_Megachilidae | 3 | 0,00376884 |
| S2 | Diptera_Tachinidae | 3 | 0,00376884 |
| S2 | Coleoptera_Staphylinidae | 3 | 0,00376884 |
| S2 | Hymenoptera_Vespidae | 3 | 0,00376884 |
| S2 | Hymenoptera_Andrenidae | 3 | 0,00376884 |
| S2 | Lepidoptera_Adelidae | 2 | 0,00251256 |
| S2 | Coleoptera_Unknown | 2 | 0,00251256 |
| S2 | Hemiptera_Miridae | 2 | 0,00251256 |
| S2 | Diptera_Nematocera (suborder) | 2 | 0,00251256 |
| S2 | Lepidoptera_Crambidae | 2 | 0,00251256 |
| S2 | Diptera_Tephritidae | 2 | 0,00251256 |
| S2 | Lepidoptera_Zygaenidae | 2 | 0,00251256 |
| S2 | Lepidoptera_Nymphalidae | 2 | 0,00251256 |
| S2 | Araneae_Lycosidae | 1 | 0,00125628 |
| S2 | Lepidoptera_Microlepidoptera (unranked) | 1 | 0,00125628 |
| S2 | Diptera_Bibionidae | 1 | 0,00125628 |
| S2 | Lepidoptera_Geometridae | 1 | 0,00125628 |
| S2 | Diptera_Phoridae | 1 | 0,00125628 |
| S2 | Diptera_Sciaridae | 1 | 0,00125628 |
| S2 | Lepidoptera_Pieridae | 1 | 0,00125628 |
| S2 | Hemiptera_Cicadellidae | 1 | 0,00125628 |
| S2 | Hemiptera_Unknown | 1 | 0,00125628 |
| S2 | Hemiptera_Scutelleridae | 1 | 0,00125628 |
| S2 | Hemiptera_Rhopalidae | 1 | 0,00125628 |
| S2 | Hymenoptera_Chalcidoidea | 1 | 0,00125628 |
| S2 | Lepidoptera_Noctuidae | 1 | 0,00125628 |
| S2 | Diptera_Sarcophagidae | 1 | 0,00125628 |
| S2 | Lepidoptera_Pyralidae | 1 | 0,00125628 |
| S2 | Araneae_Araneidae | 1 | 0,00125628 |

| | | | |
|---|---|---|---|
| S2 | Lepidoptera_Hesperiidae | 1 | 0,00125628 |
| S3 | Diptera_Schizophora (section) | 150 | 0,20188425 |
| S3 | Diptera_Muscidae | 139 | 0,18707941 |
| S3 | Diptera_Syrphidae | 95 | 0,12786003 |
| S3 | Hymenoptera_Apidae | 91 | 0,12247645 |
| S3 | Diptera_Anthomyiidae | 71 | 0,09555855 |
| S3 | Coleoptera_Staphylinidae | 70 | 0,09421265 |
| S3 | Hymenoptera_Tenthredinidae | 15 | 0,02018843 |
| S3 | Hymenoptera_Formicidae | 13 | 0,01749664 |
| S3 | Hymenoptera_Halictidae | 12 | 0,01615074 |
| S3 | Lepidoptera_Lycaenidae | 11 | 0,01480485 |
| S3 | Hemiptera_Miridae | 9 | 0,01211306 |
| S3 | Hymenoptera_Unknown | 8 | 0,01076716 |
| S3 | Coleoptera_Chrysomelidae | 6 | 0,00807537 |
| S3 | Lepidoptera_Unknown | 6 | 0,00807537 |
| S3 | Lepidoptera_Pieridae | 4 | 0,00538358 |
| S3 | Lepidoptera_Microlepidoptera (unranked) | 4 | 0,00538358 |
| S3 | Diptera_Empididae | 4 | 0,00538358 |
| S3 | Coleoptera_Cantharidae | 4 | 0,00538358 |
| S3 | Hemiptera_Cicadellidae | 3 | 0,00403769 |
| S3 | Diptera_Chloropidae | 2 | 0,00269179 |
| S3 | Lepidoptera_Crambidae | 2 | 0,00269179 |
| S3 | Diptera_Tachinidae | 2 | 0,00269179 |
| S3 | Hemiptera_Unknown | 2 | 0,00269179 |
| S3 | Araneae_Thomisidae | 2 | 0,00269179 |
| S3 | Coleoptera_Unknown | 2 | 0,00269179 |
| S3 | Diptera_Bombyliidae | 2 | 0,00269179 |
| S3 | Hymenoptera_Braconidae | 1 | 0,0013459 |
| S3 | Coleoptera_Elateridae | 1 | 0,0013459 |
| S3 | Entomobryomorpha_Entomobryidae | 1 | 0,0013459 |
| S3 | Hemiptera_Rhyparochromidae | 1 | 0,0013459 |
| S3 | Lepidoptera_Sphingidae | 1 | 0,0013459 |
| S3 | Coleoptera_Nitidulidae | 1 | 0,0013459 |
| S3 | Araneae_Philodromidae | 1 | 0,0013459 |
| S3 | Diptera_Culicidae | 1 | 0,0013459 |
| S3 | Diptera_Nematocera (suborder) | 1 | 0,0013459 |
| S3 | Coleoptera_Buprestidae | 1 | 0,0013459 |
| S3 | Coleoptera_Melyridae | 1 | 0,0013459 |
| S3 | Diptera_Phoridae | 1 | 0,0013459 |
| S3 | Lepidoptera_Nymphalidae | 1 | 0,0013459 |
| S3 | Hymenoptera_Megachilidae | 1 | 0,0013459 |
| S4 | Hymenoptera_Apidae | 70 | 0,28 |
| S4 | Diptera_Schizophora (section) | 60 | 0,24 |
| S4 | Diptera_Muscidae | 59 | 0,236 |
| S4 | Diptera_Syrphidae | 17 | 0,068 |

| | | | |
|---|---|---|---|
| S4 | Coleoptera_Staphylinidae | 11 | 0,044 |
| S4 | Hymenoptera_Tenthredinidae | 6 | 0,024 |
| S4 | Diptera_Anthomyiidae | 4 | 0,016 |
| S4 | Lepidoptera_Pieridae | 3 | 0,012 |
| S4 | Lepidoptera_Nymphalidae | 3 | 0,012 |
| S4 | Diptera_Piophilidae | 2 | 0,008 |
| S4 | Diptera_Rhagionidae | 2 | 0,008 |
| S4 | Hymenoptera_Formicidae | 2 | 0,008 |
| S4 | Lepidoptera_Microlepidoptera (unranked) | 2 | 0,008 |
| S4 | Lepidoptera_Geometridae | 1 | 0,004 |
| S4 | Lepidoptera_Unknown | 1 | 0,004 |
| S4 | Hymenoptera_Colletidae | 1 | 0,004 |
| S4 | Coleoptera_Nitidulidae | 1 | 0,004 |
| S4 | Coleoptera_Cantharidae | 1 | 0,004 |
| S4 | Coleoptera_Cerambycidae | 1 | 0,004 |
| S4 | Coleoptera_Curculionidae | 1 | 0,004 |
| S4 | Diptera_Tachinidae | 1 | 0,004 |
| S4 | Diptera_Empididae | 1 | 0,004 |

Table S5. Pollinator families' abundance. *A list of pollinator families (or higher taxa) involved in observed plant–pollinator interactions, with their corresponding overall abundance (n° of visits) and relative abundance (n° of visits / n° tot visits) within stages.*

### ‘R’ code

We report here the code used for data analysis with some useful console outputs and comments. Data analysis was done using ‘R’ software, version 4.3.1. This document was compiled with the ‘rmarkdown’ package, version 2.25. The purpose of this document is to increase reproducibility, clarity, transparency and dissemination.

#### General preliminary operations

In the first section of the code, we carry out the preliminary operations necessary for the subsequent data analysis, including: loading libraries, importing raw data and transforming raw data from a table of field observations to plant-pollinator interaction matrices.

```
###LOAD PACKAGES###
library(bbmle)
library(betareg)
library(bipartite)
library(bmotif)
library(car)
library(ggalt)
library(ggrepel)
library(ggtext)
library(gridExtra)
library(tidyverse)
library(vegan)
library(visualize)

###IMPORT RAW DATASET###
raw_data <- read.csv("raw_data.csv", sep = ";")

###TRANSFORM RAW DATASET INTO SITES' BIPARTITE NETWORKS###
##create table of observations with weights (at 'site' level)##
#remove the first two rows (excluded quadrat sampling trials)
bipnet_sites <- raw_data[-c(1, 2), ]
#remove unnecessary columns
columns_to_remove <- c("Date", "Hour", "Stage", "Method", "Replicate",
                       "Replicate_ID", "Original_Tube_ID", "Tube_ID",
                       "Family_F", "Genus_F", "Genus_I", "Species_I",
                       "Type_of_sample")
bipnet_sites <- bipnet_sites[, !(colnames(bipnet_sites) %in% columns_to_remove)]
#remove leading and trailing whitespaces
bipnet_sites$Order_I <- trimws(bipnet_sites$Order_I)
bipnet_sites$Family_I <- trimws(bipnet_sites$Family_I)
bipnet_sites$Species_F <- trimws(bipnet_sites$Species_F)
bipnet_sites$Plot <- trimws(bipnet_sites$Plot)
#merge columns "Order_I" and "Family_I"
bipnet_sites$OrdFamily_I <- paste(bipnet_sites$Order_I,
                                  bipnet_sites$Family_I, sep = "_")
bipnet_sites <- bipnet_sites[, !(colnames(bipnet_sites) %in%
```

```
                                     c("Order_I", "Family_I"))]
#create new column "Weight"
bipnet_sites$Weight <- NA
#count repeating rows, remove copies and fill "Weight" column
weight_counts <- table(apply(bipnet_sites, 1, paste, collapse = ","))
for (row in names(weight_counts)) {
  if (weight_counts[row] > 1) {
    indices <- which(apply(bipnet_sites, 1, paste, collapse = ",") == row)
    bipnet_sites$Weight[indices] <- weight_counts[row]
    bipnet_sites <- bipnet_sites[-(indices[-1]), ]
  } else if (weight_counts[row] == 1) {
  indices <- which(do.call(paste, c(bipnet_sites, sep = ",")) == row)
  bipnet_sites$Weight[indices] <- 1
  }
}
##convert the table of observations into a list of bipartite networks##
bipnet_sites <- data.frame(higher = bipnet_sites$OrdFamily_I,
                           lower = bipnet_sites$Species_F,
                           webID = bipnet_sites$Plot,
                           freq = bipnet_sites$Weight)
bipnet_sites_list <- frame2webs(bipnet_sites, varnames = c("lower", "higher",
                                                            "webID", "freq"),
                                type.out = "list", emptylist = TRUE)

###TRANSFORM RAW DATASET INTO STAGES' BIPARTITE NETWORKS###
##create table of observations with weights (at 'stage' level)##
#remove the first two rows (excluded quadrat sampling trials)
bipnet_stages <- raw_data[-c(1, 2), ]
#remove unnecessary columns
columns_to_remove_2 <- c("Date", "Hour", "Plot", "Method", "Replicate",
                         "Replicate_ID", "Original_Tube_ID", "Tube_ID",
                         "Family_F", "Genus_F", "Genus_I", "Species_I",
                         "Type_of_sample")
bipnet_stages <- bipnet_stages[, !(colnames(bipnet_stages) %in%
                                       columns_to_remove_2)]
#remove leading and trailing whitespaces
bipnet_stages$Order_I <- trimws(bipnet_stages$Order_I)
bipnet_stages$Family_I <- trimws(bipnet_stages$Family_I)
bipnet_stages$Species_F <- trimws(bipnet_stages$Species_F)
bipnet_stages$Stage <- trimws(bipnet_stages$Stage)
#merge columns "Order_I" and "Family_I"
bipnet_stages$OrdFamily_I <- paste(bipnet_stages$Order_I,
                                    bipnet_stages$Family_I, sep = "_")
bipnet_stages <- bipnet_stages[, !(colnames(bipnet_stages) %in%
                                       c("Order_I", "Family_I"))]
#create new column "Weight"
bipnet_stages$Weight <- NA
#count repeating rows, remove copies and fill "Weight" column
weight_counts_2 <- table(apply(bipnet_stages, 1, paste, collapse = ","))
for (row in names(weight_counts_2)) {
```

```
  if (weight_counts_2[row] > 1) {
    indices_2 <- which(apply(bipnet_stages, 1, paste, collapse = ",") == row)
    bipnet_stages$Weight[indices_2] <- weight_counts_2[row]
    bipnet_stages <- bipnet_stages[-(indices_2[-1]), ]
  } else if (weight_counts_2[row] == 1) {
    indices_2 <- which(do.call(paste, c(bipnet_stages, sep = ",")) == row)
    bipnet_stages$Weight[indices_2] <- 1
  }
}
##convert the table of observations into a list of bipartite networks##
bipnet_stages <- data.frame(higher = bipnet_stages$OrdFamily_I,
                            lower = bipnet_stages$Species_F,
                            webID = bipnet_stages$Stage,
                            freq = bipnet_stages$Weight)
bipnet_stages_list <- frame2webs(bipnet_stages, varnames = c("lower", "higher",
                                                             "webID", "freq"),
                                 type.out = "list", emptylist = TRUE)
```

We look at pollinator families' abundance and relative abundance.

```
###POLLINATOR FAMILIES' ABUNDANCE###
##create dataframe with pollinator abundances##
#remove the first two rows (excluded quadrat sampling trials)
poll_ab <- raw_data[-c(1, 2), ]
#remove unnecessary columns
columns_to_remove_3 <- c("Date", "Hour", "Plot", "Method", "Replicate",
                         "Replicate_ID", "Original_Tube_ID", "Tube_ID",
                         "Family_F", "Genus_F", "Species_F", "Genus_I",
                         "Species_I", "Type_of_sample")
poll_ab <- poll_ab[, !(colnames(poll_ab) %in% columns_to_remove_3)]
#remove leading and trailing whitespaces
poll_ab$Order_I <- trimws(poll_ab$Order_I)
poll_ab$Family_I <- trimws(poll_ab$Family_I)
poll_ab$Stage <- trimws(poll_ab$Stage)
#merge columns "Order_I" and "Family_I"
poll_ab$OrdFamily_I <- paste(poll_ab$Order_I, poll_ab$Family_I, sep = "_")
poll_ab <- poll_ab[, !(colnames(poll_ab) %in% c("Order_I", "Family_I"))]
#create new column "Weight"
poll_ab$Weight <- NA
#count repeating rows, remove copies and fill "Weight" column
weight_counts_3 <- table(apply(poll_ab, 1, paste, collapse = ","))
for (row in names(weight_counts_3)) {
  if (weight_counts_3[row] > 1) {
    indices_3 <- which(apply(poll_ab, 1, paste, collapse = ",") == row)
    poll_ab$Weight[indices_3] <- weight_counts_3[row]
    poll_ab <- poll_ab[-(indices_3[-1]), ]
  } else if (weight_counts_3[row] == 1) {
    indices_3 <- which(do.call(paste, c(poll_ab, sep = ",")) == row)
    poll_ab$Weight[indices_3] <- 1
  }
}
```

```
#calculate relative abundances and upload them to "Weight_Rel" column
poll_ab <- poll_ab %>%
  group_by(Stage) %>%
  mutate(Weight_Rel = Weight / sum(Weight)) %>%
  ungroup()
##plot abundances histograms##
reorder_within <- function(x, by, within, fun = median, sep = "___", ...) {
  new_x <- paste(x, within, sep = sep)
  stats::reorder(new_x, -by, FUN = fun)
}
poll_ab <- poll_ab %>%
  group_by(Stage) %>%
  mutate(OrdFamily_I_reordered = reorder_within(OrdFamily_I,
                                                 Weight_Rel,
                                                 Stage)) %>%
  ungroup()
scale_x_reordered <- function(..., sep = "___") {
  ggplot2::scale_x_discrete(labels = function(x) gsub(paste0(sep, ".*"), "", x),
                            ...)
}
figure_S5 <- ggplot(poll_ab, aes(x = OrdFamily_I_reordered, y = Weight_Rel,
                                  fill = Stage)) +
  geom_bar(stat = "identity") +
  facet_wrap(~ Stage, scales = "free_x") +
  labs(x = "Pollinator Families", y = "Relative Abundance") +
  scale_x_reordered() +
  scale_fill_manual(values = c("S1" = "blue3", "S2" = "green3",
                               "S3" = "yellow3", "S4" = "red3")) +
  theme(
    axis.text.x = element_text(angle = 90, hjust = 1, size = 7),
    panel.grid.major = element_blank(),
    panel.grid.minor = element_blank(),
  )
figure_S5
```

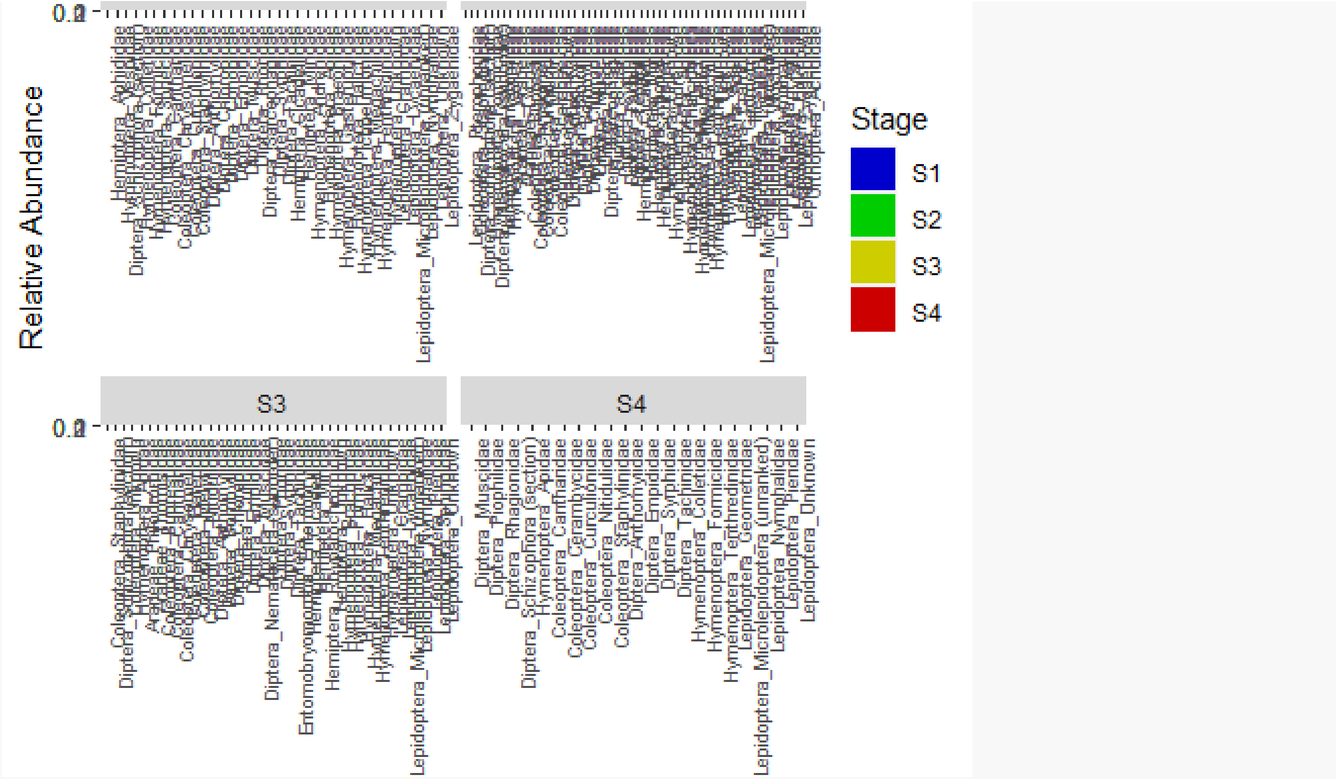

We assess, from a qualitative perspective, the degree of specialisation of species within stages' networks and of the overall networks themselves. This is done just to have an idea of how species differ in terms of specialisation along the glacier foreland.
We calculate species level specialisation using 'd'' index. The d' index, as outlined by Blüthgen et al. (2006), is derived from Kulback-Leibler distance. It quantifies the extent to which a species deviates from a random sampling of available interacting partners. Its values range between 0 (no specialisation) and 1 (perfect specialisation) (Dormann et al. 2023). For instance, within a pollination network, a pollinator solely present on a single plant species may still exhibit limited evidence of specialization if that plant species is predominant, resulting in a low d' value. Conversely, a pollinator associated with rare plants would have a substantially higher d' value (Dormann et al. 2023). 'd'' index is called through 'specieslevel' function of 'bipartite' package (Dormann et al. 2023).
We calculate network level specialisation using 'H2'' index. H2' describes the level of 'complementarity specialisation' of an entire bipartite network (Blüthgen et al. 2006). It describes to which extent observed interactions deviate from those that would be expected given the species marginal totals (Dormann et al. 2023). H2' is basically an extension of d' for the entire network (Dormann et al. 2023). 'H2'' index is called through 'networklevel' function of 'bipartite' package (Dormann et al. 2023).

```
###SPECIALISATION###
#initialise list and vector
spec_list <- list()
#loop through stages
for (i in names(bipnet_stages_list)) {
  #species level specialisation
  spec <- specieslevel(bipnet_stages_list[[i]], index = "d")
```

```r
  spec_list[[i]] <- spec
  mean_spec_lower <- mean(spec_list[[i]]$`lower level`$d, na.rm = TRUE)
  mean_spec_higher <- mean(spec_list[[i]]$`higher level`$d, na.rm = TRUE)
  print(spec)
  cat("Mean specialisation for plants in", i, "is:", mean_spec_lower, "\n")
  cat("Mean specialisation for pollinators in", i, "is:", mean_spec_higher,
      "\n")
  #network level specialisation
  spec_n[i] <- networklevel(bipnet_stages_list[[i]], index = "H2")
  cat("Overall specialisation for", i, "is:", spec_n[i], "\n")
}
```

```
## $`higher level`
##                                              d
## Coleoptera_Cantharidae               0.16830940
## Coleoptera_Chrysomelidae             0.54536919
## Coleoptera_Curculionidae             0.58370530
## Coleoptera_Staphylinidae             0.16104477
## Diptera_Anthomyiidae                 0.30979410
## Diptera_Chloropidae                  0.35014849
## Diptera_Conopidae                    0.34586054
## Diptera_Empididae                    0.31739242
## Diptera_Muscidae                     0.40429403
## Diptera_Phoridae                     0.22580062
## Diptera_Sarcophagidae                0.16104477
## Diptera_Schizophora (section)        0.20179255
## Diptera_Syrphidae                    0.16092993
## Diptera_Tachinidae                   0.16104477
## Hemiptera_Aphididae                  0.30618543
## Hemiptera_Cicadellidae               0.33352625
## Hemiptera_Miridae                    0.10681415
## Hymenoptera_Andrenidae               0.41269778
## Hymenoptera_Apidae                   0.39297457
## Hymenoptera_Braconidae               0.07320546
## Hymenoptera_Colletidae               0.38737532
## Hymenoptera_Formicidae               0.20775641
## Hymenoptera_Gasteruptiidae           0.58370530
## Hymenoptera_Halictidae               0.53488635
## Hymenoptera_Ichneumonidae            0.34347209
## Hymenoptera_Megachilidae             0.43506178
## Hymenoptera_Sphecidae                0.17460444
## Hymenoptera_Tenthredinidae           0.24414670
## Hymenoptera_Unknown                  0.30526261
## Hymenoptera_Vespidae                 0.26164708
## Lepidoptera_Crambidae                0.45897644
## Lepidoptera_Lycaenidae               0.35883497
## Lepidoptera_Microlepidoptera (unranked) 1.00000000
## Lepidoptera_Nymphalidae              0.16104477
## Lepidoptera_Unknown                  0.26574199
## Lepidoptera_Zygaenidae               0.24977219
##
```

```
## $`lower level`
##                                  d
## Achillea erba-rotta       0.5132535
## Anthyllis vulneraria      0.4262086
## Campanula barbata         0.3755983
## Campanula cochleariifolia 1.0000000
## Cerastium alpinum         0.4321013
## Cerastium arvense         0.5183481
## Epilobium fleischeri      0.4390703
## Erigeron alpinus          0.0000000
## Hieracium angustifolium   0.5000000
## Hieracium staticifolium   0.1880260
## Leontodon hispidus        0.3722499
## Leucanthemopsis alpina    0.4347294
## Leucanthemum adustum      0.2043354
## Leucanthemum vulgare      0.3018570
## Linaria alpina            0.0000000
## Lotus corniculatus        0.2374167
## Minuartia verna           0.4543354
## Pedicularis tuberosa      0.1021266
## Pilosella cymosa          0.2858749
## Pilosella officinarum     0.6494223
## Rumex scutatus            0.1238612
## Saxifraga bryoides        0.3058352
## Saxifraga paniculata      0.3718434
## Sempervivum arachnoideum  0.6015087
## Thymus praecox            0.3444293
## Trifolium ochroleucon     0.4287737
## Trifolium pallescens      0.2946203
##
## Mean specialisation for plants in S1 is: 0.3668824
## Mean specialisation for pollinators in S1 is: 0.3248395
## Overall specialisation for S1 is: 0.3617461
## $`higher level`
##                                                d
## Araneae_Araneidae                     0.30819075
## Araneae_Lycosidae                     0.06172369
## Coleoptera_Buprestidae                0.22054321
## Coleoptera_Chrysomelidae              0.31874210
## Coleoptera_Curculionidae              0.22851091
## Coleoptera_Melyridae                  0.29890437
## Coleoptera_Scarabaeidae               0.33647549
## Coleoptera_Staphylinidae              0.26474278
## Coleoptera_Unknown                    0.36405575
## Diptera_Anthomyiidae                  0.24785847
## Diptera_Bibionidae                    0.06172369
## Diptera_Bombyliidae                   0.53226492
## Diptera_Chloropidae                   0.26456004
## Diptera_Muscidae                      0.24865904
## Diptera_Nematocera (suborder)         0.72552321
## Diptera_Phoridae                      0.06172369
```

```
## Diptera_Sarcophagidae                     0.35354010
## Diptera_Schizophora (section)             0.18266024
## Diptera_Sciaridae                         0.06172369
## Diptera_Syrphidae                         0.21196979
## Diptera_Tachinidae                        0.28458781
## Diptera_Tephritidae                       0.75929718
## Hemiptera_Cicadellidae                    0.20636041
## Hemiptera_Miridae                         0.27390752
## Hemiptera_Pentatomidae                    0.92742191
## Hemiptera_Rhopalidae                      0.21672029
## Hemiptera_Scutelleridae                   0.27084294
## Hemiptera_Unknown                         0.06172369
## Hymenoptera_Andrenidae                    0.35681098
## Hymenoptera_Apidae                        0.61525783
## Hymenoptera_Chalcidoidea                  0.06172369
## Hymenoptera_Colletidae                    0.52359149
## Hymenoptera_Formicidae                    0.51186293
## Hymenoptera_Halictidae                    0.38231783
## Hymenoptera_Ichneumonidae                 0.20281115
## Hymenoptera_Megachilidae                  0.33492318
## Hymenoptera_Tenthredinidae                0.32923323
## Hymenoptera_Unknown                       0.25618301
## Hymenoptera_Vespidae                      0.22893698
## Lepidoptera_Adelidae                      0.28812246
## Lepidoptera_Crambidae                     0.24239236
## Lepidoptera_Geometridae                   0.65825935
## Lepidoptera_Hesperiidae                   0.22761167
## Lepidoptera_Lycaenidae                    0.60601498
## Lepidoptera_Microlepidoptera (unranked) 0.32518665
## Lepidoptera_Noctuidae                     0.38627174
## Lepidoptera_Nymphalidae                   0.26798454
## Lepidoptera_Pieridae                      0.32518665
## Lepidoptera_Pterophoridae                 0.60087359
## Lepidoptera_Pyralidae                     0.33422357
## Lepidoptera_Unknown                       0.45630362
## Lepidoptera_Zygaenidae                    0.62649456
## Orthoptera_Acrididae                      0.33463870
##
## $`lower level`
##                                  d
## Achillea erba-rotta      0.23023700
## Adenostyles alpina       0.15963803
## Alchemilla monticola     0.22640248
## Antennaria dioica        0.14961842
## Anthyllis vulneraria     0.64082786
## Bartsia alpina           0.95949762
## Campanula barbata        0.36074464
## Campanula scheuchzeri    0.29839619
## Carduus defloratus       0.53913294
## Centaurea nervosa        0.45568524
## Cerastium arvense        0.23460428
```

```
## Chaerophyllum villarsii  0.05291702
## Epilobium fleischeri     0.27461657
## Erigeron alpinus         0.15963803
## Galium anisophyllon      0.38879964
## Gymnadenia conopsea      0.02949657
## Hieracium angustifolium  0.25182559
## Hieracium murorum        0.35975428
## Hieracium staticifolium  0.31859048
## Hippocrepis comosa       0.32557706
## Leontodon helveticus     0.36302193
## Leontodon hispidus       0.25904315
## Leucanthemopsis alpina   0.28244549
## Leucanthemum vulgare     0.36690318
## Lotus corniculatus       0.37435281
## Myosotis alpestris       0.42884683
## Orchis mascula           0.15963803
## Phyteuma betonicifolium  0.43660521
## Pilosella cymosa         0.36940448
## Pilosella officinarum    0.23508758
## Polygonum viviparum      0.00000000
## Potentilla aurea         0.19238714
## Ranunculus montanus      0.28770125
## Ranunculus villarsii     0.16985608
## Rhinanthus minor         0.02003770
## Rhododendron ferrugineum 0.29117991
## Rumex scutatus           0.51070827
## Saxifraga aizoides       0.48264046
## Saxifraga paniculata     0.26193892
## Silene rupestris         0.15153720
## Silene vulgaris          0.12597866
## Solidago virgaurea       0.00000000
## Thymus praecox           0.54787436
## Tofieldia calyculata     0.01025392
## Trifolium badium         0.35811077
## Trifolium hybridum       0.15963803
## Trifolium ochroleucon    0.33250310
## Trifolium pallescens     0.49544526
## Trifolium pratense       0.41798011
## Veronica fruticans       0.33730808
## 
## Mean specialisation for plants in S2 is: 0.2968886
## Mean specialisation for pollinators in S2 is: 0.3365693
## Overall specialisation for S2 is: 0.3622554
## $`higher level`
##                                               d
## Araneae_Philodromidae                0.13091532
## Araneae_Thomisidae                   0.19832330
## Coleoptera_Buprestidae               0.36186381
## Coleoptera_Cantharidae               0.25578533
## Coleoptera_Chrysomelidae             0.17866648
## Coleoptera_Elateridae                0.00000000
```

```
## Coleoptera_Melyridae                        0.68838246
## Coleoptera_Nitidulidae                      0.28576335
## Coleoptera_Staphylinidae                    0.43388717
## Coleoptera_Unknown                          0.39317550
## Diptera_Anthomyiidae                        0.28519312
## Diptera_Bombyliidae                         0.79842277
## Diptera_Chloropidae                         0.53271914
## Diptera_Culicidae                           0.35482234
## Diptera_Empididae                           0.27161859
## Diptera_Muscidae                            0.10813870
## Diptera_Nematocera (suborder)               0.57457587
## Diptera_Phoridae                            0.38466882
## Diptera_Schizophora (section)               0.10927788
## Diptera_Syrphidae                           0.20829763
## Diptera_Tachinidae                          0.56757437
## Entomobryomorpha_Entomobryidae              0.28576335
## Hemiptera_Cicadellidae                      0.38981919
## Hemiptera_Miridae                           0.44222563
## Hemiptera_Rhyparochromidae                  0.51887519
## Hemiptera_Unknown                           0.08284698
## Hymenoptera_Apidae                          0.71161038
## Hymenoptera_Braconidae                      0.50337742
## Hymenoptera_Formicidae                      0.12968882
## Hymenoptera_Halictidae                      0.57875069
## Hymenoptera_Megachilidae                    0.62323508
## Hymenoptera_Tenthredinidae                  0.23005280
## Hymenoptera_Unknown                         0.20526836
## Lepidoptera_Crambidae                       0.43488217
## Lepidoptera_Lycaenidae                      0.60687703
## Lepidoptera_Microlepidoptera (unranked) 0.21209817
## Lepidoptera_Nymphalidae                     0.29569470
## Lepidoptera_Pieridae                        0.25748558
## Lepidoptera_Sphingidae                      0.14677602
## Lepidoptera_Unknown                         0.52033668
##
## $`lower level`
##                                  d
## Achillea erba-rotta      0.12305743
## Achillea millefolium     0.04592794
## Anthyllis vulneraria     0.51043000
## Campanula barbata        0.34980391
## Campanula scheuchzeri    0.14410663
## Carduus defloratus       0.23224326
## Centaurea nervosa        0.37629809
## Cerastium arvense        0.21254231
## Crepis pontana           0.00000000
## Dryas octopetala         0.18387911
## Epilobium fleischeri     0.41145702
## Gentiana nivalis         0.14927357
## Hieracium murorum        0.21470918
## Hieracium staticifolium  0.43449048
```

```
## Hippocrepis comosa       0.68918969
## Huguenina tanacetifolia  0.31888217
## Leontodon hispidus       0.18321934
## Leucanthemum vulgare     0.19050418
## Lotus corniculatus       0.39963043
## Myosotis alpestris       0.09577516
## Peucedanum ostruthium    0.23897407
## Phyteuma betonicifolium  0.21363615
## Pilosella officinarum    0.25559813
## Potentilla aurea         0.23105214
## Ranunculus montanus      0.21230486
## Ranunculus villarsii     0.20188276
## Rhododendron ferrugineum 0.37298074
## Sempervivum arachnoideum 0.86166481
## Silene dioica            0.00000000
## Silene vulgaris          0.11945703
## Thymus praecox           0.28948793
## Trifolium hybridum       0.64240872
## Trifolium ochroleucon    0.24571100
## Trifolium pallescens     0.48059961
## Trifolium pratense       0.28257848
## Trifolium repens         0.52143887
## Vaccinium vitis-idaea    0.09115778
## Veronica fruticans       0.52953896
##
## Mean specialisation for plants in S3 is: 0.2909445
## Mean specialisation for pollinators in S3 is: 0.3574434
## Overall specialisation for S3 is: 0.326648
## $`higher level`
##                                                  d
## Coleoptera_Cantharidae                  0.00000000
## Coleoptera_Cerambycidae                 0.00000000
## Coleoptera_Curculionidae                0.42763284
## Coleoptera_Nitidulidae                  0.45095688
## Coleoptera_Staphylinidae                0.22397860
## Diptera_Anthomyiidae                    0.62390540
## Diptera_Empididae                       0.46373714
## Diptera_Muscidae                        0.10386341
## Diptera_Piophilidae                     0.68180238
## Diptera_Rhagionidae                     0.00000000
## Diptera_Schizophora (section)           0.09344167
## Diptera_Syrphidae                       0.13525475
## Diptera_Tachinidae                      0.72547844
## Hymenoptera_Apidae                      0.32615860
## Hymenoptera_Colletidae                  1.00000000
## Hymenoptera_Formicidae                  0.00000000
## Hymenoptera_Tenthredinidae              0.33625694
## Lepidoptera_Geometridae                 0.00000000
## Lepidoptera_Microlepidoptera (unranked) 0.43704354
## Lepidoptera_Nymphalidae                 0.03102186
## Lepidoptera_Pieridae                    0.00000000
```

```
## Lepidoptera_Unknown                      0.56489362
##
## $`lower level`
##                                         d
## Achillea erba-rotta      0.27989275
## Anthyllis vulneraria     0.43558951
## Campanula rhomboidalis   0.18464277
## Dactylorhiza maculata    0.46638095
## Hieracium murorum        0.32360535
## Leontodon hispidus       0.34176598
## Lotus corniculatus       0.19706046
## Peucedanum ostruthium    0.22868006
## Potentilla aurea         0.30007797
## Ranunculus montanus      0.36131383
## Ranunculus platanifolius 0.03628359
## Ranunculus villarsii     0.25943601
## Rhododendron ferrugineum 0.23824985
## Sempervivum montanum     1.00000000
## Vaccinium vitis-idaea    0.03628359
## Valeriana tripteris      0.22420971
## Veronica chamaedrys      0.33312545
## Vicia sepium             0.14768876
##
## Mean specialisation for plants in S4 is: 0.2996826
## Mean specialisation for pollinators in S4 is: 0.3011557
## Overall specialisation for S4 is: 0.3182095
```

#### (a) Motifs' prevalence

In this section, we unfold all analyses concerning the frequency of occurrence of motifs.

We start by calculating the frequency of occurrence of all possible motifs up to 6 nodes in observed networks (at the 'site' level) using 'mcount' function of 'bmotif' package (Mora et al. 2018, Simmons et al. 2019b).

```
###CALCULATE MOTIFS' OCCURRENCES (SAMPLED NETWORKS)###
site_names <- c("1A", "1B", "1C", "1D", "2A", "2B", "2C", "2D", "3A", "3B",
                "3C", "3D", "4A", "4B", "4C", "4D")
for (site in site_names) {
  assign(paste("mcount_", site, sep = ""), mcount(bipnet_sites_list[[site]],
                                                   six_node = TRUE,
                                                   normalisation = TRUE,
                                                   mean_weight = TRUE,
                                                   standard_dev = TRUE))
}
```

We then identify stages' most representative motifs (those with highest mean normalised frequency of occurrence across sites).

```r
###STAGES' MOST REPRESENTATIVE MOTIFS###
#calculate mean occurrence across sites for each stage
mean_occurrence <- function(..., na.rm = TRUE) {
  data <- cbind(...)
  rowMeans(data, na.rm = na.rm)
}
mcount_S1 <- data.frame(normalise_sum = mean_occurrence(mcount_1A$normalise_sum,
                                                         mcount_1B$normalise_sum,
                                                         mcount_1C$normalise_sum,
                                                         mcount_1D$normalise_sum)
                        )
mcount_S2 <- data.frame(normalise_sum = mean_occurrence(mcount_2A$normalise_sum,
                                                         mcount_2B$normalise_sum,
                                                         mcount_2C$normalise_sum,
                                                         mcount_2D$normalise_sum)
                        )
mcount_S3 <- data.frame(normalise_sum = mean_occurrence(mcount_3A$normalise_sum,
                                                         mcount_3B$normalise_sum,
                                                         mcount_3C$normalise_sum,
                                                         mcount_3D$normalise_sum)
                        )
mcount_S4 <- data.frame(normalise_sum = mean_occurrence(mcount_4A$normalise_sum,
                                                         mcount_4B$normalise_sum,
                                                         mcount_4C$normalise_sum,
                                                         mcount_4D$normalise_sum)
                        )
#find top n most representative motifs
find_most_representative_motifs <- function(df, name, top_n = 1) {
  top_indices <- order(df$normalise_sum, decreasing = TRUE)[1:top_n]
  top_motifs <- rownames(df)[top_indices]
  cat("top", top_n, "most representative motifs of", name, "are",
      paste(top_motifs, collapse = ", "), "\n")
}
find_most_representative_motifs(mcount_S1, "Stage1", top_n = 3)

## top 3 most representative motifs of Stage1 are 39, 27, 38

find_most_representative_motifs(mcount_S2, "Stage2", top_n = 3)

## top 3 most representative motifs of Stage2 are 19, 25, 27

find_most_representative_motifs(mcount_S3, "Stage3", top_n = 3)

## top 3 most representative motifs of Stage3 are 38, 25, 13

find_most_representative_motifs(mcount_S4, "Stage4", top_n = 3)

## top 3 most representative motifs of Stage4 are 17, 7, 44
```

We build null model networks by randomising sites' networks 100 times following 4 different null model approaches (varying in level of conservativism and reshuffling method). For each null model approach, we calculate one-sample Z-scores and identify the most over- and under-represented motifs in each stage.

One-sample Z-score is defined as $\boldsymbol{Z} = \frac{x-\mu}{\sigma}$, where $\boldsymbol{x}$ is the motif frequency calculated on observed networks, $\boldsymbol{\mu}$ and $\boldsymbol{\sigma}$ are the mean and standard deviation of the 100 values of null-model motif frequencies, respectively. In a two-tailed normal distribution, the threshold for 5% confidence level (corresponding to a p-value of 0.05) is |Z|=1.96 (if |Z|>1.96, then p<0.05).

• Reference null model: "r00_ind" ('vegan' package). This method preserves grand sum and individuals are shuffled among cells of the matrix (Oksanen et al., 2022).

• Additional null models:

- 'r0_ind' ('vegan' package). This method preserves row sums and individuals are shuffled among cells of each row of the matrix (Oksanen et al., 2022).

- 'c0_ind' ('vegan' package). This method preserves column sums and individuals are shuffled among cells of each column of the matrix (Oksanen et al., 2022).

- 'vaznull' ('bipartite' package). This method produces a null model network with the main constraint of having the same connectance of the original network. Observed plant–pollinator interactions are randomised based on species-specific probabilities that are proportional to species' relative abundances (Dormann et al., 2023).

```
###REFERENCE NULL MODEL ("r00_ind")###
##initialise Z-scores dataframe##
z_scores_dataframe <- data.frame(matrix(NA, nrow = 44, ncol = 16))
##randomise networks, calculate motifs' occurrences and calculate Z-scores##
#randomise
for (i in seq_along(bipnet_sites_list)) {
  bipnet_null <- vegan::nullmodel(bipnet_sites_list[[i]], "r00_ind")
  bipnet_null_2 <- simulate(bipnet_null, nsim = 100, seed = 1)
  bipnet_null_2_list <- vector("list", length = 100)
  #upload simulations into the list
  for (k in seq_len(100)) {
    bipnet_null_2_list[[k]] <- as.matrix(bipnet_null_2[, , k])
    rownames(bipnet_null_2_list[[k]]) <- NULL
    colnames(bipnet_null_2_list[[k]]) <- NULL
  }
  results_matrix <- matrix(NA, nrow = 44, ncol = 100)
  #calculate motifs' occurrences
  for (j in 1:100) {
    mcount_null <- mcount(bipnet_null_2_list[[j]], six_node = TRUE,
                          normalisation = TRUE, mean_weight = TRUE,
                          standard_dev = TRUE)
    normalise_sum_column <- mcount_null$normalise_sum
    results_matrix[, j] <- normalise_sum_column
  }
  #calculate mean null model value for each motif
  normalise_sum_null_mean <- rowMeans(results_matrix, na.rm = TRUE)
  #calculate standard deviation on null model values
  sd_null_mcount <- apply(results_matrix, 1, sd, na.rm = TRUE)
  #get mcount dataframes' names
```

```
  mcount_df_name <- paste0("mcount_", names(bipnet_sites_list)[i])
  #calculate Z-scores
  z_scores <- (((get(mcount_df_name)$normalise_sum) - normalise_sum_null_mean) /
                 (sd_null_mcount))
  #upload Z-scores in the dataframe
  z_scores_dataframe[, i] <- z_scores
  #rename columns
  col_name <- paste0("z_score_", names(bipnet_sites_list)[i])
  colnames(z_scores_dataframe)[i] <- col_name
  #replace infinite values with NaNs
  z_scores_dataframe[z_scores_dataframe == -Inf] <- NaN
  z_scores_dataframe[z_scores_dataframe == Inf] <- NaN
}
##percentage of entries with |Z|>1.96 across all stages and sites##
num_entries_above_threshold <- sum(abs(z_scores_dataframe) > 1.96 &
                                     !is.na(z_scores_dataframe))
total_entries_adjusted <- sum(!is.na(z_scores_dataframe))
percentage_above_threshold <- (num_entries_above_threshold /
                                 total_entries_adjusted) * 100
cat("number of entries above threshold:", num_entries_above_threshold, "\n")

## number of entries above threshold: 460

cat("percentage of entries above threshold:", percentage_above_threshold, "%\n")

## percentage of entries above threshold: 74.07407 %

##percentage of entries with Z>1.96 across all stages and sites##
num_entries_above_threshold_over <- sum(z_scores_dataframe > 1.96 &
                                          !is.na(z_scores_dataframe))
percentage_above_threshold_over <- (num_entries_above_threshold_over /
                                      total_entries_adjusted) * 100
cat("number of over-represented entries:", num_entries_above_threshold_over,
    "\n")

## number of over-represented entries: 279

cat("percentage of over-represented entries:", percentage_above_threshold_over,
    "%\n")

## percentage of over-represented entries: 44.92754 %

##percentage of entries with Z<-1.96 across all stages and sites##
num_entries_above_threshold_under <- sum(z_scores_dataframe < -1.96 &
                                           !is.na(z_scores_dataframe))
percentage_above_threshold_under <- (num_entries_above_threshold_under /
                                       total_entries_adjusted) * 100
cat("number of under-represented entries:", num_entries_above_threshold_under,
    "\n")
```

```
## number of under-represented entries: 181

cat("percentage of under-represented entries:", percentage_above_threshold_under
    , "%\n")

## percentage of under-represented entries: 29.14654 %

##stages' most over-represented motifs##
#divide Z-scores dataframe into separate dataframes (one per stage)
z_scores_S1 <- z_scores_dataframe[, 1:4]
z_scores_S2 <- z_scores_dataframe[, 5:8]
z_scores_S3 <- z_scores_dataframe[, 9:12]
z_scores_S4 <- z_scores_dataframe[, 13:16]
#calculate mean Z-score accross sites within each stage
z_scores_stages_mean <- data.frame(
  Stage1 = rowMeans(z_scores_S1, na.rm = TRUE),
  Stage2 = rowMeans(z_scores_S2, na.rm = TRUE),
  Stage3 = rowMeans(z_scores_S3, na.rm = TRUE),
  Stage4 = rowMeans(z_scores_S4, na.rm = TRUE)
)
#find max value
max_row_indices <- apply(z_scores_stages_mean, 2, which.max)
#print results
messages <- character(4)
for (i in 1:4) {
  column_name <- colnames(z_scores_stages_mean)[i]
  max_row_name <- rownames(z_scores_stages_mean)[max_row_indices[i]]
  messages[i] <- paste("the most over-represented motif of", column_name,
                       "is n°", max_row_name)
}
cat(messages, sep = "\n")

## the most over-represented motif of Stage1 is n° 1

## the most over-represented motif of Stage2 is n° 4

## the most over-represented motif of Stage3 is n° 1

## the most over-represented motif of Stage4 is n° 25

##stages' most under-represented motifs##
#find min value
min_row_indices_2 <- apply(z_scores_stages_mean, 2, which.min)
#print results
messages_2 <- character(4)
for (i in 1:4) {
  column_name_2 <- colnames(z_scores_stages_mean)[i]
  min_row_name_2 <- rownames(z_scores_stages_mean)[min_row_indices_2[i]]
  messages_2[i] <- paste("the most under-represented motif of", column_name_2,
                         "is n°", min_row_name_2)
}
```

```r
cat(messages_2, sep = "\n")

## the most under-represented motif of Stage1 is n° 34

## the most under-represented motif of Stage2 is n° 34

## the most under-represented motif of Stage3 is n° 41

## the most under-represented motif of Stage4 is n° 8

###ADDITIONAL NULL MODELS ("r0_ind", "c0_ind" and "vaznull")###
###"r0_ind"###
##initialise Z-scores dataframe##
z_scores_dataframe_r <- data.frame(matrix(NA, nrow = 44, ncol = 16))
##randomise networks, calculate motifs' occurrences and calculate Z-scores##
#randomise
for (i in seq_along(bipnet_sites_list)) {
  bipnet_null_r <- vegan::nullmodel(bipnet_sites_list[[i]], "r0_ind")
  bipnet_null_2_r <- simulate(bipnet_null_r, nsim = 100, seed = 1)
  bipnet_null_2_list_r <- vector("list", length = 100)
  #upload simulations into the list
  for (k in seq_len(100)) {
    bipnet_null_2_list_r[[k]] <- as.matrix(bipnet_null_2_r[, , k])
    rownames(bipnet_null_2_list_r[[k]]) <- NULL
    colnames(bipnet_null_2_list_r[[k]]) <- NULL
  }
  results_matrix_r <- matrix(NA, nrow = 44, ncol = 100)
  #calculate motifs' occurrences
  for (j in 1:100) {
    mcount_null_r <- mcount(bipnet_null_2_list_r[[j]], six_node = TRUE,
                            normalisation = TRUE, mean_weight = TRUE,
                            standard_dev = TRUE)
    normalise_sum_column_r <- mcount_null_r$normalise_sum
    results_matrix_r[, j] <- normalise_sum_column_r
  }
  #calculate mean null model value for each motif
  normalise_sum_null_mean_r <- rowMeans(results_matrix_r, na.rm = TRUE)
  #calculate standard deviation on null model values
  sd_null_mcount_r <- apply(results_matrix_r, 1, sd, na.rm = TRUE)
  #get mcount dataframes' names
  mcount_df_name_r <- paste0("mcount_", names(bipnet_sites_list)[i])
  #calculate Z-scores
  z_scores_r <- (((get(mcount_df_name_r)$normalise_sum) -
                  normalise_sum_null_mean_r) / (sd_null_mcount_r))
  #upload Z-scores in the dataframe
  z_scores_dataframe_r[, i] <- z_scores_r
  #rename columns
  col_name_r <- paste0("z_score_", names(bipnet_sites_list)[i])
  colnames(z_scores_dataframe_r)[i] <- col_name_r
  #replace infinite values with NaNs
```

```r
  z_scores_dataframe_r[z_scores_dataframe_r == -Inf] <- NaN
  z_scores_dataframe_r[z_scores_dataframe_r == Inf] <- NaN
}
##percentage of entries with |Z|>1.96 across all stages and sites##
num_entries_above_threshold_r <- sum(abs(z_scores_dataframe_r) > 1.96 &
                                       !is.na(z_scores_dataframe_r))
total_entries_adjusted_r <- sum(!is.na(z_scores_dataframe_r))
percentage_above_threshold_r <- (num_entries_above_threshold_r /
                                   total_entries_adjusted_r) * 100
cat("number of entries above threshold:", num_entries_above_threshold_r,
    "\n")

## number of entries above threshold: 451

cat("percentage of entries above threshold:", percentage_above_threshold_r,
    "%\n")

## percentage of entries above threshold: 75.92593 %

##percentage of entries with Z>1.96 across all stages and sites##
num_entries_above_threshold_over_r <- sum(z_scores_dataframe_r > 1.96 &
                                            !is.na(z_scores_dataframe_r))
percentage_above_threshold_over_r <- (num_entries_above_threshold_over_r /
                                        total_entries_adjusted_r) * 100
cat("number of over-represented entries:", num_entries_above_threshold_over_r,
    "\n")

## number of over-represented entries: 261

cat("percentage of over-represented entries:", percentage_above_threshold_over_r
    , "%\n")

## percentage of over-represented entries: 43.93939 %

##percentage of entries with Z<-1.96 across all stages and sites##
num_entries_above_threshold_under_r <- sum(z_scores_dataframe_r < -1.96 &
                                             !is.na(z_scores_dataframe_r))
percentage_above_threshold_under_r <- (num_entries_above_threshold_under_r /
                                         total_entries_adjusted_r) * 100
cat("number of under-represented entries:", num_entries_above_threshold_under_r,
    "\n")

## number of under-represented entries: 190

cat("percentage of under-represented entries:",
    percentage_above_threshold_under_r, "%\n")

## percentage of under-represented entries: 31.98653 %

##stages' most over-represented motifs##
```

```r
#divide Z-scores dataframe into separate dataframes (one per stage)
z_scores_S1_r <- z_scores_dataframe_r[, 1:4]
z_scores_S2_r <- z_scores_dataframe_r[, 5:8]
z_scores_S3_r <- z_scores_dataframe_r[, 9:12]
z_scores_S4_r <- z_scores_dataframe_r[, 13:16]
#calculate mean Z-score accross sites within each stage
z_scores_stages_mean_r <- data.frame(
  Stage1 = rowMeans(z_scores_S1_r, na.rm = TRUE),
  Stage2 = rowMeans(z_scores_S2_r, na.rm = TRUE),
  Stage3 = rowMeans(z_scores_S3_r, na.rm = TRUE),
  Stage4 = rowMeans(z_scores_S4_r, na.rm = TRUE)
)
#find max value
max_row_indices_r <- apply(z_scores_stages_mean_r, 2, which.max)
#print results
messages_r <- character(4)
for (i in 1:4) {
  column_name_r <- colnames(z_scores_stages_mean_r)[i]
  max_row_name_r <- rownames(z_scores_stages_mean_r)[max_row_indices_r[i]]
  messages_r[i] <- paste("the most over-represented motif of", column_name_r,
                         "is n°", max_row_name_r)
}
cat(messages_r, sep = "\n")

## the most over-represented motif of Stage1 is n° 3

## the most over-represented motif of Stage2 is n° 18

## the most over-represented motif of Stage3 is n° 18

## the most over-represented motif of Stage4 is n° 19

##stages' most under-represented motifs##
#find min value
min_row_indices_2_r <- apply(z_scores_stages_mean_r, 2, which.min)
#print results
messages_2_r <- character(4)
for (i in 1:4) {
  column_name_2_r <- colnames(z_scores_stages_mean_r)[i]
  min_row_name_2_r <- rownames(z_scores_stages_mean_r)[min_row_indices_2_r[i]]
  messages_2_r[i] <- paste("the most under-represented motif of",
                           column_name_2_r, "is n°", min_row_name_2_r)
}
cat(messages_2_r, sep = "\n")

## the most under-represented motif of Stage1 is n° 17

## the most under-represented motif of Stage2 is n° 40

## the most under-represented motif of Stage3 is n° 40
```

```r
## the most under-represented motif of Stage4 is n° 8

###"c0_ind"###
##initialise Z-scores dataframe##
z_scores_dataframe_c <- data.frame(matrix(NA, nrow = 44, ncol = 16))
##randomise networks, calculate motifs' occurrences and calculate Z-scores##
#randomise
for (i in seq_along(bipnet_sites_list)) {
  bipnet_null_c <- vegan::nullmodel(bipnet_sites_list[[i]], "c0_ind")
  bipnet_null_2_c <- simulate(bipnet_null_c, nsim = 100, seed = 1)
  bipnet_null_2_list_c <- vector("list", length = 100)
  #upload simulations into the list
  for (k in seq_len(100)) {
    bipnet_null_2_list_c[[k]] <- as.matrix(bipnet_null_2_c[, , k])
    rownames(bipnet_null_2_list_c[[k]]) <- NULL
    colnames(bipnet_null_2_list_c[[k]]) <- NULL
  }
  results_matrix_c <- matrix(NA, nrow = 44, ncol = 100)
  #calculate motifs' occurrences
  for (j in 1:100) {
    mcount_null_c <- mcount(bipnet_null_2_list_c[[j]], six_node = TRUE,
                            normalisation = TRUE, mean_weight = TRUE,
                            standard_dev = TRUE)
    normalise_sum_column_c <- mcount_null_c$normalise_sum
    results_matrix_c[, j] <- normalise_sum_column_c
  }
  #calculate mean null model value for each motif
  normalise_sum_null_mean_c <- rowMeans(results_matrix_c, na.rm = TRUE)
  #calculate standard deviation on null model values
  sd_null_mcount_c <- apply(results_matrix_c, 1, sd, na.rm = TRUE)
  #get mcount dataframes' names
  mcount_df_name_c <- paste0("mcount_", names(bipnet_sites_list)[i])
  #calculate Z-scores
  z_scores_c <- (((get(mcount_df_name_c)$normalise_sum) -
                    normalise_sum_null_mean_c) / (sd_null_mcount_c))
  #upload Z-scores in the dataframe
  z_scores_dataframe_c[, i] <- z_scores_c
  #rename columns
  col_name_c <- paste0("z_score_", names(bipnet_sites_list)[i])
  colnames(z_scores_dataframe_c)[i] <- col_name_c
  #replace infinite values with NaNs
  z_scores_dataframe_c[z_scores_dataframe_c == -Inf] <- NaN
  z_scores_dataframe_c[z_scores_dataframe_c == Inf] <- NaN
}
##percentage of entries with |Z|>1.96 across all stages and sites##
num_entries_above_threshold_c <- sum(abs(z_scores_dataframe_c) > 1.96 &
                                       !is.na(z_scores_dataframe_c))
total_entries_adjusted_c <- sum(!is.na(z_scores_dataframe_c))
percentage_above_threshold_c <- (num_entries_above_threshold_c /
                                   total_entries_adjusted_c) * 100
```

```r
cat("number of entries above threshold:", num_entries_above_threshold_c, "\n")

## number of entries above threshold: 438

cat("percentage of entries above threshold:", percentage_above_threshold_c,
    "%\n")

## percentage of entries above threshold: 71.1039 %

##percentage of entries with Z>1.96 across all stages and sites##
num_entries_above_threshold_over_c <- sum(z_scores_dataframe_c > 1.96 &
                                          !is.na(z_scores_dataframe_c))
percentage_above_threshold_over_c <- (num_entries_above_threshold_over_c /
                                      total_entries_adjusted_c) * 100
cat("number of over-represented entries:", num_entries_above_threshold_over_c,
    "\n")

## number of over-represented entries: 222

cat("percentage of over-represented entries:", percentage_above_threshold_over_c
    , "%\n")

## percentage of over-represented entries: 36.03896 %

##percentage of entries with Z<-1.96 across all stages and sites##
num_entries_above_threshold_under_c <- sum(z_scores_dataframe_c < -1.96 &
                                           !is.na(z_scores_dataframe_c))
percentage_above_threshold_under_c <- (num_entries_above_threshold_under_c /
                                       total_entries_adjusted_c) * 100
cat("number of under-represented entries:", num_entries_above_threshold_under_c,
    "\n")

## number of under-represented entries: 216

cat("percentage of under-represented entries:",
    percentage_above_threshold_under_c, "%\n")

## percentage of under-represented entries: 35.06494 %

##stages' most over-represented motifs##
#divide Z-scores dataframe into separate dataframes (one per stage)
z_scores_S1_c <- z_scores_dataframe_c[, 1:4]
z_scores_S2_c <- z_scores_dataframe_c[, 5:8]
z_scores_S3_c <- z_scores_dataframe_c[, 9:12]
z_scores_S4_c <- z_scores_dataframe_c[, 13:16]
#calculate mean Z-score accross sites within each stage
z_scores_stages_mean_c <- data.frame(
  Stage1 = rowMeans(z_scores_S1_c, na.rm = TRUE),
  Stage2 = rowMeans(z_scores_S2_c, na.rm = TRUE),
  Stage3 = rowMeans(z_scores_S3_c, na.rm = TRUE),
```

```
  Stage4 = rowMeans(z_scores_S4_c, na.rm = TRUE)
)
#find max value
max_row_indices_c <- apply(z_scores_stages_mean_c, 2, which.max)
#print results
messages_c <- character(4)
for (i in 1:4) {
  column_name_c <- colnames(z_scores_stages_mean_c)[i]
  max_row_name_c <- rownames(z_scores_stages_mean_c)[max_row_indices_c[i]]
  messages_c[i] <- paste("the most over-represented motif of", column_name_c,
                         "is n°", max_row_name_c)
}
cat(messages_c, sep = "\n")

## the most over-represented motif of Stage1 is n° 2

## the most over-represented motif of Stage2 is n° 2

## the most over-represented motif of Stage3 is n° 2

## the most over-represented motif of Stage4 is n° 7

##stages' most under-represented motifs##
#find min value
min_row_indices_2_c <- apply(z_scores_stages_mean_c, 2, which.min)
#print results
messages_2_c <- character(4)
for (i in 1:4) {
  column_name_2_c <- colnames(z_scores_stages_mean_c)[i]
  min_row_name_2_c <- rownames(z_scores_stages_mean_c)[min_row_indices_2_c[i]]
  messages_2_c[i] <- paste("the most under-represented motif of",
                           column_name_2_c, "is n°", min_row_name_2_c)
}
cat(messages_2_c, sep = "\n")

## the most under-represented motif of Stage1 is n° 11

## the most under-represented motif of Stage2 is n° 8

## the most under-represented motif of Stage3 is n° 8

## the most under-represented motif of Stage4 is n° 23

###"vaznull"###
##initialise Z-scores dataframe##
z_scores_dataframe_v <- data.frame(matrix(NA, nrow = 44, ncol = 16))
##randomise networks, calculate motifs' occurrences and calculate Z-scores##
#randomise
set.seed(1)
for (i in seq_along(bipnet_sites_list)) {
  bipnet_null_v <- vaznull(100, bipnet_sites_list[[i]])
  results_matrix_v <- matrix(NA, nrow = 44, ncol = 100)
```

```
  #calculate motifs' occurrences
  for (j in 1:100) {
    mcount_null_v <- mcount(bipnet_null_v[[j]], six_node = TRUE,
                            normalisation = TRUE, mean_weight = TRUE,
                            standard_dev = TRUE)
    normalise_sum_column_v <- mcount_null_v$normalise_sum
    results_matrix_v[, j] <- normalise_sum_column_v
  }
  #calculate mean null model value for each motif
  normalise_sum_null_mean_v <- rowMeans(results_matrix_v, na.rm = TRUE)
  #calculate standard deviation on null model values
  sd_null_mcount_v <- apply(results_matrix_v, 1, sd, na.rm = TRUE)
  #get mcount dataframes' names
  mcount_df_name_v <- paste0("mcount_", names(bipnet_sites_list)[i])
  #calculate Z-scores
  z_scores_v <- (((get(mcount_df_name_v)$normalise_sum) -
                  normalise_sum_null_mean_v) / (sd_null_mcount_v))
  #upload Z-scores in the dataframe
  z_scores_dataframe_v[, i] <- z_scores_v
  #rename columns
  col_name_v <- paste0("z_score_", names(bipnet_sites_list)[i])
  colnames(z_scores_dataframe_v)[i] <- col_name_v
  #replace infinite values with NaNs
  z_scores_dataframe_v[z_scores_dataframe_v == -Inf] <- NaN
  z_scores_dataframe_v[z_scores_dataframe_v == Inf] <- NaN
}
##percentage of entries with |Z|>1.96 across all stages and sites##
num_entries_above_threshold_v <- sum(abs(z_scores_dataframe_v) > 1.96 &
                                       !is.na(z_scores_dataframe_v))
total_entries_adjusted_v <- sum(!is.na(z_scores_dataframe_v))
percentage_above_threshold_v <- (num_entries_above_threshold_v /
                                   total_entries_adjusted_v) * 100
cat("number of entries above threshold:", num_entries_above_threshold_v, "\n")

## number of entries above threshold: 131

cat("percentage of entries above threshold:", percentage_above_threshold_v,
    "%\n")

## percentage of entries above threshold: 22.16582 %

##percentage of entries with Z>1.96 across all stages and sites##
num_entries_above_threshold_over_v <- sum(z_scores_dataframe_v > 1.96 &
                                            !is.na(z_scores_dataframe_v))
percentage_above_threshold_over_v <- (num_entries_above_threshold_over_v /
                                        total_entries_adjusted_v) * 100
cat("number of over-represented entries:", num_entries_above_threshold_over_v,
    "\n")

## number of over-represented entries: 117
```

```r
cat("percentage of over-represented entries:", percentage_above_threshold_over_v
    , "%\n")

## percentage of over-represented entries: 19.79695 %

##percentage of entries with Z<-1.96 across all stages and sites##
num_entries_above_threshold_under_v <- sum(z_scores_dataframe_v < -1.96 &
                                            !is.na(z_scores_dataframe_v))
percentage_above_threshold_under_v <- (num_entries_above_threshold_under_v /
                                        total_entries_adjusted_v) * 100
cat("number of under-represented entries:", num_entries_above_threshold_under_v,
    "\n")

## number of under-represented entries: 14

cat("percentage of under-represented entries:",
    percentage_above_threshold_under_v, "%\n")

## percentage of under-represented entries: 2.368866 %

##stages' most over-represented motifs##
#divide Z-scores dataframe into separate dataframes (one per stage)
z_scores_S1_v <- z_scores_dataframe_v[, 1:4]
z_scores_S2_v <- z_scores_dataframe_v[, 5:8]
z_scores_S3_v <- z_scores_dataframe_v[, 9:12]
z_scores_S4_v <- z_scores_dataframe_v[, 13:16]
#calculate mean Z-score accross sites within each stage
z_scores_stages_mean_v <- data.frame(
  Stage1 = rowMeans(z_scores_S1_v, na.rm = TRUE),
  Stage2 = rowMeans(z_scores_S2_v, na.rm = TRUE),
  Stage3 = rowMeans(z_scores_S3_v, na.rm = TRUE),
  Stage4 = rowMeans(z_scores_S4_v, na.rm = TRUE)
)
#find max value
max_row_indices_v <- apply(z_scores_stages_mean_v, 2, which.max)
#print results
messages_v <- character(4)
for (i in 1:4) {
  column_name_v <- colnames(z_scores_stages_mean_v)[i]
  max_row_name_v <- rownames(z_scores_stages_mean_v)[max_row_indices_v[i]]
  messages_v[i] <- paste("the most over-represented motif of", column_name_v,
                         "is n°", max_row_name_v)
}
cat(messages_v, sep = "\n")

## the most over-represented motif of Stage1 is n° 11

## the most over-represented motif of Stage2 is n° 30

## the most over-represented motif of Stage3 is n° 24
```

```
## the most over-represented motif of Stage4 is n° 36

##stages' most under-represented motifs##
#find min value
min_row_indices_2_v <- apply(z_scores_stages_mean_v, 2, which.min)
#print results
messages_2_v <- character(4)
for (i in 1:4) {
  column_name_2_v <- colnames(z_scores_stages_mean_v)[i]
  min_row_name_2_v <- rownames(z_scores_stages_mean_v)[min_row_indices_2_v[i]]
  messages_2_v[i] <- paste("the most under-represented motif of",
                          column_name_2_v, "is n°", min_row_name_2_v)
}
cat(messages_2_v, sep = "\n")

## the most under-represented motif of Stage1 is n° 38

## the most under-represented motif of Stage2 is n° 38

## the most under-represented motif of Stage3 is n° 29

## the most under-represented motif of Stage4 is n° 26
```

After calculating the frequency of occurrence of motifs in both observed and null model networks, we proceed to perform an NMDS on all possible motifs up to 6 nodes, evaluating via PERMANOVA and multivariate homogeneity of groups' dispersions whether different stages are significantly different in terms of motifs' prevalence.
We perform the NMDS using 'metaMDS' function of 'vegan' package (Oksanen et al. 2022, Oksanen 2023). NMDS is an ordination technique that projects an N–dimensional space of N variables (in our case N=44, i.e. the frequencies of the 44 motifs) associated with n objects (in our case n=16, i.e. sites) into a two-dimensional Cartesian space in which each object finds its specific location based on dissimilarities between pairs of objects. To construct the distance matrix, we considered Bray-Curtis dissimilarities (Bray and Curtis 1957). The resulting two-dimensional space contains n points whose distance is proportional to the dissimilarity among motifs' prevalence (Oksanen et al. 2022, Oksanen 2023). In simpler terms, the closer the sites on the ordination plot, the similar their pattern of motif representation.
We perform PERMANOVA using 'adonis2' function of package 'vegan' (Oksanen et al. 2022). For a given grouping factor ('stage' in this case), PERMANOVA allows us to evaluate the hypothesis that the centroids of the stages are significantly different (Oksanen et al. 2022). We also addressed the issue of multivariate homogeneity of groups' dispersions using function 'betadisper' of 'vegan' package (Oksanen et al. 2022), which is often applied in the context of multivariate analysis of variance (MANOVA) to examine whether groups have similar or different dispersions in multivariate space.

```
###ORDINATION###
##prepare dataframe for ordination##
mcount_list <- lapply(site_names, function(site_name) get(paste("mcount_",
                                                                site_name,
                                                                sep = "")))
```

```r
normalise_sum_columns <- lapply(mcount_list, function(df) df$normalise_sum)
nmds_mcount_data <- data.frame(matrix(unlist(normalise_sum_columns), nrow =
                                        length(site_names), byrow = TRUE))
row.names(nmds_mcount_data) <- site_names
colnames(nmds_mcount_data) <- paste("M", 1:44, sep = "")
##NMDS##
set.seed(1)
nmds_mcount <- metaMDS(nmds_mcount_data, distance = "bray")

## Run 0 stress 0.02290611
## Run 1 stress 0.02290614
## ... Procrustes: rmse 0.01947766  max resid 0.05509088
## Run 2 stress 0.02290618
## ... Procrustes: rmse 0.01947774  max resid 0.05510304
## Run 3 stress 0.03467617
## Run 4 stress 0.0229063
## ... Procrustes: rmse 0.01947953  max resid 0.05520161
## Run 5 stress 0.02290617
## ... Procrustes: rmse 0.01947795  max resid 0.05511081
## Run 6 stress 0.02290624
## ... Procrustes: rmse 0.01947597  max resid 0.05553506
## Run 7 stress 0.03467595
## Run 8 stress 0.02290615
## ... Procrustes: rmse 0.01947548  max resid 0.05548116
## Run 9 stress 0.0229062
## ... Procrustes: rmse 0.01947569  max resid 0.05551819
## Run 10 stress 0.03467596
## Run 11 stress 0.02290738
## ... Procrustes: rmse 0.01948963  max resid 0.05558677
## Run 12 stress 0.03467594
## Run 13 stress 0.03467591
## Run 14 stress 0.02290604
## ... New best solution
## ... Procrustes: rmse 0.01947648  max resid 0.0551822
## Run 15 stress 0.03467603
## Run 16 stress 0.02290613
## ... Procrustes: rmse 0.0001071126  max resid 0.000356022
## ... Similar to previous best
## Run 17 stress 0.03613925
## Run 18 stress 0.03467594
## Run 19 stress 0.03784628
## Run 20 stress 0.02290626
## ... Procrustes: rmse 0.0003686499  max resid 0.001226714
## ... Similar to previous best
## *** Best solution repeated 2 times

#diagnostic plots
plot(nmds_mcount)
```

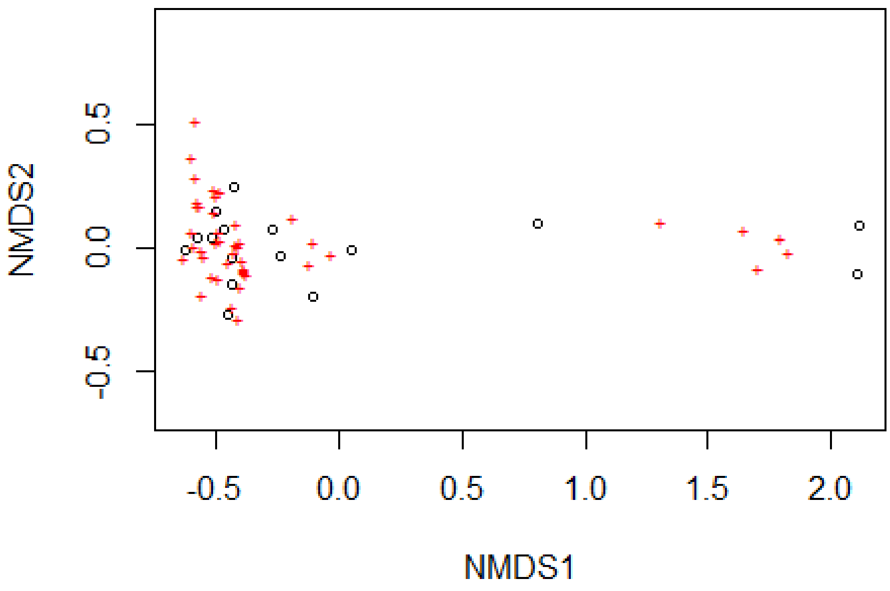

```r
stressplot(nmds_mcount)
```

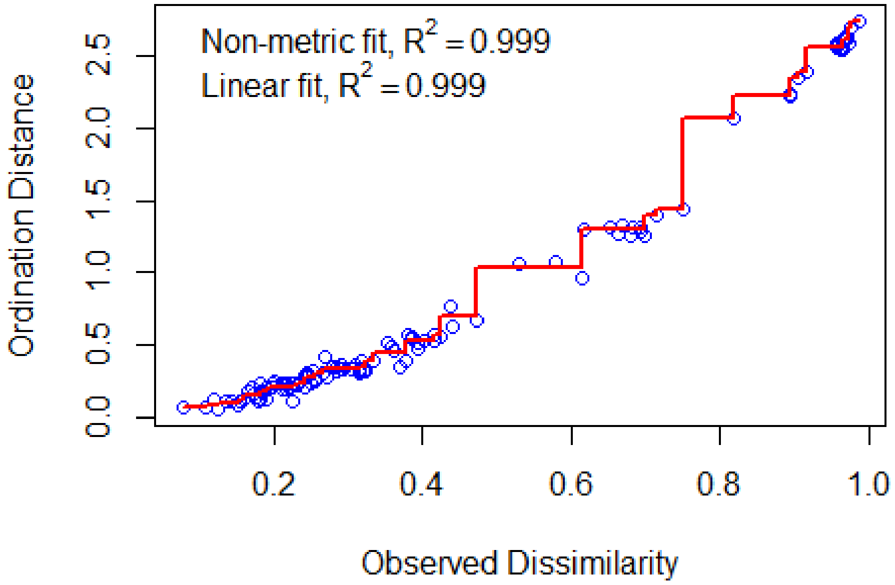

```r
#stress value
nmds_mcount$stress
```

```
## [1] 0.02290604

#scores
scores(nmds_mcount)

## $sites
##          NMDS1        NMDS2
## 1A -0.10769489 -0.196548161
## 1B -0.45300630 -0.269242525
## 1C -0.43977136 -0.148598488
## 1D -0.23750741 -0.030321761
## 2A -0.43622626 -0.039657747
## 2B -0.62633996 -0.010472222
## 2C -0.47457117  0.071519422
## 2D -0.51883750  0.039788032
## 3A -0.43322039  0.246520183
## 3B  0.80833897  0.100616175
## 3C -0.27185868  0.074379380
## 3D -0.57939355  0.038266666
## 4A  2.10913271 -0.106192524
## 4B  2.11226499  0.086481864
## 4C  0.04898201 -0.007335619
## 4D -0.50029120  0.150797326
##
## $species
##           NMDS1        NMDS2
## M1   1.30776751  0.103051052
## M2   1.64808575  0.070619496
## M3  -0.03608272 -0.024061034
## M4  -0.41709970  0.011258291
## M5  -0.12471979 -0.067939051
## M6  -0.38443183 -0.093053263
## M7   1.79037151  0.039164170
## M8  -0.57910898  0.187964007
## M9  -0.43274186 -0.013454653
## M10 -0.45789942 -0.061383345
## M11 -0.50242081  0.024375037
## M12 -0.51272738  0.234298512
## M13 -0.10768350  0.022699179
## M14 -0.41596072 -0.291166207
## M15 -0.38922689 -0.085937901
## M16 -0.39005057 -0.096361617
## M17  1.82297433 -0.021378500
## M18 -0.60105355  0.370053024
## M19 -0.57019387  0.171767287
## M20 -0.59547732  0.009443626
## M21 -0.58063150  0.172367997
## M22 -0.60027666  0.063596115
## M23 -0.58819168  0.289746388
```

```
## M24 -0.58996915  0.521469848
## M25 -0.42506801  0.099787873
## M26 -0.42110573  0.014664822
## M27 -0.49658282 -0.121972635
## M28 -0.55818889 -0.187303647
## M29 -0.49665746  0.062335038
## M30 -0.51737353 -0.116380596
## M31 -0.50174361  0.216458813
## M32 -0.55009286 -0.036437002
## M33 -0.48836526  0.034431332
## M34 -0.63886885 -0.039065927
## M35 -0.56536498 -0.012506870
## M36 -0.51039259  0.145528788
## M37 -0.48823143  0.225854411
## M38 -0.19359899  0.118820481
## M39 -0.43419380 -0.239979581
## M40 -0.40638617  0.023756724
## M41 -0.40114289 -0.155912822
## M42 -0.39475977 -0.049497121
## M43 -0.38392355 -0.104390595
## M44  1.69890404 -0.084598418

#create NMDS scores dataframe
nmds_mcount_scores <- data.frame(scores(nmds_mcount)$sites)
nmds_mcount_scores$Stage <- nmds_mcount_data_2$Stage
#calculate centroids
centroids_mcount <- aggregate(cbind(NMDS1, NMDS2) ~ Stage,
                              data = nmds_mcount_scores, FUN = mean)
##test difference between ordination groups (stages)##
nmds_mcount_data_2 <- nmds_mcount_data
nmds_mcount_data_2$Stage <- c("Stage1","Stage1","Stage1","Stage1","Stage2",
                              "Stage2","Stage2","Stage2", "Stage3", "Stage3",
                              "Stage3", "Stage3", "Stage4", "Stage4","Stage4",
                              "Stage4")
#adonis
set.seed(1)
adonis_mcount <- adonis2(nmds_mcount_data ~ nmds_mcount_data_2$Stage,
                         data = nmds_mcount_data, permutations = 1000,
                         method = "bray")
adonis_mcount

## Permutation test for adonis under reduced model
## Terms added sequentially (first to last)
## Permutation: free
## Number of permutations: 1000
##
## adonis2(formula = nmds_mcount_data ~ nmds_mcount_data_2$Stage, data =
nmds_mcount_data, permutations = 1000, method = "bray")
##                          Df SumOfSqs      R2      F   Pr(>F)
## nmds_mcount_data_2$Stage  3  0.84376 0.37922 2.4435 0.03896 *
```

```
## Residual                  12  1.38125 0.62078
## Total                     15  2.22502 1.00000
## ---
## Signif. codes:  0 '***' 0.001 '**' 0.01 '*' 0.05 '.' 0.1 ' ' 1

distance_matrix_mcount <- vegdist(nmds_mcount_data, method = "bray")
set.seed(1)
betadisper_mcount <- betadisper(distance_matrix_mcount,
                                nmds_mcount_data_2$Stage)
permutest(betadisper_mcount, pairwise = TRUE)

##
## Permutation test for homogeneity of multivariate dispersions
## Permutation: free
## Number of permutations: 999
##
## Response: Distances
##           Df  Sum Sq  Mean Sq      F N.Perm Pr(>F)
## Groups     3 0.29767 0.099222 4.2816    999  0.026 *
## Residuals 12 0.27809 0.023174
## ---
## Signif. codes:  0 '***' 0.001 '**' 0.01 '*' 0.05 '.' 0.1 ' ' 1
##
## Pairwise comparisons:
## (Observed p-value below diagonal, permuted p-value above diagonal)
##           Stage1    Stage2    Stage3 Stage4
## Stage1           0.1240000 0.4720000  0.015
## Stage2 0.1362847           0.2380000  0.003
## Stage3 0.4469513 0.2270974            0.213
## Stage4 0.0233708 0.0099019 0.2174736
```

We conclude this section by drawing the ordination plot.

```
###GRAPH###
figure_2 <- ggplot(nmds_mcount_scores, aes(x = NMDS1, y = NMDS2,
                                           colour = Stage)) +
                   geom_point(data = centroids_mcount, size = 4) +
                   scale_color_manual(values = c("blue3", "green3",
                                                 "yellow3", "red3")) +
                   geom_point() +
                   coord_fixed() +
                   theme_bw() +
                   theme(legend.position = "right",
                         legend.text = element_text(size = 10),
                         legend.direction = 'vertical',
                         panel.grid.major = element_blank(),
                         panel.grid.minor = element_blank()) +
                   scale_y_continuous(expand = c(0.15, 0.15))

figure_2
```

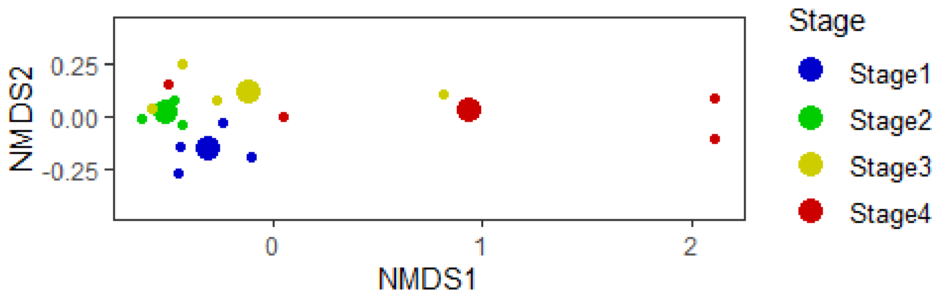

#### (b) Network robustness

In this section, we unfold all analyses concerning network robustness.

First, we import and manipulate the datasheet containing a list of plant species with corresponding Habitat Specificity scores and mean coverage ('Mean_Coverage'). Each species has a Habitat Specificity score for each stage ('HS1', 'HS2', 'HS3', 'HS4'), defined as the mean of the ratio 'n°of occupied sites in a stage'/'total n°of sites in the stage' calculated on both our data and data collected by [Anonymous for DBPR] and [Anonymous for DBPR] in the same sites in 2022. Mean coverage values derive from data collected by [Anonymous for DBPR] and [Anonymous for DBPR] in the same sites in 2022. We assign a successional status ('Type') and an associated extinction probability ('HSp') to plant species based on their set of HS scores following the rules presented below (those are the assumptions of the test model). We then calculate the final extinction probability ('Ext_Prob') as 'Ep'='HSp'/'Mean_Coverage'. Finally, we normalise extinction probabilities within each site.

```
###IMPORT AND MANIPULATE FLORA DATA###
#import
flora_data <- read.csv("flora_data.csv", sep = ";")
#assign succesional status based on habitat specificity across stages
flora_data <- flora_data %>%
  mutate(Type = case_when(
    HS1 >= 0.25 & HS2 < 0.25 & HS3 < 0.25 & HS4 < 0.25 ~ "Early-successional",
```

```
    (HS1 >= 0.25 & HS2 >= 0.25 & HS3 < 0.25 & HS4 < 0.25) |
      (HS1 >= 0.25 & HS3 >= 0.25 & HS2 < 0.25 & HS4 < 0.25) ~
      "Early to mid-successional",
    (HS2 >= 0.25 & HS1 < 0.25 & HS3 < 0.25 & HS4 < 0.25) |
      (HS3 >= 0.25 & HS1 < 0.25 & HS2 < 0.25 & HS4 < 0.25) |
        (HS2 >= 0.25 & HS3 >= 0.25 & HS1 < 0.25 & HS4 < 0.25)  ~
      "Mid-successional",
    HS1 >= 0.25 & HS2 >= 0.25 & HS3 >= 0.25 & HS4 < 0.25 ~ "Early-ubiquitous",
    HS4 >= 0.25 & HS1 < 0.25 & HS2 < 0.25 & HS3 < 0.25 ~ "Late-successional",
    (HS2 >= 0.25 & HS4 >= 0.25 & HS1 < 0.25 & HS3 < 0.25) |
      (HS3 >= 0.25 & HS4 >= 0.25 & HS1 < 0.25 & HS2 < 0.25) ~
      "Mid to late-successional",
    HS2 >= 0.25 & HS3 >= 0.25 & HS4 >= 0.25 & HS1 < 0.25 ~ "Late-ubiquitous",
    (HS1 >= 0.25 & HS2 >= 0.25 & HS3 >= 0.25 & HS4 >= 0.25) |
      (HS1 >= 0.25 & HS2 >= 0.25 & HS4 >= 0.25 & HS3 < 0.25) |
        (HS1 >= 0.25 & HS3 >= 0.25 & HS4 >= 0.25 & HS2 < 0.25) ~ "Ubiquitous",
    HS1 < 0.25 & HS2 < 0.25 & HS3 < 0.25 & HS4 < 0.25 ~ "Rare",
    TRUE ~ NA_character_
  ))
#assign extinction probabilities to each successional status (HSp)
flora_data <- flora_data %>%
  mutate(HSp = case_when(
    Type == "Rare" ~ 0.9,
    Type == "Early-successional" ~ 0.8,
    Type == "Early to mid-successional" ~ 0.7,
    Type == "Mid-successional" ~ 0.6,
    Type == "Early-ubiquitous" ~ 0.5,
    Type == "Late-successional" ~ 0.4,
    Type == "Mid to late-successional" ~ 0.3,
    Type == "Late-ubiquitous" ~ 0.2,
    Type == "Ubiquitous" ~ 0.1,
    TRUE ~ NA_real_
  ))

###PREPARE MATRICES AND DATAFRAMES FOR EXTINCTION CASCADES###
##convert bipnet_sites_list elements into separate matrices##
for (site in site_names) {
  assign(paste("bipnet_", site, sep = ""), bipnet_sites_list[[site]])
}
##calculate extinction probabilities##
flora_data$Ext_Prob <- flora_data$HSp / flora_data$Mean_Coverage
##normalise extinction probabilities within each site##
for (site in site_names) {
  #get species names from bipartite network
  site_species <- rownames(bipnet_sites_list[[site]])
  #filter flora_data for matching species
  matching_species <- flora_data$Species %in% site_species
  ext_prob_df <- data.frame(
    Species_F = flora_data$Species[matching_species],
    Ext_Probs = flora_data$Ext_Prob[matching_species],
```

```
    stringsAsFactors = FALSE
  )
  #scale probabilities to 1
  ext_prob_df$Norm_Probs <- ext_prob_df$Ext_Probs / sum(ext_prob_df$Ext_Probs)
  #assign the dataframe to the corresponding site
  assign(paste("ext_prob_", site, sep = ""), ext_prob_df)
}
```

We proceed to build the function to calculate robustness in two version: the first one extracts plant species with a probabilistic approach (test model), the second one extracts plant species randomly (null model). The robustness function consists of three different functions that are called sequentially: I) 'second_ext_function', II) 'scale_coord' and III)'robustness_calc'.
I)'second_ext_function' takes a pollination matrix (matrix_obj) with plant species as rows and pollinator families as columuns and performs a secondary extinction cascade exterminating plant species based on their extinction probabilities with a probabilistic approach (the highest the probability, the highest the chance of the species being sampled by the function). Extinction probabilities are provided by a dataframe (df_obj) containing: i) the same list of plant species of the matrix, as the column named "Species_F", ii) the normalised extinction probabilities of each species, as the column named "Norm_Probs". The function can of course operate with different kinds of interaction matrices and probabilities.
II)'scale_coord' takes a dataframe and scales to 1 the values in columns "Iteration" and "Remaining_Poll_Families", creating new columns "Iteration_Fraction" and "Remaining_Poll_Families_Fraction".
III)'robustness_calc' takes a dataframe, interpolates a function on coordinates "Iteration_Fraction" (x) and "Remaining_Poll_Families" (y) and calculates the area under the curve as its definite integral from 0 to 1.

```
###ROBUSTNESS FUNCTION###
##test model##
robustness_function <- function(matrix_obj, df_obj) {
  second_ext_function <- function(matrix_obj, df_obj) {
    #initialise a dataframe to keep track of iterations
    ext_table <- data.frame(Iteration = integer(0),
                            Removed_Plant_Species = character(0),
                            Remaining_Poll_Families = integer(0))
    iteration <- 0
    #continue until all pollinator families are extinct
    while (sum(colSums(matrix_obj) > 0) > 0) {
      iteration <- iteration + 1
      #filter out plant species with extinction probability equal to 0 (i.e.
already selected)
      non_zero_probs_df <- df_obj[df_obj$Norm_Probs > 0, ]
      #store extinction probabilities based on the filtered dataframe
      ext_probs <- non_zero_probs_df$Norm_Probs
      if (length(ext_probs) == 0) {
        break  #if there are no plant species left in df_obj with non-zero
probabilities, exit the loop
      }
      #select a plant species in df_obj based on extinction probabilities
```

```
      selected_plant <- sample(non_zero_probs_df$Species_F, size = 1,
                               prob = ext_probs)
      #find the row index of the selected plant species in matrix_obj
      plant_index <- which(rownames(matrix_obj) == selected_plant)
      #update to zero the selected plant species corresponding entries in
matrix_obj (rows)
      matrix_obj[plant_index, ] <- 0
      #update to zero the pollinator families without any links left in matrix_obj
(columns)
      matrix_obj[, colSums(matrix_obj) == 0] <- 0
      #update the extinction probabilities of already selected plants to zero in
df_obj
      df_obj$Norm_Probs[df_obj$Species_F == selected_plant] <- 0
      #update the extinction table
      remaining_families <- sum(colSums(matrix_obj) > 0)
      ext_table <- rbind(ext_table, data.frame(Iteration = iteration,
                                               Removed_Plant_Species =
                                                 selected_plant,
                                               Remaining_Poll_Families =
                                                 remaining_families))
    }
    return(ext_table)
  }
  scale_coord <- function(df_obj_2) {
    #ensure the dataframe has 'Iteration' and 'Remaining_Poll_Families' columns
    if (!all(c("Iteration", "Remaining_Poll_Families") %in% colnames(df_obj_2)))
      {
      stop("Dataframe must have 'Iteration' and 'Remaining_Poll_Families'
           columns.")
    }
    #calculate the max
    max_Iteration <- max(df_obj_2$Iteration)
    max_Poll_Families <- ncol(matrix_obj)
    #scale
    df_obj_2$Iteration_Fraction <- df_obj_2$Iteration / max_Iteration
    df_obj_2$Remaining_Poll_Families_Fraction <-
      df_obj_2$Remaining_Poll_Families / max_Poll_Families
    initial_conditions <- data.frame(Iteration = 0,
                                     Remaining_Poll_Families =
                                       max_Poll_Families,
                                     Removed_Plant_Species = "None",
                                     Iteration_Fraction = 0,
                                     Remaining_Poll_Families_Fraction = 1)
    df_obj_2 <- rbind(df_obj_2, initial_conditions)
    return(df_obj_2)
  }
  robustness_calc <- function(df_obj_3) {
    #create an interpolated function based on ATC coordinates
    interp_func <- approxfun(df_obj_3$Iteration_Fraction,
                             df_obj_3$Remaining_Poll_Families_Fraction)
    #integrate the function
```

```r
    robustness_result <- integrate(interp_func,
                                   lower = min(df_obj_3$Iteration_Fraction),
                                   upper = max(df_obj_3$Iteration_Fraction))
    #print the result
    return(robustness_result[["value"]])
  }
  #call the functions sequentially
  df_obj_2 <- second_ext_function(matrix_obj, df_obj)
  df_obj_3 <- scale_coord(df_obj_2)
  print(df_obj_3)
  robustness_calc(df_obj_3)
}
##null model##
robustness_function_null <- function(matrix_obj, df_obj) {
  second_ext_function_null <- function(matrix_obj, df_obj) {
    ext_table <- data.frame(Iteration = integer(0),
                            Removed_Plant_Species = character(0),
                            Remaining_Poll_Families = integer(0))
    iteration <- 0
    while (sum(colSums(matrix_obj) > 0) > 0) {
      iteration <- iteration + 1
      non_zero_probs_df <- df_obj[df_obj$Norm_Probs > 0, ]
      ext_probs <- non_zero_probs_df$Norm_Probs
      if (length(ext_probs) == 0) {
        break
      }
      selected_plant <- sample(non_zero_probs_df$Species_F, size = 1)
      plant_index <- which(rownames(matrix_obj) == selected_plant)
      matrix_obj[plant_index, ] <- 0
      matrix_obj[, colSums(matrix_obj) == 0] <- 0
      df_obj$Norm_Probs[df_obj$Species_F == selected_plant] <- 0
      remaining_families <- sum(colSums(matrix_obj) > 0)
      ext_table <- rbind(ext_table, data.frame(Iteration = iteration,
                                               Removed_Plant_Species =
                                                 selected_plant,
                                               Remaining_Poll_Families =
                                                 remaining_families))
    }
    print(ext_table)
  }
  scale_coord <- function(df_obj_2) {
    if (!all(c("Iteration", "Remaining_Poll_Families") %in% colnames(df_obj_2)))
      {
      stop("Dataframe must have 'Iteration' and 'Remaining_Poll_Families'
           columns.")
    }
    max_Iteration <- max(df_obj_2$Iteration)
    max_Poll_Families <- ncol(matrix_obj)
    df_obj_2$Iteration_Fraction <- df_obj_2$Iteration / max_Iteration
    df_obj_2$Remaining_Poll_Families_Fraction <-
      df_obj_2$Remaining_Poll_Families / max_Poll_Families
```

```
    initial_conditions <- data.frame(Iteration = 0,
                                     Remaining_Poll_Families =
                                       max_Poll_Families,
                                     Removed_Plant_Species = "None",
                                     Iteration_Fraction = 0,
                                     Remaining_Poll_Families_Fraction = 1)
    df_obj_2 <- rbind(df_obj_2, initial_conditions)
    return(df_obj_2)
  }
  robustness_calc <- function(df_obj_3) {
    interp_func <- approxfun(df_obj_3$Iteration_Fraction,
                             df_obj_3$Remaining_Poll_Families_Fraction)
    robustness_result <- integrate(interp_func,
                                   lower = min(df_obj_3$Iteration_Fraction),
                                   upper = max(df_obj_3$Iteration_Fraction))
    return(robustness_result[["value"]])
  }
  df_obj_2 <- second_ext_function_null(matrix_obj, df_obj)
  df_obj_3 <- scale_coord(df_obj_2)
  print(df_obj_3)
  robustness_calc(df_obj_3)
}
```

We calculate robustness in each site for both the test model and the null model as the mean of 1’000 iterations produced by the robustness function. We then calculate two-sample Z-scores and compute Mann-Whitney tests to evaluate if the test model and the null model are significantly different.

Two-sample Z-score is defined as $\boldsymbol{Z} = \frac{\bar{x}_1 - \bar{x}_2}{\sqrt{\left({\sigma_1}^2/n_1\right)+\left({\sigma_2}^2/n_2\right)}}$, where $\bar{\boldsymbol{x}}_1$ and $\bar{\boldsymbol{x}}_2$ are the mean robustness values under the test model and the null model, $\boldsymbol{\sigma}_1$ and $\boldsymbol{\sigma}_2$ are the standard deviation values under the test model and the null model, $\boldsymbol{n}_1$ and $\boldsymbol{n}_2$ are population dimensions (i.e. 1’000). Mann-Whitney test was performed using ‘wilcox.test’ function of ‘stats’ package (R Core Team 2023).

```
###ROBUSTNESS CALCULATION###
#initialise a new dataframe to store robustness results
robustness_data <- data.frame(Plot = character(0), Robustness = numeric(0),
                              Robustness_Null = numeric(0),
                              Z_Scores = numeric(0), p_Wilcox = numeric(0),
                              stringsAsFactors = FALSE)
#loop across sites
for (site in site_names) {
  set.seed(1)
  #provide the interaction matrix and the extinction probabilities dataframe
  matrix_obj <- get(paste("bipnet_", site, sep = ""))
  df_obj <- get(paste("ext_prob_", site, sep = ""))
  #initialise vectors to store robustness results
  robustness_values <- numeric(1000)
  robustness_values_null <- numeric(1000)
  #loop the function 1000 times for test model
```

```
  for (i in 1:1000) {
    robustness_values[i] <- robustness_function(matrix_obj, df_obj)
  }
  #calculate the mean of 1000 iterations for test model
  mean_robustness <- mean(robustness_values)
  #loop the function 1000 times for null model
  for (i in 1:1000) {
    robustness_values_null[i] <- robustness_function_null(matrix_obj, df_obj)
  }
  #calculate the mean of 1000 iterations for null model
  mean_robustness_null <- mean(robustness_values_null)
  #calculate standard deviations
  sd_value <- sd(robustness_values)
  sd_value_null <- sd(robustness_values_null)
  #calculate Z-score
  z_score <- ((mean_robustness - mean_robustness_null) / sqrt(((sd_value^2) /
                                          1000) + ((sd_value_null^2) / 1000)))
  #perform Mann-Whitney test
  wilcox_result <- wilcox.test(robustness_values, robustness_values_null)
  #extract p-value
  p_wilcox <- wilcox_result$p.value
  #store the results in the dataframe
  robustness_data <- rbind(robustness_data, data.frame(Plot = site,
                                                       Robustness =
                                                         mean_robustness,
                                                       Robustness_Null =
                                                         mean_robustness_null,
                                                       Z_Scores = z_score,
                                                       p_Wilcox = p_wilcox))
}
```

We model test model robustness, null model robustness and Z-scores in response to time since deglaciation and NMD1 coordinates of motifs' ordination.

```
###ROBUSTNESS~TIME+NMDS1###
##add stages and time since deglaciation columns to robustness_data##
robustness_data$Stage <- nmds_mcount_data_2$Stage
robustness_data$Time <- as.numeric(c("17", "17", "17", "17", "66", "66", "66",
                                     "66", "111", "111", "111", "111", "141",
                                     "141", "141", "141"))
robustness_data$NMDS1 <- nmds_mcount_scores$NMDS1
##linear models##
lm_robustness_1 <- lm(Robustness ~ as.factor(Time) + NMDS1,
                      data = robustness_data)
summary(lm_robustness_1)

##
## Call:
## lm(formula = Robustness ~ as.factor(Time) + NMDS1, data = robustness_data)
##
## Residuals:
```

```
##       Min        1Q    Median        3Q       Max 
## -0.099273 -0.023012 -0.005014  0.029147  0.109301 
## 
## Coefficients:
##                     Estimate Std. Error t value Pr(>|t|)    
## (Intercept)         0.638374   0.032616  19.572 6.74e-10 ***
## as.factor(Time)66   0.079124   0.045169   1.752   0.1076    
## as.factor(Time)111  0.064814   0.045133   1.436   0.1788    
## as.factor(Time)141 -0.003113   0.054137  -0.057   0.9552    
## NMDS1              -0.061708   0.024160  -2.554   0.0268 *  
## ---
## Signif. codes:  0 '***' 0.001 '**' 0.01 '*' 0.05 '.' 0.1 ' ' 1
## 
## Residual standard error: 0.0635 on 11 degrees of freedom
## Multiple R-squared:  0.6769, Adjusted R-squared:  0.5594 
## F-statistic: 5.762 on 4 and 11 DF,  p-value: 0.00945
```

```
lm_robustness_2 <- lm(Robustness ~ Time + NMDS1, data = robustness_data)
summary(lm_robustness_2)
```

```
## 
## Call:
## lm(formula = Robustness ~ Time + NMDS1, data = robustness_data)
## 
## Residuals:
##       Min        1Q    Median        3Q       Max 
## -0.149972 -0.029755  0.002276  0.041127  0.124939 
## 
## Coefficients:
##               Estimate Std. Error t value Pr(>|t|)    
## (Intercept)  0.6537844  0.0393142  16.630 3.84e-10 ***
## Time         0.0002364  0.0004222   0.560  0.58513    
## NMDS1       -0.0852407  0.0228968  -3.723  0.00256 ** 
## ---
## Signif. codes:  0 '***' 0.001 '**' 0.01 '*' 0.05 '.' 0.1 ' ' 1
## 
## Residual standard error: 0.06872 on 13 degrees of freedom
## Multiple R-squared:  0.5527, Adjusted R-squared:  0.4839 
## F-statistic: 8.032 on 2 and 13 DF,  p-value: 0.005355
```

```
lm_robustness_3 <- lm(Robustness ~  poly(Time, 2) + NMDS1,
                      data = robustness_data)
summary(lm_robustness_3)
```

```
## 
## Call:
## lm(formula = Robustness ~ poly(Time, 2) + NMDS1, data = robustness_data)
## 
## Residuals:
##      Min       1Q   Median       3Q      Max 
## -0.10095 -0.02145 -0.00591  0.02814  0.10742 
```

```
##
## Coefficients:
##                  Estimate Std. Error t value Pr(>|t|)
## (Intercept)      0.673580   0.015209  44.288 1.14e-14 ***
## poly(Time, 2)1  0.004565   0.072505   0.063   0.9508
## poly(Time, 2)2 -0.147694   0.068946  -2.142   0.0534 .
## NMDS1           -0.062087   0.022971  -2.703   0.0192 *
## ---
## Signif. codes:  0 '***' 0.001 '**' 0.01 '*' 0.05 '.' 0.1 ' ' 1
##
## Residual standard error: 0.06084 on 12 degrees of freedom
## Multiple R-squared:  0.6764, Adjusted R-squared:  0.5956
## F-statistic: 8.363 on 3 and 12 DF,  p-value: 0.002859

#model selection
aic_lm_robustness <- AIC(lm_robustness_1, lm_robustness_2, lm_robustness_3)
aic_lm_robustness

##                  df       AIC
## lm_robustness_1  6 -36.80624
## lm_robustness_2  4 -35.60164
## lm_robustness_3  5 -38.78284

#diagnostic
shapiro.test(lm_robustness_3$residuals)

##
##  Shapiro-Wilk normality test
##
## data:  lm_robustness_3$residuals
## W = 0.9537, p-value = 0.5504

plot(lm_robustness_3
```

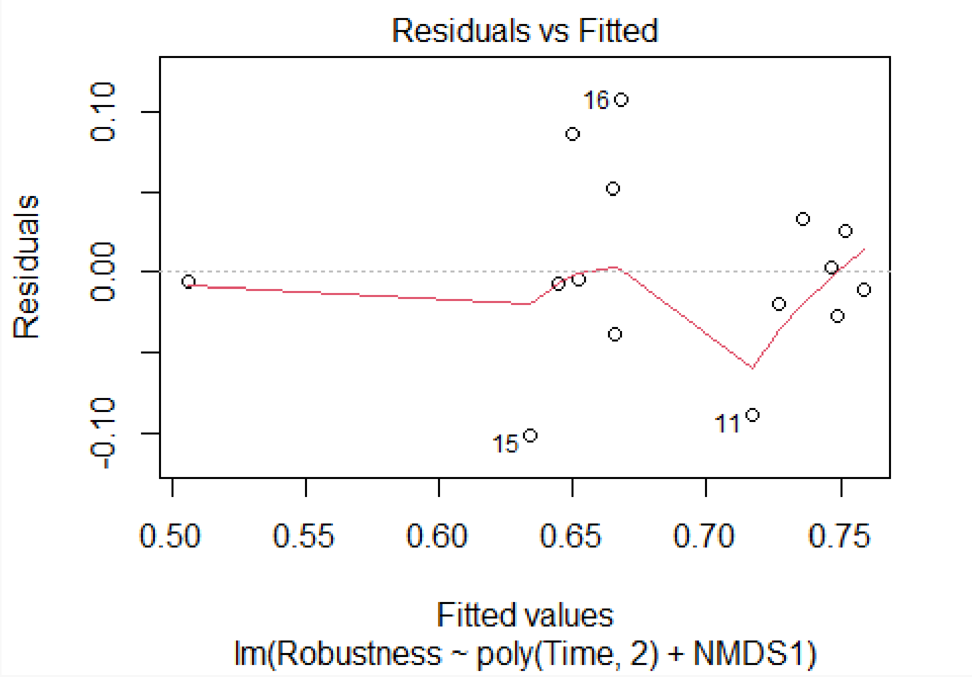
Residuals vs Fitted
Residuals
0.10
0.00
-0.10
16
15
11
0.50
0.55
0.60
0.65
0.70
0.75
Fitted values
lm(Robustness ~ poly(Time, 2) + NMDS1)

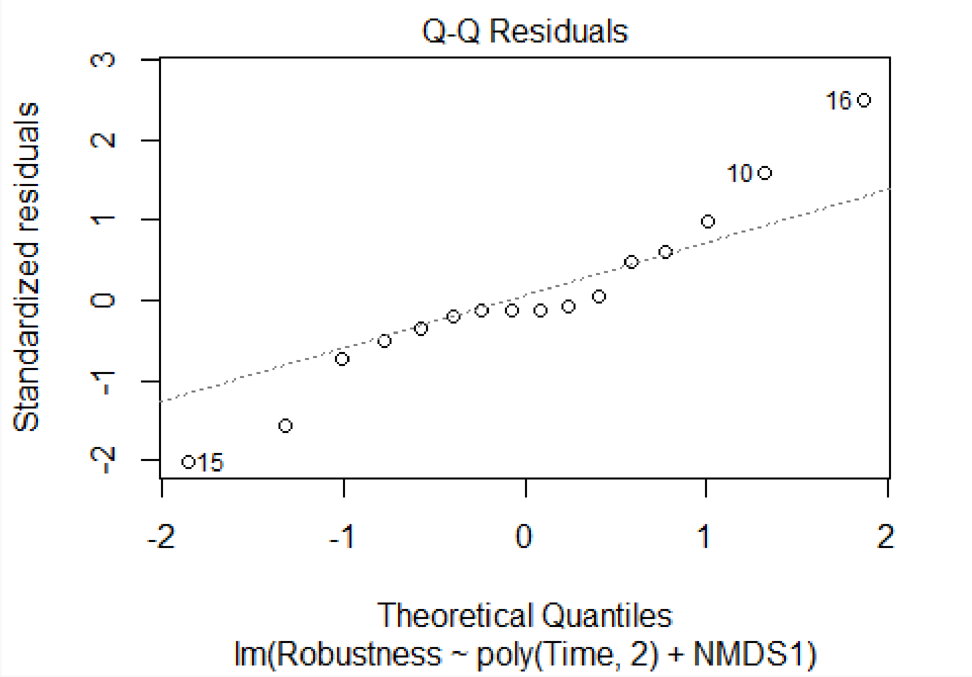
Q-Q Residuals
Standardized residuals
3
2
1
0
-1
-2
16
10
15
-2
-1
0
1
2
Theoretical Quantiles
lm(Robustness ~ poly(Time, 2) + NMDS1)

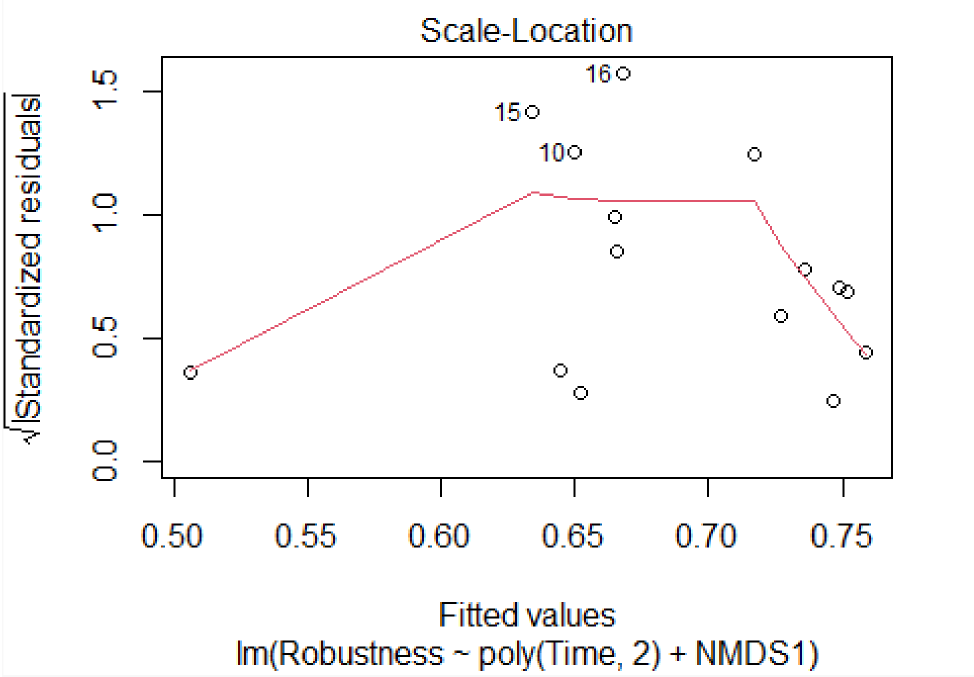

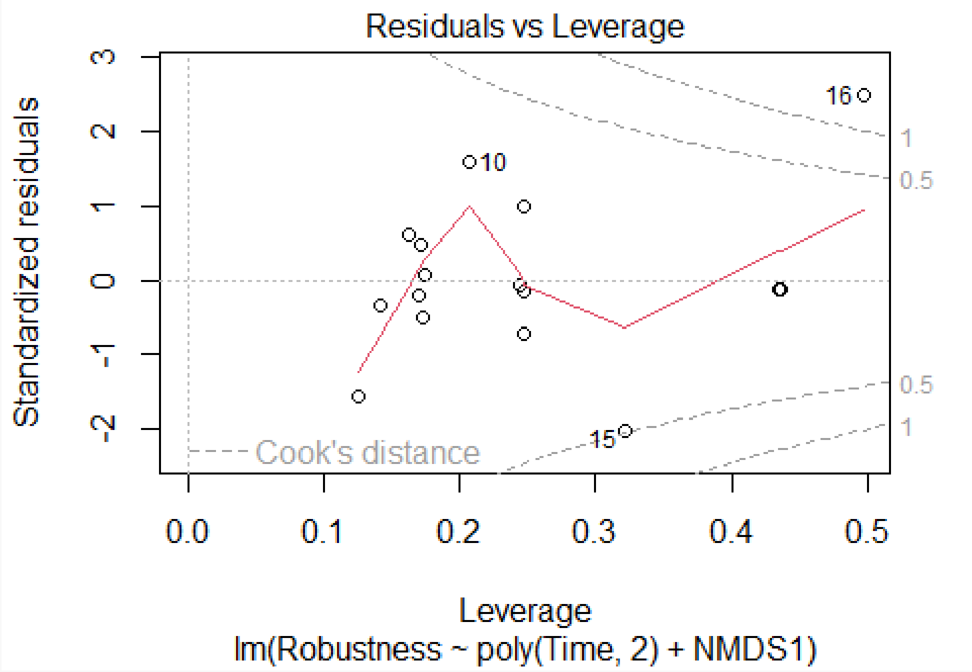

```
##beta regression models##
betareg_robustness_1 <- betareg(Robustness ~ as.factor(Time) + NMDS1,
                                data = robustness_data)
summary(betareg_robustness_1)
```

```
##
## Call:
## betareg(formula = Robustness ~ as.factor(Time) + NMDS1, data = robustness_data)
##
## Standardized weighted residuals 2:
##     Min      1Q  Median      3Q     Max
## -2.3952 -0.5645 -0.1101  0.6951  3.0981
##
## Coefficients (mean model with logit link):
##                    Estimate Std. Error z value Pr(>|z|)
## (Intercept)         0.56357    0.12109   4.654 3.25e-06 ***
## as.factor(Time)66   0.37814    0.17533   2.157  0.03102 *
## as.factor(Time)111  0.31123    0.17206   1.809  0.07047 .
## as.factor(Time)141  0.02276    0.20158   0.113  0.91009
## NMDS1              -0.27006    0.08857  -3.049  0.00229 **
##
## Phi coefficients (precision model with identity link):
##       Estimate Std. Error z value Pr(>|z|)
## (phi)    78.47      27.59   2.844  0.00445 **
## ---
## Signif. codes:  0 '***' 0.001 '**' 0.01 '*' 0.05 '.' 0.1 ' ' 1
##
## Type of estimator: ML (maximum likelihood)
## Log-likelihood: 24.71 on 6 Df
## Pseudo R-squared: 0.6551
## Number of iterations: 43 (BFGS) + 3 (Fisher scoring)
```

```r
betareg_robustness_2 <- betareg(Robustness ~ Time + NMDS1,
                                data = robustness_data)
summary(betareg_robustness_2)
```

```
##
## Call:
## betareg(formula = Robustness ~ Time + NMDS1, data = robustness_data)
##
## Standardized weighted residuals 2:
##     Min      1Q  Median      3Q     Max
## -2.5865 -0.5537 -0.0018  0.7280  2.0987
##
## Coefficients (mean model with logit link):
##              Estimate Std. Error z value Pr(>|z|)
## (Intercept)  0.628509   0.161761   3.885 0.000102 ***
## Time         0.001288   0.001754   0.734 0.462816
## NMDS1       -0.368245   0.091409  -4.029 5.61e-05 ***
##
## Phi coefficients (precision model with identity link):
##       Estimate Std. Error z value Pr(>|z|)
## (phi)    56.92      19.97    2.85  0.00437 **
## ---
## Signif. codes:  0 '***' 0.001 '**' 0.01 '*' 0.05 '.' 0.1 ' ' 1
##
```

```
## Type of estimator: ML (maximum likelihood)
## Log-likelihood: 22.15 on 4 Df
## Pseudo R-squared: 0.5308
## Number of iterations: 32 (BFGS) + 3 (Fisher scoring)

betareg_robustness_3 <- betareg(Robustness ~ poly(Time, 2) + NMDS1,
                                data = robustness_data)
summary(betareg_robustness_3)

##
## Call:
## betareg(formula = Robustness ~ poly(Time, 2) + NMDS1, data = robustness_data)
##
## Standardized weighted residuals 2:
##     Min      1Q  Median      3Q     Max
## -2.3395 -0.5146 -0.1114  0.6463  2.9358
##
## Coefficients (mean model with logit link):
##                 Estimate Std. Error z value Pr(>|z|)
## (Intercept)      0.74163    0.06085  12.188  < 2e-16 ***
## poly(Time, 2)1   0.06561    0.28474   0.230  0.81777
## poly(Time, 2)2  -0.67069    0.27631  -2.427  0.01521 *
## NMDS1           -0.27054    0.08800  -3.074  0.00211 **
##
## Phi coefficients (precision model with identity link):
##       Estimate Std. Error z value Pr(>|z|)
## (phi)    78.45      27.58   2.844  0.00445 **
## ---
## Signif. codes:  0 '***' 0.001 '**' 0.01 '*' 0.05 '.' 0.1 ' ' 1
##
## Type of estimator: ML (maximum likelihood)
## Log-likelihood: 24.71 on 5 Df
## Pseudo R-squared: 0.6552
## Number of iterations: 95 (BFGS) + 2 (Fisher scoring)

#model selection
aic_betareg_robustness <- AIC(betareg_robustness_1, betareg_robustness_2,
                              betareg_robustness_3)
aic_betareg_robustness

##                      df       AIC
## betareg_robustness_1  6 -37.42081
## betareg_robustness_2  4 -36.30713
## betareg_robustness_3  5 -39.41682

#diagnostic
shapiro.test(betareg_robustness_3$residuals)

##
##  Shapiro-Wilk normality test
##
```

```
## data:  betareg_robustness_3$residuals
## W = 0.95406, p-value = 0.5566
```

```
plot(betareg_robustness_3)
```

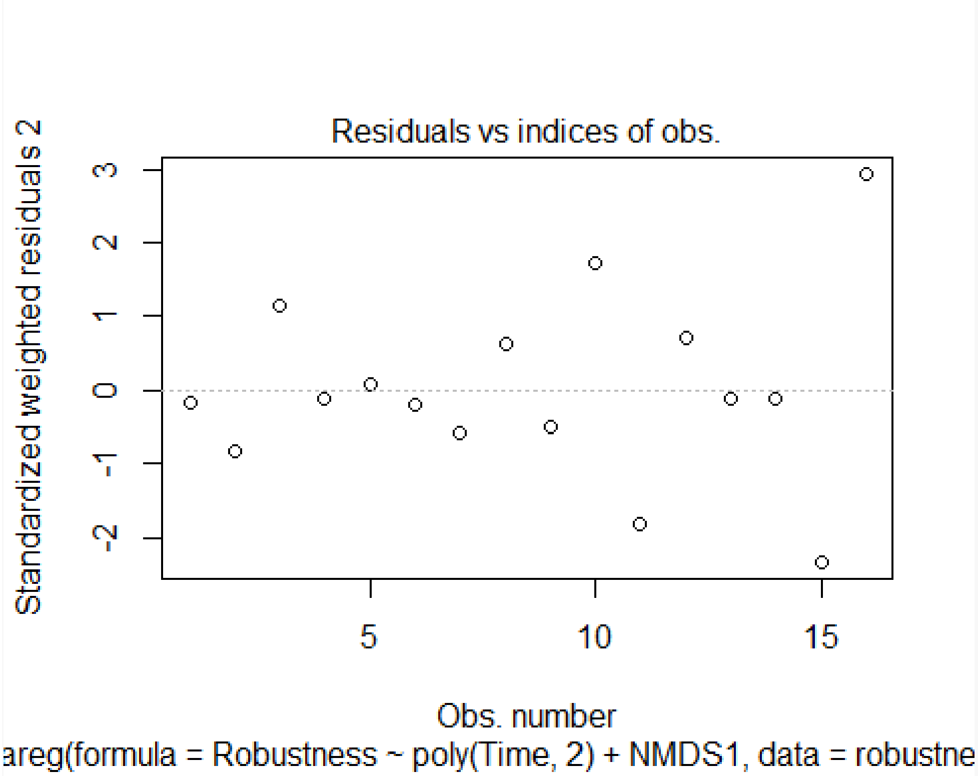

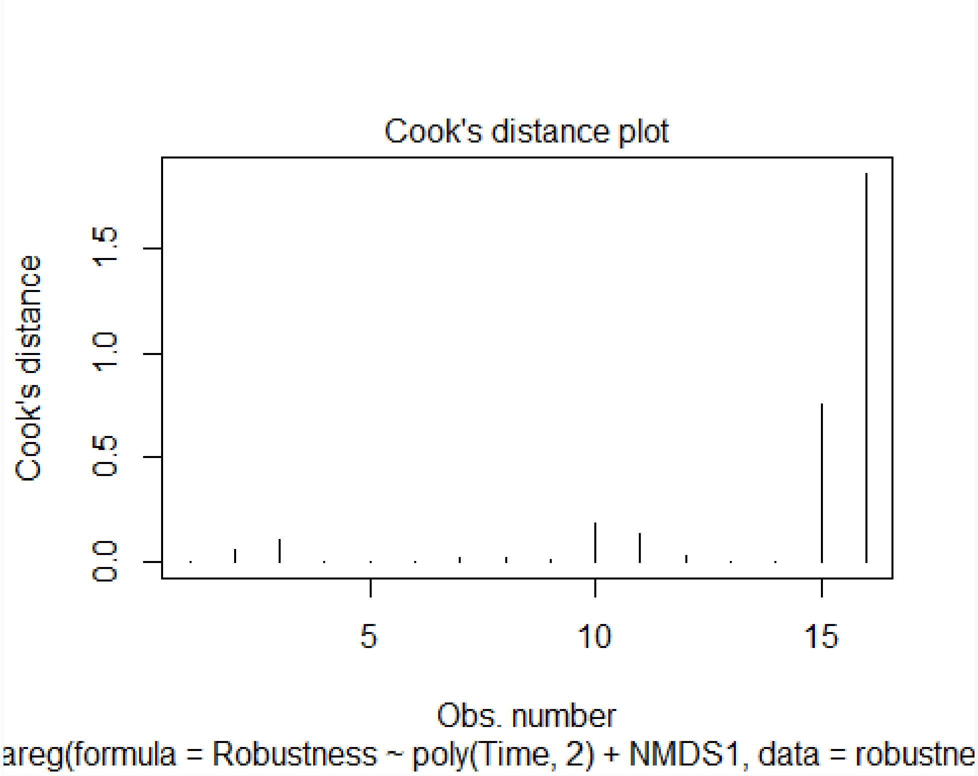

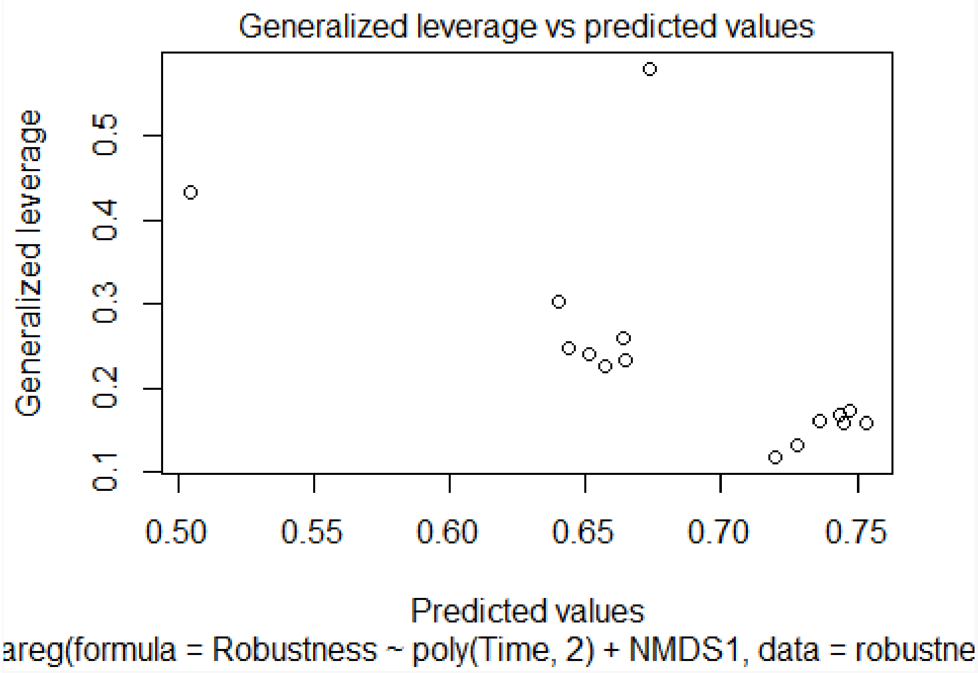

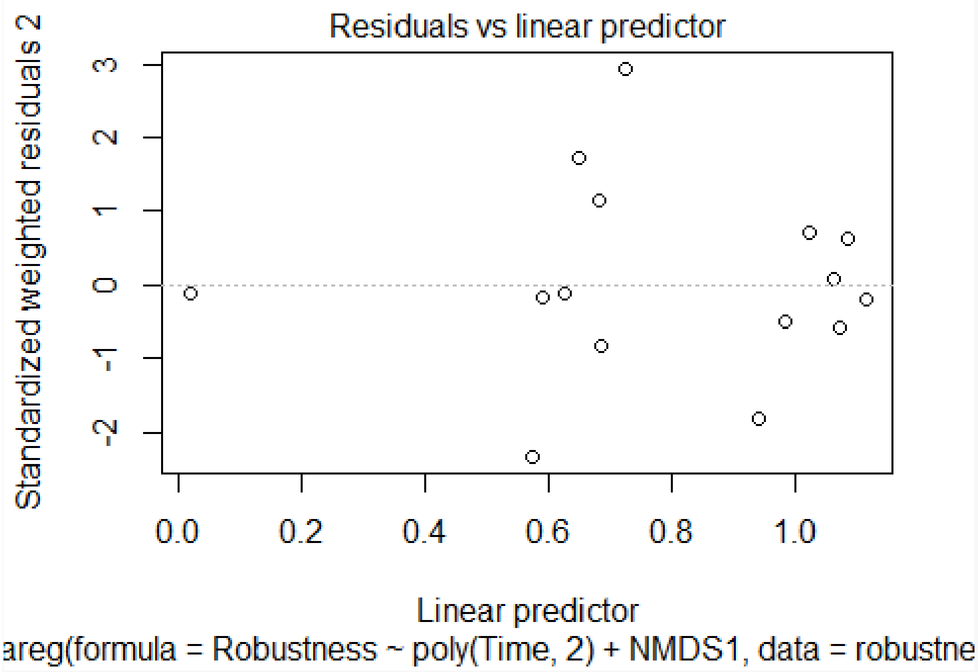

```
#Type-II ANOVA
anovaII_robustness <- Anova(betareg_robustness_3, type = "II")
anovaII_robustness
```

```
## Analysis of Deviance Table (Type II tests)
##
## Response: Robustness
##                Df  Chisq Pr(>Chisq)
## poly(Time, 2)  2 6.6874   0.035306 *
## NMDS1          1 9.4519   0.002109 **
## ---
## Signif. codes:  0 '***' 0.001 '**' 0.01 '*' 0.05 '.' 0.1 ' ' 1
```

```
#collinearity
time_NMDS1_coll <- lm(Time ~ NMDS1, data = robustness_data)
summary(time_NMDS1_coll)
```

```
##
## Call:
## lm(formula = Time ~ NMDS1, data = robustness_data)
##
## Residuals:
##     Min      1Q  Median      3Q     Max
## -63.850 -18.140  -0.257  35.657  70.722
##
## Coefficients:
##             Estimate Std. Error t value Pr(>|t|)
## (Intercept)    83.75      10.88   7.701 2.13e-06 ***
## NMDS1          26.93      12.58   2.141   0.0504 .
## ---
## Signif. codes:  0 '***' 0.001 '**' 0.01 '*' 0.05 '.' 0.1 ' ' 1
##
## Residual standard error: 43.5 on 14 degrees of freedom
## Multiple R-squared:  0.2466, Adjusted R-squared:  0.1928
## F-statistic: 4.582 on 1 and 14 DF,  p-value: 0.05039
```

```
###ROBUSTNESS_NULL~TIME+NMDS1###
##linear models##
lm_robustness_null_1 <- lm(Robustness_Null ~ as.factor(Time) + NMDS1,
                           data = robustness_data)
summary(lm_robustness_null_1)
```

```
##
## Call:
## lm(formula = Robustness_Null ~ as.factor(Time) + NMDS1, data = robustness_data)
##
## Residuals:
##       Min        1Q    Median        3Q       Max
## -0.043354 -0.008781  0.006971  0.012804  0.031579
##
## Coefficients:
##                    Estimate Std. Error t value Pr(>|t|)
## (Intercept)        0.606363   0.012989  46.684 5.33e-14 ***
## as.factor(Time)66  0.023114   0.017988   1.285 0.225185
## as.factor(Time)111 0.033477   0.017973   1.863 0.089418 .
```

```
## as.factor(Time)141 -0.009297   0.021559  -0.431 0.674639
## NMDS1               -0.049652   0.009621  -5.161 0.000313 ***
## ---
## Signif. codes:  0 '***' 0.001 '**' 0.01 '*' 0.05 '.' 0.1 ' ' 1
##
## Residual standard error: 0.02529 on 11 degrees of freedom
## Multiple R-squared:  0.8622, Adjusted R-squared:  0.812
## F-statistic:  17.2 on 4 and 11 DF,  p-value: 0.0001061
```

```
lm_robustness_null_2 <- lm(Robustness_Null ~ Time + NMDS1,
                           data = robustness_data)
summary(lm_robustness_null_2)
```

```
##
## Call:
## lm(formula = Robustness_Null ~ Time + NMDS1, data = robustness_data)
##
## Residuals:
##       Min        1Q    Median        3Q       Max
## -0.047815 -0.005799  0.004013  0.010406  0.043126
##
## Coefficients:
##               Estimate Std. Error t value Pr(>|t|)
## (Intercept)  6.102e-01  1.642e-02  37.164 1.38e-14 ***
## Time         9.557e-05  1.763e-04   0.542    0.597
## NMDS1       -6.046e-02  9.562e-03  -6.323 2.65e-05 ***
## ---
## Signif. codes:  0 '***' 0.001 '**' 0.01 '*' 0.05 '.' 0.1 ' ' 1
##
## Residual standard error: 0.0287 on 13 degrees of freedom
## Multiple R-squared:  0.7901, Adjusted R-squared:  0.7578
## F-statistic: 24.47 on 2 and 13 DF,  p-value: 3.915e-05
```

```
lm_robustness_null_3 <- lm(Robustness_Null ~ poly(Time, 2) + NMDS1,
                            data = robustness_data)
summary(lm_robustness_null_3)
```

```
##
## Call:
## lm(formula = Robustness_Null ~ poly(Time, 2) + NMDS1, data = robustness_data)
##
## Residuals:
##       Min        1Q    Median        3Q       Max
## -0.051478 -0.010425  0.005161  0.011965  0.040368
##
## Coefficients:
##                 Estimate Std. Error t value Pr(>|t|)
## (Intercept)     0.618186   0.006408  96.469  < 2e-16 ***
## poly(Time, 2)1  0.001709   0.030549   0.056 0.956320
## poly(Time, 2)2 -0.060225   0.029049  -2.073 0.060345 .
## NMDS1          -0.051023   0.009678  -5.272 0.000197 ***
```

```
## ---
## Signif. codes:  0 '***' 0.001 '**' 0.01 '*' 0.05 '.' 0.1 ' ' 1
##
## Residual standard error: 0.02563 on 12 degrees of freedom
## Multiple R-squared:  0.8455, Adjusted R-squared:  0.8068
## F-statistic: 21.89 on 3 and 12 DF,  p-value: 3.719e-05
```

```
#model selection
aic_lm_robustness_null <- AIC(lm_robustness_null_1, lm_robustness_null_2,
                              lm_robustness_null_3)
aic_lm_robustness_null
```

```
##                      df       AIC
## lm_robustness_null_1  6 -66.26977
## lm_robustness_null_2  4 -63.54319
## lm_robustness_null_3  5 -66.44154
```

```
#diagnostic
shapiro.test(lm_robustness_null_3$residuals)
```

```
##
##  Shapiro-Wilk normality test
##
## data:  lm_robustness_null_3$residuals
## W = 0.94526, p-value = 0.4185
```

```
plot(lm_robustness_null_3)
```

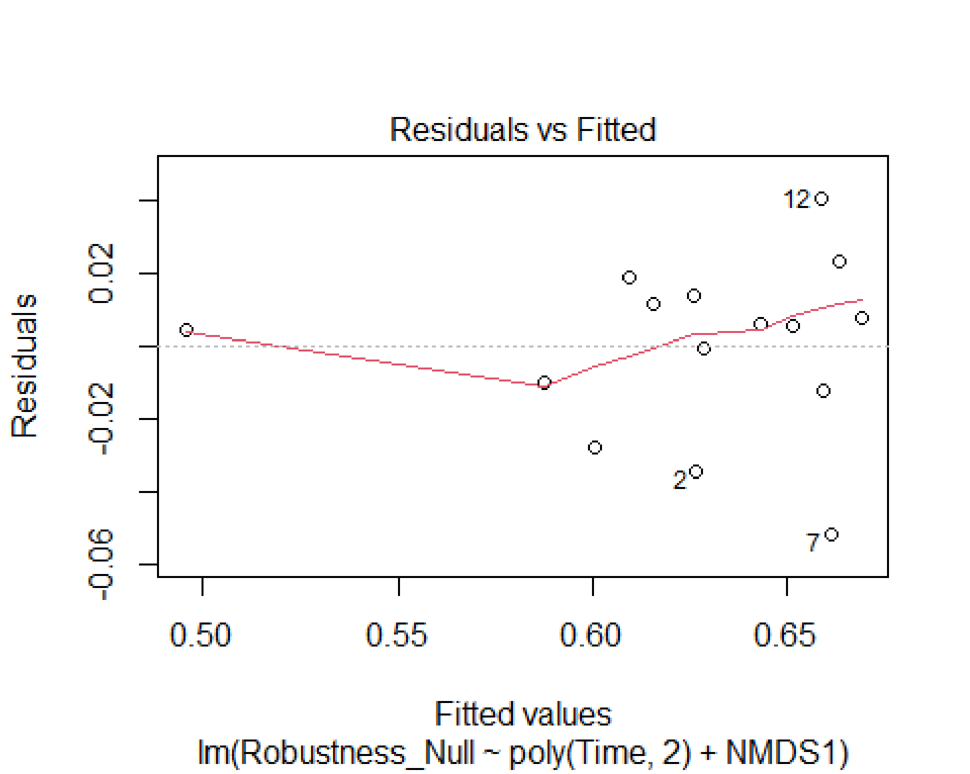

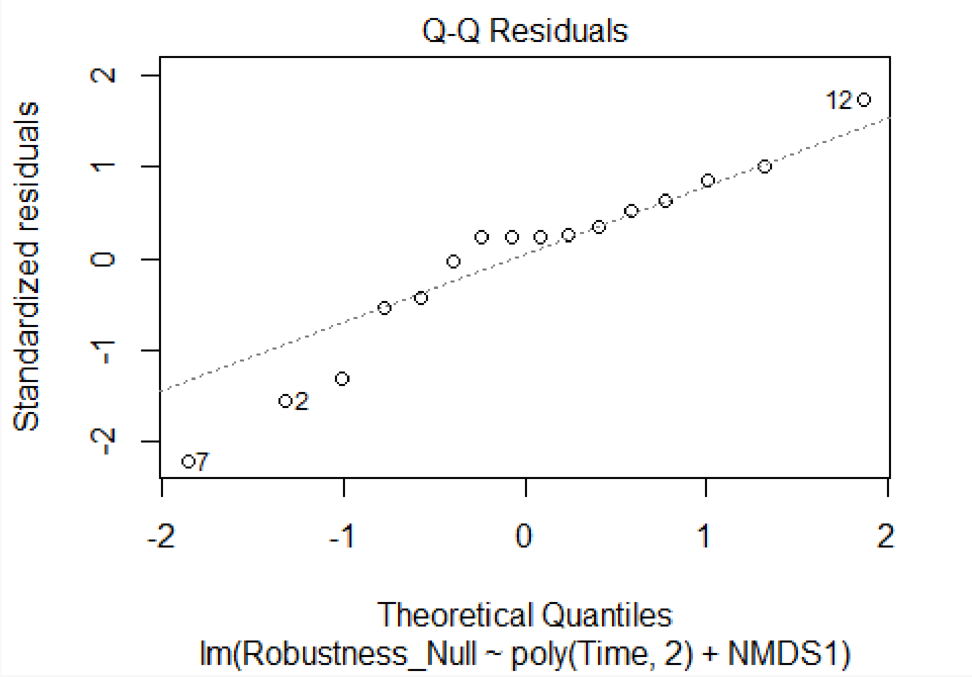
Q-Q Residuals
Standardized residuals
Theoretical Quantiles
lm(Robustness_Null ~ poly(Time, 2) + NMDS1)
12
2
7

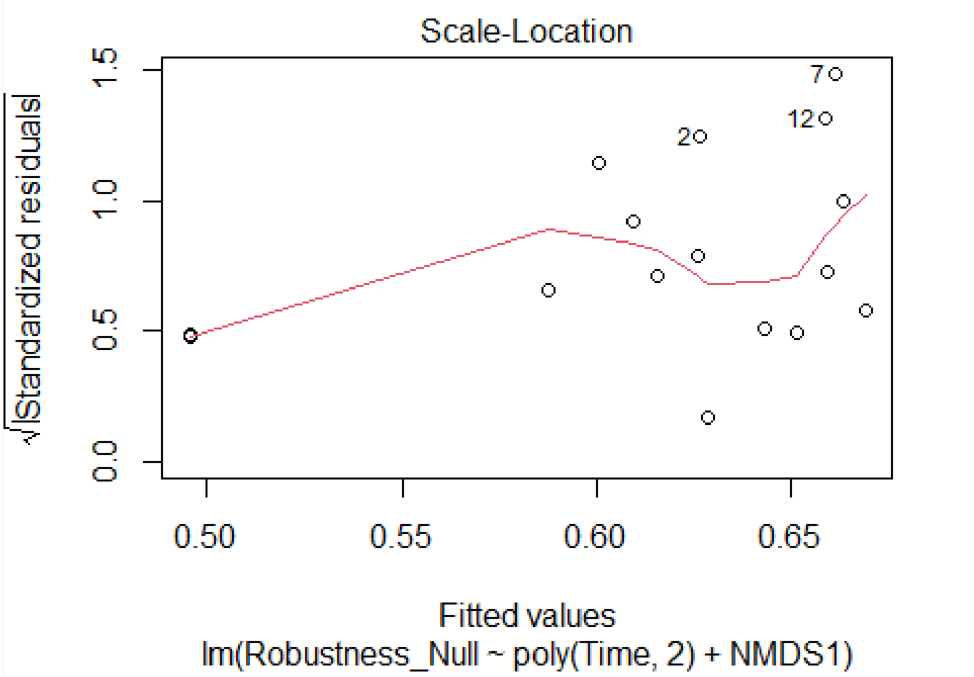
Scale-Location
√|Standardized residuals|
Fitted values
lm(Robustness_Null ~ poly(Time, 2) + NMDS1)
7
12
2

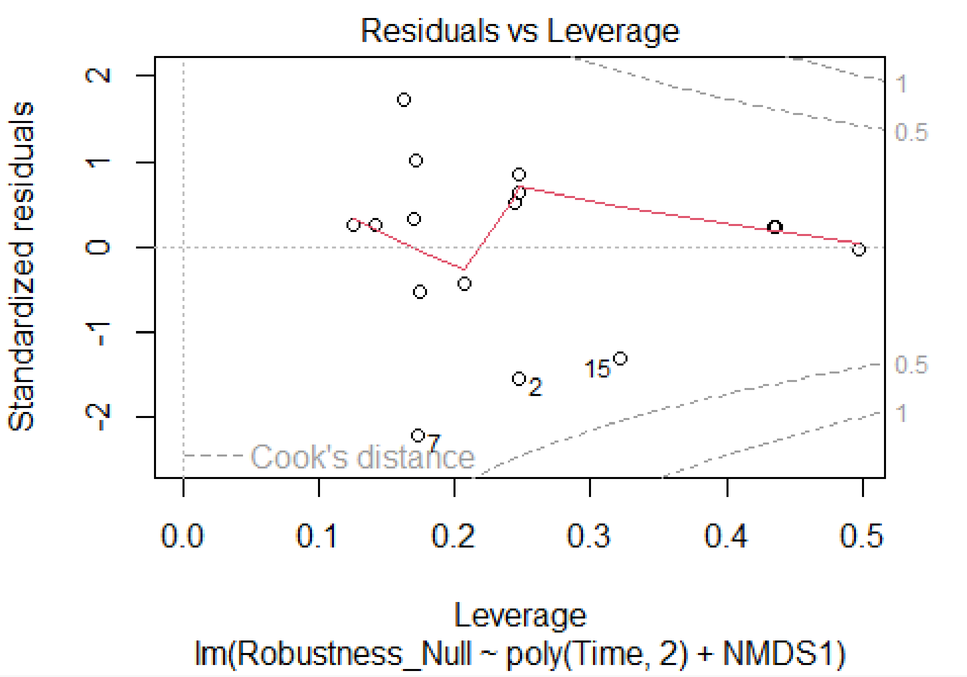

```r
##beta regression models##
betareg_robustness_null_1 <- betareg(Robustness_Null ~ as.factor(Time) + NMDS1,
                                      data = robustness_data)
summary(betareg_robustness_null_1)

##
## Call:
## betareg(formula = Robustness_Null ~ as.factor(Time) + NMDS1, data =
robustness_data)
##
## Standardized weighted residuals 2:
##     Min      1Q  Median      3Q     Max
## -2.3465 -0.5359  0.4287  0.6864  1.8052
##
## Coefficients (mean model with logit link):
##                     Estimate Std. Error z value Pr(>|z|)
## (Intercept)          0.43381    0.04686   9.258  < 2e-16 ***
## as.factor(Time)66    0.10344    0.06559   1.577   0.1148
## as.factor(Time)111   0.14396    0.06543   2.200   0.0278 *
## as.factor(Time)141  -0.03702    0.07760  -0.477   0.6333
## NMDS1               -0.20393    0.03429  -5.948 2.72e-09 ***
##
## Phi coefficients (precision model with identity link):
##       Estimate Std. Error z value Pr(>|z|)
## (phi)    509.1      179.8   2.831  0.00464 **
## ---
## Signif. codes:  0 '***' 0.001 '**' 0.01 '*' 0.05 '.' 0.1 ' ' 1
```

```
##
## Type of estimator: ML (maximum likelihood)
## Log-likelihood: 38.83 on 6 Df
## Pseudo R-squared: 0.8487
## Number of iterations: 99 (BFGS) + 3 (Fisher scoring)
```

```
betareg_robustness_null_2 <- betareg(Robustness_Null ~ Time + NMDS1,
                                      data = robustness_data)
summary(betareg_robustness_null_2)
```

```
##
## Call:
## betareg(formula = Robustness_Null ~ Time + NMDS1, data = robustness_data)
##
## Standardized weighted residuals 2:
##     Min      1Q  Median      3Q     Max
## -2.0376 -0.2832  0.1999  0.4037  1.8787
##
## Coefficients (mean model with logit link):
##              Estimate Std. Error z value Pr(>|z|)
## (Intercept)  0.4499453  0.0642657   7.001 2.54e-12 ***
## Time         0.0004310  0.0006922   0.623    0.533
## NMDS1       -0.2506519  0.0367928  -6.813 9.59e-12 ***
##
## Phi coefficients (precision model with identity link):
##       Estimate Std. Error z value Pr(>|z|)
## (phi)    339.6      119.9   2.832  0.00462 **
## ---
## Signif. codes:  0 '***' 0.001 '**' 0.01 '*' 0.05 '.' 0.1 ' ' 1
##
## Type of estimator: ML (maximum likelihood)
## Log-likelihood: 35.59 on 4 Df
## Pseudo R-squared: 0.7728
## Number of iterations: 122 (BFGS) + 3 (Fisher scoring)
```

```
betareg_robustness_null_3 <- betareg(Robustness_Null ~ poly(Time, 2) + NMDS1,
                                      data = robustness_data)
summary(betareg_robustness_null_3)
```

```
##
## Call:
## betareg(formula = Robustness_Null ~ poly(Time, 2) + NMDS1, data =
robustness_data)
##
## Standardized weighted residuals 2:
##     Min      1Q  Median      3Q     Max
## -2.4925 -0.6099  0.2780  0.5908  2.0590
##
## Coefficients (mean model with logit link):
##                 Estimate Std. Error z value Pr(>|z|)
## (Intercept)     0.486488   0.024088  20.196  < 2e-16 ***
```

```
## poly(Time, 2)1  0.007327   0.114517   0.064   0.9490
## poly(Time, 2)2 -0.263517   0.109462  -2.407   0.0161 *
## NMDS1          -0.209430   0.035769  -5.855 4.77e-09 ***
##
## Phi coefficients (precision model with identity link):
##       Estimate Std. Error z value Pr(>|z|)
## (phi)    461.2      162.9   2.831  0.00464 **
## ---
## Signif. codes:  0 '***' 0.001 '**' 0.01 '*' 0.05 '.' 0.1 ' ' 1
##
## Type of estimator: ML (maximum likelihood)
## Log-likelihood: 38.04 on 5 Df
## Pseudo R-squared: 0.8323
## Number of iterations: 304 (BFGS) + 3 (Fisher scoring)
```

```
#model selection
aic_betareg_robustness_null <- AIC(betareg_robustness_null_1,
                                   betareg_robustness_null_2,
                                   betareg_robustness_null_3)
aic_betareg_robustness_null
```

```
##                           df       AIC
## betareg_robustness_null_1  6 -65.65743
## betareg_robustness_null_2  4 -63.17915
## betareg_robustness_null_3  5 -66.07731
```

```
#diagnostic
shapiro.test(betareg_robustness_null_3$residuals)
```

```
##
##  Shapiro-Wilk normality test
##
## data:  betareg_robustness_null_3$residuals
## W = 0.94342, p-value = 0.3931
```

```
plot(betareg_robustness_null_3)
```

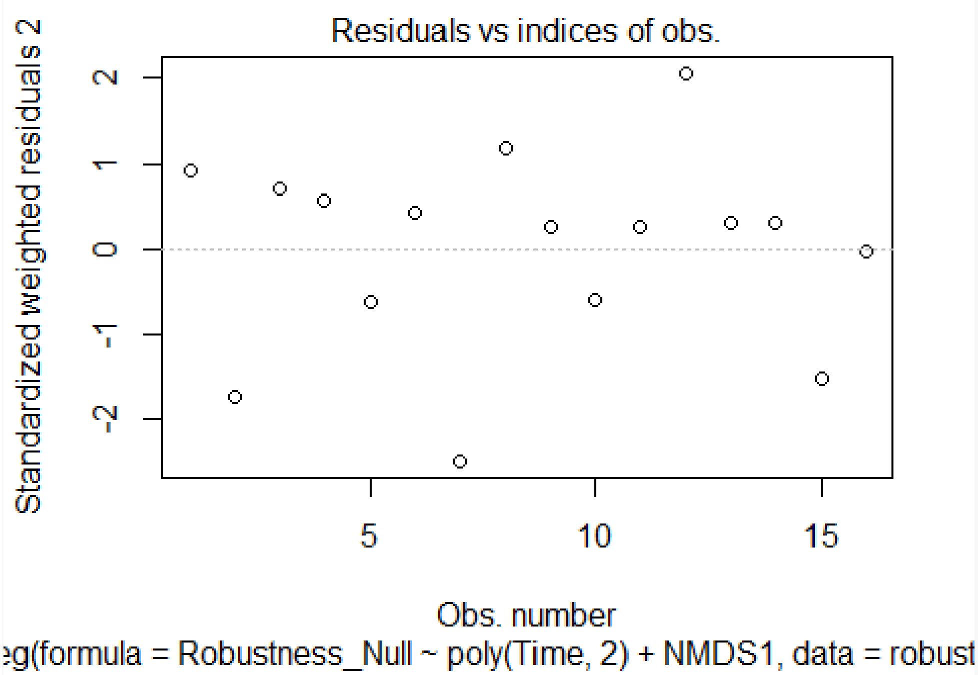
Residuals vs indices of obs.
Standardized weighted residuals 2
Obs. number
eg(formula = Robustness_Null ~ poly(Time, 2) + NMDS1, data = robust

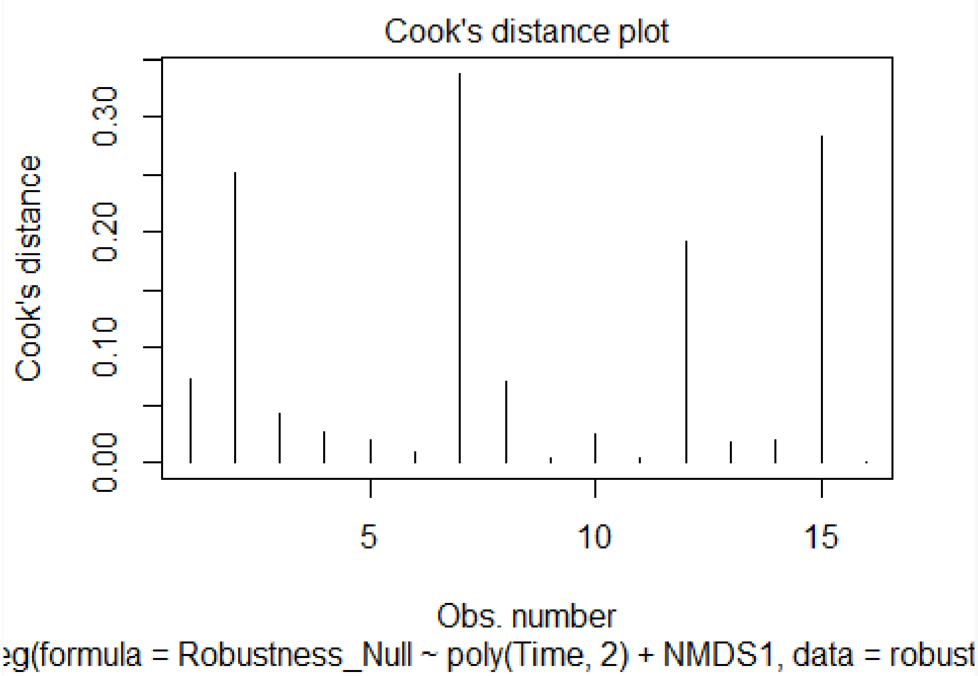
Cook's distance plot
Cook's distance
Obs. number
eg(formula = Robustness_Null ~ poly(Time, 2) + NMDS1, data = robust

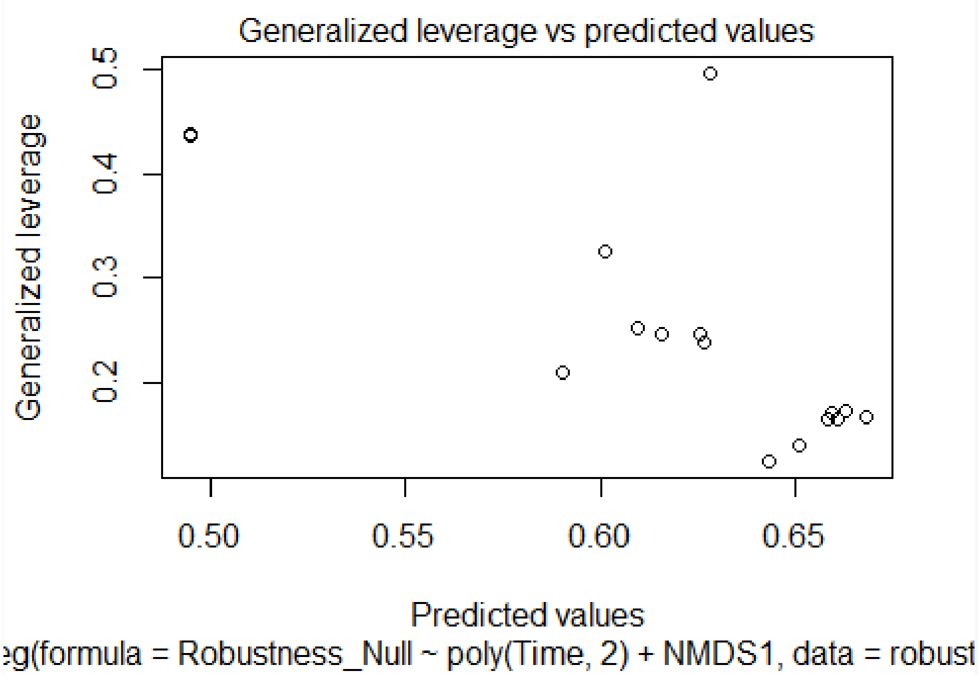

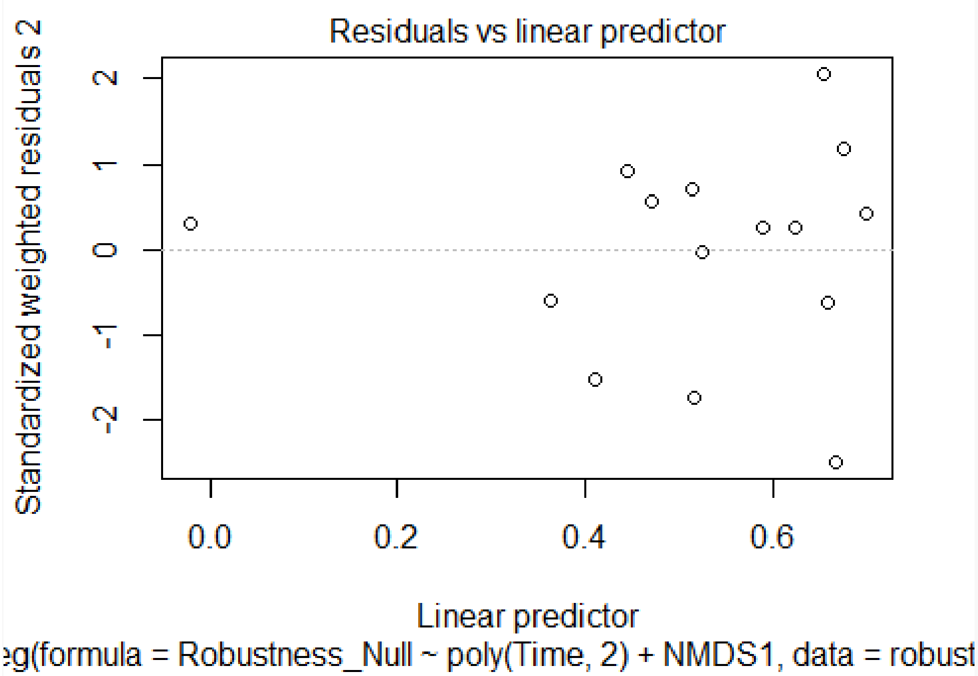

```
#Type-II ANOVA
anovaII_robustness_null <- Anova(betareg_robustness_null_3, type = "II")
anovaII_robustness_null
```

```
## Analysis of Deviance Table (Type II tests)
##
## Response: Robustness_Null
##                Df   Chisq Pr(>Chisq)
## poly(Time, 2)  2  6.2928    0.04301 *
## NMDS1          1 34.2823  4.767e-09 ***
## ---
## Signif. codes:  0 '***' 0.001 '**' 0.01 '*' 0.05 '.' 0.1 ' ' 1
```

```
###Z-SCORES~TIME+NMDS1###
##linear models##
lm_zscores_1 <- lm(Z_Scores ~ as.factor(Time) + NMDS1, data = robustness_data)
summary(lm_zscores_1)
```

```
##
## Call:
## lm(formula = Z_Scores ~ as.factor(Time) + NMDS1, data = robustness_data)
##
## Residuals:
##      Min       1Q   Median       3Q      Max
## -26.9824  -7.9248  -0.1945  10.2155  26.9824
##
## Coefficients:
##                     Estimate Std. Error t value Pr(>|t|)
## (Intercept)          12.8669    10.2080   1.260    0.239
## as.factor(Time)66    18.5076    13.2033   1.402    0.195
## as.factor(Time)111    0.4908    13.1555   0.037    0.971
## as.factor(Time)141    3.7336    15.7740   0.237    0.818
## NMDS1               -10.2380    15.0847  -0.679    0.514
##
## Residual standard error: 18.16 on 9 degrees of freedom
##   (2 osservazioni eliminate a causa di valori mancanti)
## Multiple R-squared:  0.3191, Adjusted R-squared:  0.01653
## F-statistic: 1.055 on 4 and 9 DF,  p-value: 0.432
```

```
lm_zscores_2 <- lm(Z_Scores ~ Time + NMDS1, data = robustness_data)
summary(lm_zscores_2)
```

```
##
## Call:
## lm(formula = Z_Scores ~ Time + NMDS1, data = robustness_data)
##
## Residuals:
##      Min       1Q   Median       3Q      Max
## -27.5337 -11.8999   0.2303  12.0688  22.9301
##
## Coefficients:
##               Estimate Std. Error t value Pr(>|t|)
## (Intercept)  15.583372  11.444425   1.362    0.201
## Time          0.008745   0.113331   0.077    0.940
## NMDS1       -18.808043  14.153651  -1.329    0.211
```

```
##
## Residual standard error: 18.45 on 11 degrees of freedom
##   (2 osservazioni eliminate a causa di valori mancanti)
## Multiple R-squared:  0.1411, Adjusted R-squared:  -0.0151
## F-statistic: 0.9033 on 2 and 11 DF,  p-value: 0.4333
```

```
lm_zscores_3 <- lm(Z_Scores ~ poly(Time, 2) + NMDS1, data = robustness_data)
summary(lm_zscores_3)
```

```
##
## Call:
## lm(formula = Z_Scores ~ poly(Time, 2) + NMDS1, data = robustness_data)
##
## Residuals:
##     Min      1Q  Median      3Q     Max
## -29.529 -10.040  -1.898  11.970  31.474
##
## Coefficients:
##                 Estimate Std. Error t value Pr(>|t|)
## (Intercept)       16.134      6.439   2.506   0.0311 *
## poly(Time, 2)1    -4.208     21.904  -0.192   0.8515
## poly(Time, 2)2   -21.691     20.841  -1.041   0.3225
## NMDS1            -15.919     14.371  -1.108   0.2939
## ---
## Signif. codes:  0 '***' 0.001 '**' 0.01 '*' 0.05 '.' 0.1 ' ' 1
##
## Residual standard error: 18.38 on 10 degrees of freedom
##   (2 osservazioni eliminate a causa di valori mancanti)
## Multiple R-squared:  0.225,  Adjusted R-squared:  -0.007481
## F-statistic: 0.9678 on 3 and 10 DF,  p-value: 0.4456
```

Finally, we plot test model robustness in relation to time since deglaciation and NMDS1 coordinates of motifs' ordination.

```
###GRAPHS###
#robustness~time
figure_4a <- ggplot(data = robustness_data,
                    mapping = aes(Time, Robustness, color = Stage)) +
                    geom_point() +
                    geom_smooth(method = glm, formula = y ~ poly(x, 2),
                                color = "black", linewidth = 0.8) +
                    scale_color_manual(values = c("blue3", "green3",
                                                  "yellow3", "red3")) +
                    theme_bw() +
                    theme(panel.grid.major = element_blank(),
                          panel.grid.minor = element_blank(),
                          axis.title = element_text(size = 10)) +
                    labs(x = "Time [years since deglaciation]",
                         y = "Robustness [R]",
                         title = "A")
figure_4a
```

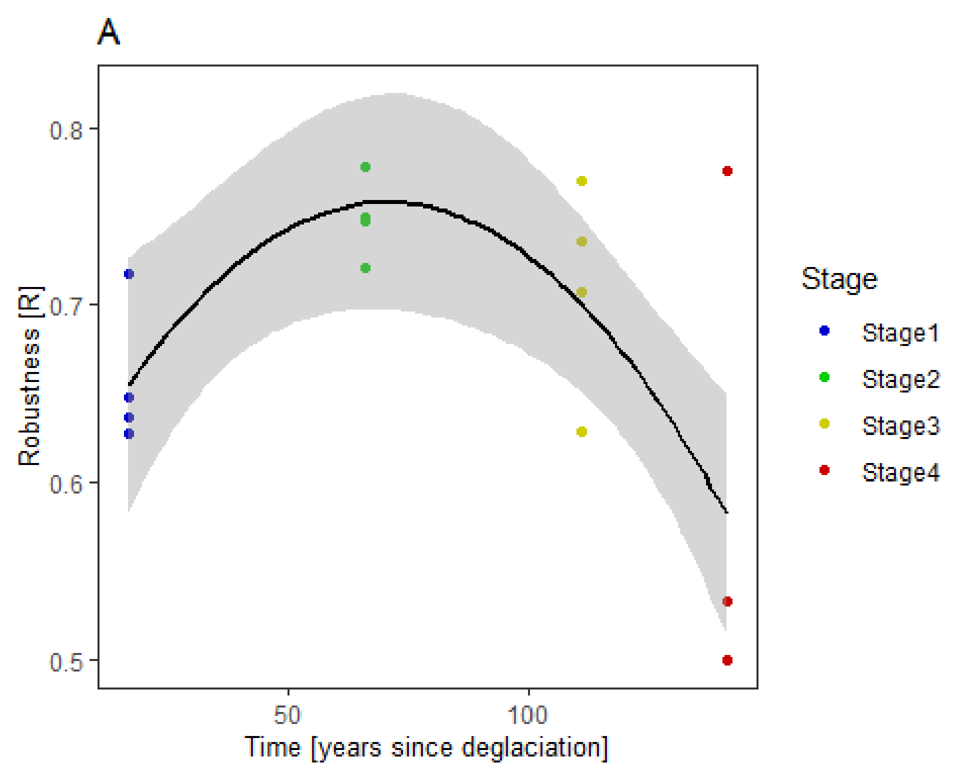

```
#robustness~NMDS1
figure_4b <- ggplot(data = robustness_data,
                    mapping = aes(NMDS1, Robustness, color = Stage)) +
                    geom_point() +
                    geom_smooth(method = glm, formula = y ~ x, color="black",
                                linewidth = 0.8) +
                    scale_color_manual(values = c("blue3", "green3",
                                                  "yellow3", "red3")) +
                    theme_bw() +
                    theme(panel.grid.major = element_blank(),
                          panel.grid.minor = element_blank(),
                          axis.title = element_text(size = 10)) +
                    labs(x = "NMDS1 [x coords of motifs NMDS]",
                         y = "Robustness [R]",
                         title = "B")
figure_4b
```

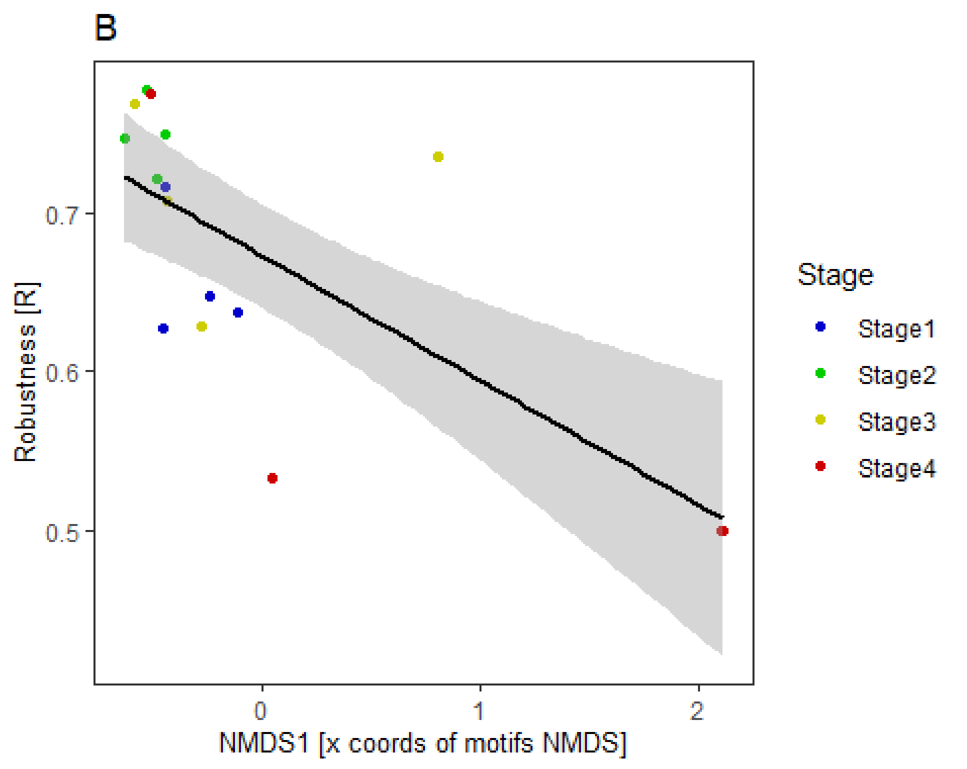

#### (c) Plant species' roles

In this section, we unfold all analyses concerning plant species roles within network motifs.

We begin by calculating plant species' contribution to node positions within all possible motifs up to 5 nodes in observed networks (at 'stage' level) using 'node_positions' function of 'bmotif' package (Mora et al. 2018, Simmons et al. 2019b). For a given species in a given position in a motif, contribution was defined as the sum of focal species' interactions divided by the sum of all interactions in that motif (Simmons et al. 2019b).

```
###CALCULATE NODE POSITIONS (SAMPLED NETWORKS)###
stage_names <- c("S1", "S2", "S3", "S4")
for (stage in stage_names) {
  assign(paste0("np_", stage), node_positions(
    bipnet_stages_list[[stage]],
    six_node = FALSE,
    level = "rows",
    weights_method = "all",
    weights_combine = "mean",
    normalisation = "none"
  ))
}
```

We then identify plant species most representative node positions for each stage (those with highest contribution).

```
###MOST REPRESENTATIVE NODE POSITIONS FOR SELECTED SPECIES###
#create separate dataframes for "contribution"
df_S1 <- np_S1[["contribution"]]
df_S2 <- np_S2[["contribution"]]
df_S3 <- np_S3[["contribution"]]
df_S4 <- np_S4[["contribution"]]
#upload contribution dataframes to a list
df_list <- list(df_S1 = df_S1, df_S2 = df_S2, df_S3 = df_S3, df_S4 = df_S4)
#select species of interest
species_of_interest <- "Linaria alpina"
#find top 5 most representative node positions for selected species in each stage
for (i in seq_along(df_list)) {
  df <- df_list[[i]]
  #skip stages where the species is not present
  row_of_interest <- df[species_of_interest, , drop = FALSE]
  if(all(is.na(row_of_interest))) {
    cat(species_of_interest, "is not present in Stage", i, "\n")
    next
  }
  row_of_interest <- row_of_interest[, !is.na(row_of_interest)]
  row_values <- unlist(row_of_interest)
  top_indices <- order(row_values, decreasing = TRUE)[1:5]
  top_values <- row_values[top_indices]
  top_column_names <- names(top_values)
  cat("top 5 values for", species_of_interest, "in Stage", i, "are:",
      paste(top_values, collapse = ", "), "\n")
  cat("top 5 node positions for:", species_of_interest, "in Stage", i,
      "are:", paste(top_column_names, collapse = ", "), "\n\n")
}

## top 5 values for Linaria alpina in Stage 1 are: 1, 0.282303798712467,
0.124099891400876, 0.11381709414287, 0.0865448566126484
## top 5 node positions for: Linaria alpina in Stage 1 are: np2, np6, np8, np11,
np24
##
## Linaria alpina is not present in Stage 2
## Linaria alpina is not present in Stage 3
## Linaria alpina is not present in Stage 4
```

We build null model networks by randomising observed networks 100 times following 4 different null model approaches (the same ones described in the 'motifs' prevalence' section). For each null model approach, we calculate Z-scores and identify the most over- and under-represented node position within each stage and across all stages.

```
###REFERENCE NULL MODEL ("r00_ind")###
##initialise vectors to store entries above treshold##
#total (|Z|>1.96)
num_entries_above_threshold_vec <- numeric(length(stage_names))
```

```r
total_entries_adjusted_vec <- numeric(length(stage_names))
#over (Z>1.96)
num_entries_above_threshold_over_vec <- numeric(length(stage_names))
total_entries_adjusted_over_vec <- numeric(length(stage_names))
#under (Z<-1.96)
num_entries_above_threshold_under_vec <- numeric(length(stage_names))
total_entries_adjusted_under_vec <- numeric(length(stage_names))
##initialise list to store Z-scores for each stage##
z_scores_np_list <- list()
##randomise networks, calculate node positions and calculate Z-scores##
#loop across stages
for (stage in stage_names) {
  #randomise
  bipnet_np_null <- vegan::nullmodel(bipnet_stages_list[[stage]], "r00_ind")
  bipnet_np_null_2 <- simulate(bipnet_np_null, nsim = 100, seed = 1)
  bipnet_np_null_2_list <- lapply(seq_len(100),
                                  function(index) bipnet_np_null_2[, , index])
  bipnet_np_null_2_list <- lapply(bipnet_np_null_2_list, function(mat) {
    rownames(mat) <- NULL
    colnames(mat) <- NULL
    return(mat)
  })
  #initialise a list to store results
  matrix_list_np <- list()
  #calculate contribution to node positions
  for (i in 1:100) {
    result_matrix_np <- node_positions(bipnet_np_null_2_list[[i]],
                                       six_node = FALSE,
                                       level = "rows",
                                       weights_method = "contribution",
                                       weights_combine = "mean",
                                       normalisation = "none")
    result_matrix_np[is.na(result_matrix_np)] <- 0
    matrix_list_np[[i]] <- result_matrix_np
  }
  #get the right np list
  np_list <- get(paste0("np_", stage), envir = .GlobalEnv)
  #create mean and sd matrices
  mean_matrix_np <- apply(array(unlist(matrix_list_np),
                                c(dim(matrix_list_np[[1]]),
                                  length(matrix_list_np))), c(1, 2), mean)
  sd_matrix_np <- apply(array(unlist(matrix_list_np),
                              c(dim(matrix_list_np[[1]]),
                                length(matrix_list_np))), c(1, 2), sd)
  #calculate mean and sd
  results_mean_np <- as.data.frame(mean_matrix_np, row.names =
                                     rownames(np_list[["contribution"]]),
                                   col.names =
                                     colnames(np_list[["contribution"]]))
  results_sd_np <- as.data.frame(sd_matrix_np, row.names =
                                   rownames(np_list[["contribution"]]),
```

```r
                                  col.names = colnames(np_list[["contribution"]]))
  #calculate Z-scores and upload to the list
  np_0 <- np_list[["contribution"]]
  np_0[is.na(np_0)] <- 0
  z_scores_np <- ((np_0 - results_mean_np) / (results_sd_np))
  z_scores_np <- as.data.frame(z_scores_np, row.names = rownames(np_0),
                               col.names = colnames(np_0))
  z_scores_np[z_scores_np == -Inf] <- NaN
  z_scores_np[z_scores_np == Inf] <- NaN
  z_scores_np_list[[paste0("z_scores_np_", stage)]] <- z_scores_np
  ##percentage of entries with |Z|>1.96##
  num_entries_above_threshold_np <- sum(abs(z_scores_np) > 1.96 &
                                        !is.na(z_scores_np))
  total_entries_adjusted_np <- sum(!is.na(z_scores_np))
  num_entries_above_threshold_vec[stage] <- sum(abs(z_scores_np) > 1.96 &
                                                !is.na(z_scores_np))
  total_entries_adjusted_vec[stage] <- sum(!is.na(z_scores_np))
  cat("number of entries above threshold in", stage, "is:",
      num_entries_above_threshold_np, "\n")
  percentage_above_threshold_np <- (num_entries_above_threshold_np /
                                    total_entries_adjusted_np) * 100
  cat("percentage of entries above threshold in", stage, "is:",
      percentage_above_threshold_np, "%\n\n")
  ##percentage of entries with Z>1.96##
  num_entries_above_threshold_over_np <- sum(z_scores_np > 1.96 &
                                             !is.na(z_scores_np))
  total_entries_adjusted_over_np <- sum(!is.na(z_scores_np))
  num_entries_above_threshold_over_vec[stage] <- sum(z_scores_np > 1.96 &
                                                     !is.na(z_scores_np))
  total_entries_adjusted_over_vec[stage] <- sum(!is.na(z_scores_np))
  cat("number of over-represented entries in", stage, "is:",
      num_entries_above_threshold_over_np, "\n")
  percentage_above_threshold_over_np <- (num_entries_above_threshold_over_np /
total_entries_adjusted_over_np) * 100
  cat("percentage of over-represented entries in", stage, "is:",
      percentage_above_threshold_over_np, "%\n\n")
  ##percentage of entries with Z<-1.96##
  num_entries_above_threshold_under_np <- sum(z_scores_np < -1.96 &
                                              !is.na(z_scores_np))
  total_entries_adjusted_under_np <- sum(!is.na(z_scores_np))
  num_entries_above_threshold_under_vec[stage] <- sum(z_scores_np < -1.96 &
                                                      !is.na(z_scores_np))
  total_entries_adjusted_under_vec[stage] <- sum(!is.na(z_scores_np))
  cat("number of under-represented entries in", stage, "is:",
      num_entries_above_threshold_under_np, "\n")
  percentage_above_threshold_under_np <- (num_entries_above_threshold_under_np /
total_entries_adjusted_under_np) * 100
  cat("percentage of under-represented entries in", stage, "is:",
      percentage_above_threshold_under_np, "%\n\n")
  ##most over-represented and under-represented node positions##
  column_means_np <- colMeans(z_scores_np)
```

```
  max_mean_column_np <- names(column_means_np)[which.max(column_means_np)]
  min_mean_column_np <- names(column_means_np)[which.min(column_means_np)]
  cat("the most over-represented node position in", stage, "is",
      max_mean_column_np, "\n")
  cat("the most under-represented node position in", stage, "is",
      min_mean_column_np, "\n")
}
```

```
## number of entries above threshold in S1 is: 437

## percentage of entries above threshold in S1 is: 84.68992 %

## number of over-represented entries in S1 is: 97

## percentage of over-represented entries in S1 is: 18.79845 %

## number of under-represented entries in S1 is: 340

## percentage of under-represented entries in S1 is: 65.89147 %

## the most over-represented node position in S1 is np18

## the most under-represented node position in S1 is np35

## number of entries above threshold in S2 is: 819

## percentage of entries above threshold in S2 is: 86.11987 %

## number of over-represented entries in S2 is: 184

## percentage of over-represented entries in S2 is: 19.34805 %

## number of under-represented entries in S2 is: 635

## percentage of under-represented entries in S2 is: 66.77182 %

## the most over-represented node position in S2 is np6

## the most under-represented node position in S2 is np35

## number of entries above threshold in S3 is: 637

## percentage of entries above threshold in S3 is: 88.22715 %

## number of over-represented entries in S3 is: 150

## percentage of over-represented entries in S3 is: 20.77562 %

## number of under-represented entries in S3 is: 487

## percentage of under-represented entries in S3 is: 67.45152 %

## the most over-represented node position in S3 is np6

## the most under-represented node position in S3 is np35
```

```
## number of entries above threshold in S4 is: 266

## percentage of entries above threshold in S4 is: 77.55102 %

## number of over-represented entries in S4 is: 30

## percentage of over-represented entries in S4 is: 8.746356 %

## number of under-represented entries in S4 is: 236

## percentage of under-represented entries in S4 is: 68.80466 %

## the most over-represented node position in S4 is np18

## the most under-represented node position in S4 is np35

##percentage of entries with |Z|>1.96 across all stages##
num_entries_above_threshold_np_tot <- sum(num_entries_above_threshold_vec)
total_entries_adjusted_np_tot <- sum(total_entries_adjusted_vec)
percentage_above_threshold_np_tot <- (num_entries_above_threshold_np_tot /
                                        total_entries_adjusted_np_tot) * 100
cat("number of entries above threshold across all stages: ",
    num_entries_above_threshold_np_tot, "\n")

## number of entries above threshold across all stages:  2159

cat("percentage of entries above threshold across all stages: ",
    percentage_above_threshold_np_tot, "%\n")

## percentage of entries above threshold across all stages:  85.26856 %

##percentage of entries with Z>1.96 across all stages##
num_entries_above_threshold_np_over_tot <-
  sum(num_entries_above_threshold_over_vec)
total_entries_adjusted_np_over_tot <- sum(total_entries_adjusted_over_vec)
percentage_above_threshold_np_over_tot <-
  (num_entries_above_threshold_np_over_tot /
     total_entries_adjusted_np_over_tot) * 100
cat("number of over-represented entries across all stages: ",
    num_entries_above_threshold_np_over_tot, "\n")

## number of over-represented entries across all stages:  461

cat("percentage of over-represented entries across all stages: ",
    percentage_above_threshold_np_over_tot, "%\n")

## percentage of over-represented entries across all stages:  18.20695 %

##percentage of entries with Z<-1.96 across all stages##
num_entries_above_threshold_np_under_tot <-
  sum(num_entries_above_threshold_under_vec)
total_entries_adjusted_np_under_tot <- sum(total_entries_adjusted_under_vec)
```

```r
percentage_above_threshold_np_under_tot <-
  (num_entries_above_threshold_np_under_tot /
     total_entries_adjusted_np_under_tot) * 100
cat("number of under-represented entries across all stages: ",
    num_entries_above_threshold_np_under_tot, "\n")

## number of under-represented entries across all stages:  1698

cat("percentage of under-represented entries across all stages: ",
    percentage_above_threshold_np_under_tot, "%\n")

## percentage of under-represented entries across all stages:  67.06161 %

###ADDITIONAL NULL MODELS ("r0_ind", "c0_ind" and "vaznull")###
###"r0_ind"###
##initialise vectors to store entries above treshold##
#total (|Z|>1.96)
num_entries_above_threshold_vec_r <- numeric(length(stage_names))
total_entries_adjusted_vec_r <- numeric(length(stage_names))
#over (Z>1.96)
num_entries_above_threshold_over_vec_r <- numeric(length(stage_names))
total_entries_adjusted_over_vec_r <- numeric(length(stage_names))
#under (Z<-1.96)
num_entries_above_threshold_under_vec_r <- numeric(length(stage_names))
total_entries_adjusted_under_vec_r <- numeric(length(stage_names))
##initialise list to store Z-scores for each stage##
z_scores_np_list_r <- list()
##randomise networks, calculate node positions and calculate Z-scores##
#loop across stages
for (stage in stage_names) {
  #randomise
  bipnet_np_null_r <- vegan::nullmodel(bipnet_stages_list[[stage]], "r0_ind")
  bipnet_np_null_2_r <- simulate(bipnet_np_null_r, nsim = 100, seed = 1)
  bipnet_np_null_2_list_r <- lapply(seq_len(100),
                                    function(index)
                                      bipnet_np_null_2_r[, , index])
  bipnet_np_null_2_list_r <- lapply(bipnet_np_null_2_list_r, function(mat) {
    rownames(mat) <- NULL
    colnames(mat) <- NULL
    return(mat)
  })
  #initialise a list to store results
  matrix_list_np_r <- list()
  #calculate contribution to node positions
  for (i in 1:100) {
    result_matrix_np_r <- node_positions(bipnet_np_null_2_list_r[[i]],
                                         six_node = FALSE,
                                         level = "rows",
                                         weights_method = "contribution",
```

```
                                            weights_combine = "mean",
                                            normalisation = "none")
  result_matrix_np_r[is.na(result_matrix_np_r)] <- 0
  matrix_list_np_r[[i]] <- result_matrix_np_r
}
#get the right np list
np_list_r <- get(paste0("np_", stage), envir = .GlobalEnv)
#create mean and sd matrices
mean_matrix_np_r <- apply(array(unlist(matrix_list_np_r),
                                c(dim(matrix_list_np_r[[1]]),
                                  length(matrix_list_np_r))), c(1, 2), mean)
sd_matrix_np_r <- apply(array(unlist(matrix_list_np_r),
                              c(dim(matrix_list_np_r[[1]]),
                                length(matrix_list_np_r))), c(1, 2), sd)
#calculate mean and sd
results_mean_np_r <- as.data.frame(mean_matrix_np_r, row.names =
                                     rownames(np_list_r[["contribution"]]),
                                   col.names =
                                     colnames(np_list_r[["contribution"]]))
results_sd_np_r <- as.data.frame(sd_matrix_np_r, row.names =
                                   rownames(np_list_r[["contribution"]]),
                                 col.names =
                                   colnames(np_list_r[["contribution"]]))
#calculate Z-scores and upload the list
np_0_r <- np_list_r[["contribution"]]
np_0_r[is.na(np_0_r)] <- 0
z_scores_np_r <- ((np_0_r - results_mean_np_r) / (results_sd_np_r))
z_scores_np_r <- as.data.frame(z_scores_np_r, row.names = rownames(np_0_r),
                               col.names = colnames(np_0_r))
z_scores_np_r[z_scores_np_r == -Inf] <- NaN
z_scores_np_r[z_scores_np_r == Inf] <- NaN
z_scores_np_list_r[[paste0("z_scores_np_r_", stage)]] <- z_scores_np_r
##percentage of entries with |Z|>1.96##
num_entries_above_threshold_np_r <- sum(abs(z_scores_np_r) > 1.96 &
                                          !is.na(z_scores_np_r))
total_entries_adjusted_np_r <- sum(!is.na(z_scores_np_r))
num_entries_above_threshold_vec_r[stage] <- sum(abs(z_scores_np_r) > 1.96 &
                                                  !is.na(z_scores_np_r))
total_entries_adjusted_vec_r[stage] <- sum(!is.na(z_scores_np_r))
cat("number of entries above threshold in", stage, "is:",
    num_entries_above_threshold_np_r, "\n")
percentage_above_threshold_np_r <- (num_entries_above_threshold_np_r /
                                      total_entries_adjusted_np_r) * 100
cat("percentage of entries above threshold in", stage, "is:",
    percentage_above_threshold_np_r, "%\n\n")
##percentage of entries with Z>1.96##
num_entries_above_threshold_over_np_r <- sum(z_scores_np_r > 1.96 &
                                               !is.na(z_scores_np_r))
total_entries_adjusted_over_np_r <- sum(!is.na(z_scores_np_r))
num_entries_above_threshold_over_vec_r[stage] <- sum(z_scores_np_r > 1.96 &
                                                       !is.na(z_scores_np_r))
```

```
  total_entries_adjusted_over_vec_r[stage] <- sum(!is.na(z_scores_np_r))
  cat("number of over-represented entries in", stage, "is:",
      num_entries_above_threshold_over_np_r, "\n")
  percentage_above_threshold_over_np_r <-
    (num_entries_above_threshold_over_np_r /
       total_entries_adjusted_over_np_r) * 100
  cat("percentage of over-represented entries in", stage, "is:",
      percentage_above_threshold_over_np_r, "%\n\n")
  ##percentage of entries with Z<-1.96##
  num_entries_above_threshold_under_np_r <- sum(z_scores_np_r < -1.96 &
                                                 !is.na(z_scores_np_r))
  total_entries_adjusted_under_np_r <- sum(!is.na(z_scores_np_r))
  num_entries_above_threshold_under_vec_r[stage] <- sum(z_scores_np_r < -1.96 &
                                                         !is.na(z_scores_np_r))
  total_entries_adjusted_under_vec_r[stage] <- sum(!is.na(z_scores_np_r))
  cat("number of under-represented entries in", stage, "is:",
      num_entries_above_threshold_under_np_r, "\n")
  percentage_above_threshold_under_np_r <-
    (num_entries_above_threshold_under_np_r /
       total_entries_adjusted_under_np_r) * 100
  cat("percentage of under-represented entries in", stage, "is:",
      percentage_above_threshold_under_np_r, "%\n\n")
  ##most over-represented and under-represented node positions##
  column_means_np_r <- colMeans(z_scores_np_r)
  max_mean_column_np_r <- names(column_means_np_r)[which.max(column_means_np_r)]
  min_mean_column_np_r <- names(column_means_np_r)[which.min(column_means_np_r)]
  cat("the most over-represented node position in", stage, "is",
      max_mean_column_np_r, "\n")
  cat("the most under-represented node position in", stage, "is",
      min_mean_column_np_r, "\n")
}

## number of entries above threshold in S1 is: 309

## percentage of entries above threshold in S1 is: 71.36259 %

## number of over-represented entries in S1 is: 116

## percentage of over-represented entries in S1 is: 26.78984 %

## number of under-represented entries in S1 is: 194

## percentage of under-represented entries in S1 is: 44.57275 %

## the most over-represented node position in S1 is np6

## the most under-represented node position in S1 is np24

## number of entries above threshold in S2 is: 593

## percentage of entries above threshold in S2 is: 70.01181 %

## number of over-represented entries in S2 is: 237
```

```
## percentage of over-represented entries in S2 is: 27.98111 %

## number of under-represented entries in S2 is: 356

## percentage of under-represented entries in S2 is: 42.0307 %

## the most over-represented node position in S2 is np34

## the most under-represented node position in S2 is np28

## number of entries above threshold in S3 is: 438

## percentage of entries above threshold in S3 is: 66.87023 %

## number of over-represented entries in S3 is: 195

## percentage of over-represented entries in S3 is: 29.77099 %

## number of under-represented entries in S3 is: 243

## percentage of under-represented entries in S3 is: 37.09924 %

## the most over-represented node position in S3 is np34

## the most under-represented node position in S3 is np24

## number of entries above threshold in S4 is: 120

## percentage of entries above threshold in S4 is: 42.25352 %

## number of over-represented entries in S4 is: 64

## percentage of over-represented entries in S4 is: 22.53521 %

## number of under-represented entries in S4 is: 56

## percentage of under-represented entries in S4 is: 19.71831 %

## the most over-represented node position in S4 is np6

## the most under-represented node position in S4 is np18

##percentage of entries with |Z|>1.96 across all stages##
num_entries_above_threshold_np_tot_r <- sum(num_entries_above_threshold_vec_r)
total_entries_adjusted_np_tot_r <- sum(total_entries_adjusted_vec_r)
percentage_above_threshold_np_tot_r <- (num_entries_above_threshold_np_tot_r /
total_entries_adjusted_np_tot_r) * 100
cat("number of entries above threshold across all stages: ",
    num_entries_above_threshold_np_tot_r, "\n")

## number of entries above threshold across all stages:  1460

cat("percentage of entries above threshold across all stages: ",
    percentage_above_threshold_np_tot_r, "%\n")
```

```
## percentage of entries above threshold across all stages:  65.7954 %

##percentage of entries with Z>1.96 across all stages##
num_entries_above_threshold_np_over_tot_r <-
  sum(num_entries_above_threshold_over_vec_r)
total_entries_adjusted_np_over_tot_r <-
  sum(total_entries_adjusted_over_vec_r)
percentage_above_threshold_np_over_tot_r <-
  (num_entries_above_threshold_np_over_tot_r /
     total_entries_adjusted_np_over_tot_r) * 100
cat("number of over-represented entries across all stages: ",
    num_entries_above_threshold_np_over_tot_r, "\n")

## number of over-represented entries across all stages:  612

cat("percentage of over-represented entries across all stages: ",
    percentage_above_threshold_np_over_tot_r, "%\n")

## percentage of over-represented entries across all stages:  27.57999 %

##percentage of entries with Z<-1.96 across all stages##
num_entries_above_threshold_np_under_tot_r <-
  sum(num_entries_above_threshold_under_vec_r)
total_entries_adjusted_np_under_tot_r <- sum(total_entries_adjusted_under_vec_r)
percentage_above_threshold_np_under_tot_r <-
  (num_entries_above_threshold_np_under_tot_r /
     total_entries_adjusted_np_under_tot_r) * 100
cat("number of under-represented entries across all stages: ",
    num_entries_above_threshold_np_under_tot_r, "\n")

## number of under-represented entries across all stages:  848

cat("percentage of under-represented entries across all stages: ",
    percentage_above_threshold_np_under_tot_r, "%\n")

## percentage of under-represented entries across all stages:  38.21541 %

###"c0_ind"###
##initialise vectors to store entries above treshold##
#total (|Z|>1.96)
num_entries_above_threshold_vec_c <- numeric(length(stage_names))
total_entries_adjusted_vec_c <- numeric(length(stage_names))
#over (Z>1.96)
num_entries_above_threshold_over_vec_c <- numeric(length(stage_names))
total_entries_adjusted_over_vec_c <- numeric(length(stage_names))
#under (Z<-1.96)
num_entries_above_threshold_under_vec_c <- numeric(length(stage_names))
total_entries_adjusted_under_vec_c <- numeric(length(stage_names))
##initialise list to store Z-scores for each stage##
z_scores_np_list_c <- list()
```

```r
##randomise networks, calculate node positions and calculate Z-scores##
#loop across stages
for (stage in stage_names) {
  #randomise
  bipnet_np_null_c <- vegan::nullmodel(bipnet_stages_list[[stage]], "c0_ind")
  bipnet_np_null_2_c <- simulate(bipnet_np_null_c, nsim = 100, seed = 1)
  bipnet_np_null_2_list_c <- lapply(seq_len(100),
                                    function(index)
                                      bipnet_np_null_2_c[, , index])
  bipnet_np_null_2_list_c <- lapply(bipnet_np_null_2_list_c, function(mat) {
    rownames(mat) <- NULL
    colnames(mat) <- NULL
    return(mat)
  })
  #initialise a list to store results
  matrix_list_np_c <- list()
  #calculate contribution to node positions
  for (i in 1:100) {
    result_matrix_np_c <- node_positions(bipnet_np_null_2_list_c[[i]],
                                         six_node = FALSE,
                                         level = "rows",
                                         weights_method = "contribution",
                                         weights_combine = "mean",
                                         normalisation = "none")
    result_matrix_np_c[is.na(result_matrix_np_c)] <- 0
    matrix_list_np_c[[i]] <- result_matrix_np_c
  }
  #get the right np list
  np_list_c <- get(paste0("np_", stage), envir = .GlobalEnv)
  #create mean and sd matrices
  mean_matrix_np_c <- apply(array(unlist(matrix_list_np_c),
                                  c(dim(matrix_list_np_c[[1]]),
                                    length(matrix_list_np_c))), c(1, 2), mean)
  sd_matrix_np_c <- apply(array(unlist(matrix_list_np_c),
                                c(dim(matrix_list_np_c[[1]]),
                                  length(matrix_list_np_c))), c(1, 2), sd)
  #calculate mean and sd
  results_mean_np_c <- as.data.frame(mean_matrix_np_c, row.names =
                                       rownames(np_list_c[["contribution"]]),
                                     col.names =
                                       colnames(np_list_c[["contribution"]]))
  results_sd_np_c <- as.data.frame(sd_matrix_np_c, row.names =
                                     rownames(np_list_c[["contribution"]]),
                                   col.names =
                                     colnames(np_list_c[["contribution"]]))
  #calculate Z-scores and upload the list
  np_0_c <- np_list_c[["contribution"]]
  np_0_c[is.na(np_0_c)] <- 0
  z_scores_np_c <- ((np_0_c - results_mean_np_c) / (results_sd_np_c))
  z_scores_np_c <- as.data.frame(z_scores_np_c, row.names = rownames(np_0_c),
                                 col.names = colnames(np_0_c))
```

```
z_scores_np_c[z_scores_np_c == -Inf] <- NaN
z_scores_np_c[z_scores_np_c == Inf] <- NaN
z_scores_np_list_c[[paste0("z_scores_np_c_", stage)]] <- z_scores_np_c
##percentage of entries with |Z|>1.96##
num_entries_above_threshold_np_c <- sum(abs(z_scores_np_c) > 1.96 &
                                        !is.na(z_scores_np_c))
total_entries_adjusted_np_c <- sum(!is.na(z_scores_np_c))
num_entries_above_threshold_vec_c[stage] <- sum(abs(z_scores_np_c) > 1.96 &
                                                !is.na(z_scores_np_c))
total_entries_adjusted_vec_c[stage] <- sum(!is.na(z_scores_np_c))
cat("number of entries above threshold in", stage, "is:",
    num_entries_above_threshold_np_c, "\n")
percentage_above_threshold_np_c <- (num_entries_above_threshold_np_c /
                                      total_entries_adjusted_np_c) * 100
cat("percentage of entries above threshold in", stage, "is:",
    percentage_above_threshold_np_c, "%\n\n")
##percentage of entries with Z>1.96##
num_entries_above_threshold_over_np_c <- sum(z_scores_np_c > 1.96 &
                                               !is.na(z_scores_np_c))
total_entries_adjusted_over_np_c <- sum(!is.na(z_scores_np_c))
num_entries_above_threshold_over_vec_c[stage] <- sum(z_scores_np_c > 1.96 &
                                                       !is.na(z_scores_np_c))
total_entries_adjusted_over_vec_c[stage] <- sum(!is.na(z_scores_np_c))
cat("number of over-represented entries in", stage, "is:",
    num_entries_above_threshold_over_np_c, "\n")
percentage_above_threshold_over_np_c <-
  (num_entries_above_threshold_over_np_c /
     total_entries_adjusted_over_np_c) * 100
cat("percentage of over-represented entries in", stage, "is:",
    percentage_above_threshold_over_np_c, "%\n\n")
##percentage of entries with Z<-1.96##
num_entries_above_threshold_under_np_c <- sum(z_scores_np_c < -1.96 &
                                                !is.na(z_scores_np_c))
total_entries_adjusted_under_np_c <- sum(!is.na(z_scores_np_c))
num_entries_above_threshold_under_vec_c[stage] <- sum(z_scores_np_c < -1.96 &
                                                        !is.na(z_scores_np_c))
total_entries_adjusted_under_vec_c[stage] <- sum(!is.na(z_scores_np_c))
cat("number of under-represented entries in", stage, "is:",
    num_entries_above_threshold_under_np_c, "\n")
percentage_above_threshold_under_np_c <-
  (num_entries_above_threshold_under_np_c /
     total_entries_adjusted_under_np_c) * 100
cat("percentage of under-represented entries in", stage, "is:",
    percentage_above_threshold_under_np_c, "%\n\n")
##most over-represented and under-represented node positions##
column_means_np_c <- colMeans(z_scores_np_c)
max_mean_column_np_c <- names(column_means_np_c)[which.max(column_means_np_c)]
min_mean_column_np_c <- names(column_means_np_c)[which.min(column_means_np_c)]
cat("the most over-represented node position in", stage, "is",
    max_mean_column_np_c, "\n")
cat("the most under-represented node position in", stage, "is",
```

```
      min_mean_column_np_c, "\n")
}

## number of entries above threshold in S1 is: 384

## percentage of entries above threshold in S1 is: 73.56322 %

## number of over-represented entries in S1 is: 63

## percentage of over-represented entries in S1 is: 12.06897 %

## number of under-represented entries in S1 is: 321

## percentage of under-represented entries in S1 is: 61.49425 %

## the most over-represented node position in S1 is np18

## the most under-represented node position in S1 is np35

## number of entries above threshold in S2 is: 637

## percentage of entries above threshold in S2 is: 66.28512 %

## number of over-represented entries in S2 is: 123

## percentage of over-represented entries in S2 is: 12.79917 %

## number of under-represented entries in S2 is: 514

## percentage of under-represented entries in S2 is: 53.48595 %

## the most over-represented node position in S2 is np6

## the most under-represented node position in S2 is np35

## number of entries above threshold in S3 is: 495

## percentage of entries above threshold in S3 is: 68.46473 %

## number of over-represented entries in S3 is: 105

## percentage of over-represented entries in S3 is: 14.52282 %

## number of under-represented entries in S3 is: 390

## percentage of under-represented entries in S3 is: 53.94191 %

## the most over-represented node position in S3 is np6

## the most under-represented node position in S3 is np44

## number of entries above threshold in S4 is: 226

## percentage of entries above threshold in S4 is: 61.58038 %

## number of over-represented entries in S4 is: 21
```

```
## percentage of over-represented entries in S4 is: 5.722071 %

## number of under-represented entries in S4 is: 205

## percentage of under-represented entries in S4 is: 55.85831 %

## the most over-represented node position in S4 is np25

## the most under-represented node position in S4 is np41

##percentage of entries with |Z|>1.96 across all stages##
num_entries_above_threshold_np_tot_c <- sum(num_entries_above_threshold_vec_c)
total_entries_adjusted_np_tot_c <- sum(total_entries_adjusted_vec_c)
percentage_above_threshold_np_tot_c <- (num_entries_above_threshold_np_tot_c /
total_entries_adjusted_np_tot_c) * 100
cat("number of entries above threshold across all stages: ",
    num_entries_above_threshold_np_tot_c, "\n")

## number of entries above threshold across all stages:  1742

cat("percentage of entries above threshold across all stages: ",
    percentage_above_threshold_np_tot_c, "%\n")

## percentage of entries above threshold across all stages:  67.70307 %

##percentage of entries with Z>1.96 across all stages##
num_entries_above_threshold_np_over_tot_c <-
  sum(num_entries_above_threshold_over_vec_c)
total_entries_adjusted_np_over_tot_c <- sum(total_entries_adjusted_over_vec_c)
percentage_above_threshold_np_over_tot_c <-
  (num_entries_above_threshold_np_over_tot_c /
     total_entries_adjusted_np_over_tot_c) * 100
cat("number of over-represented entries across all stages: ",
    num_entries_above_threshold_np_over_tot_c, "\n")

## number of over-represented entries across all stages:  312

cat("percentage of over-represented entries across all stages: ",
    percentage_above_threshold_np_over_tot_c, "%\n")

## percentage of over-represented entries across all stages:  12.12592 %

##percentage of entries with Z<-1.96 across all stages##
num_entries_above_threshold_np_under_tot_c <-
  sum(num_entries_above_threshold_under_vec_c)
total_entries_adjusted_np_under_tot_c <- sum(total_entries_adjusted_under_vec_c)
percentage_above_threshold_np_under_tot_c <-
  (num_entries_above_threshold_np_under_tot_c /
     total_entries_adjusted_np_under_tot_c) * 100
cat("number of under-represented entries across all stages: ",
    num_entries_above_threshold_np_under_tot_c, "\n")
```

```r
## number of under-represented entries across all stages:  1430

cat("percentage of under-represented entries across all stages: ",
    percentage_above_threshold_np_under_tot_c, "%\n")

## percentage of under-represented entries across all stages:  55.57715 %

###"vaznull"###
##initialise vectors to store entries above treshold##
#total (|Z|>1.96)
num_entries_above_threshold_vec_v <- numeric(length(stage_names))
total_entries_adjusted_vec_v <- numeric(length(stage_names))
#over (Z>1.96)
num_entries_above_threshold_over_vec_v <- numeric(length(stage_names))
total_entries_adjusted_over_vec_v <- numeric(length(stage_names))
#under (Z<-1.96)
num_entries_above_threshold_under_vec_v <- numeric(length(stage_names))
total_entries_adjusted_under_vec_v <- numeric(length(stage_names))
##initialise list to store Z-scores for each stage##
z_scores_np_list_v <- list()
##randomise networks, calculate node positions and calculate Z-scores##
#loop across stages
for (stage in stage_names) {
  #randomise
  set.seed(1)
  bipnet_np_null_v <- vaznull(100, bipnet_stages_list[[stage]])
  #initialise a list to store results
  matrix_list_np_v <- list()
  #calculate contribution to node positions
  for (i in 1:100) {
    result_matrix_np_v <- node_positions(bipnet_np_null_v[[i]],
                                         six_node = FALSE,
                                         level = "rows",
                                         weights_method = "contribution",
                                         weights_combine = "mean",
                                         normalisation = "none")
    result_matrix_np_v[is.na(result_matrix_np_v)] <- 0
    matrix_list_np_v[[i]] <- result_matrix_np_v
  }
  #get the right np list
  np_list_v <- get(paste0("np_", stage), envir = .GlobalEnv)
  #create mean and sd matrices
  mean_matrix_np_v <- apply(array(unlist(matrix_list_np_v),
                                  c(dim(matrix_list_np_v[[1]]),
                                    length(matrix_list_np_v))), c(1, 2), mean)
  sd_matrix_np_v <- apply(array(unlist(matrix_list_np_v),
                                c(dim(matrix_list_np_v[[1]]),
                                  length(matrix_list_np_v))), c(1, 2), sd)
  #calculate mean and sd
```

```r
results_mean_np_v <- as.data.frame(mean_matrix_np_v, row.names =
                                     rownames(np_list_v[["contribution"]]),
                                   col.names =
                                     colnames(np_list_v[["contribution"]]))
results_sd_np_v <- as.data.frame(sd_matrix_np_v, row.names =
                                   rownames(np_list_v[["contribution"]]),
                                 col.names =
                                   colnames(np_list_v[["contribution"]]))
#calculate Z-scores and upload the list
np_0_v <- np_list_v[["contribution"]]
np_0_v[is.na(np_0_v)] <- 0
z_scores_np_v <- ((np_0_v - results_mean_np_v) / (results_sd_np_v))
z_scores_np_v <- as.data.frame(z_scores_np_v, row.names = rownames(np_0_v),
                               col.names = colnames(np_0_v))
z_scores_np_v[z_scores_np_v == -Inf] <- NaN
z_scores_np_v[z_scores_np_v == Inf] <- NaN
z_scores_np_list_v[[paste0("z_scores_np_v_", stage)]] <- z_scores_np_v
##percentage of entries with |Z|>1.96##
num_entries_above_threshold_np_v <- sum(abs(z_scores_np_v) > 1.96 &
                                          !is.na(z_scores_np_v))
total_entries_adjusted_np_v <- sum(!is.na(z_scores_np_v))
num_entries_above_threshold_vec_v[stage] <- sum(abs(z_scores_np_v) > 1.96 &
                                                  !is.na(z_scores_np_v))
total_entries_adjusted_vec_v[stage] <- sum(!is.na(z_scores_np_v))
cat("number of entries above threshold in", stage, "is:",
    num_entries_above_threshold_np_v, "\n")
percentage_above_threshold_np_v <- (num_entries_above_threshold_np_v /
                                      total_entries_adjusted_np_v) * 100
cat("percentage of entries above threshold in", stage, "is:",
    percentage_above_threshold_np_v, "%\n\n")
##percentage of entries with Z>1.96##
num_entries_above_threshold_over_np_v <- sum(z_scores_np_v > 1.96 &
                                               !is.na(z_scores_np_v))
total_entries_adjusted_over_np_v <- sum(!is.na(z_scores_np_v))
num_entries_above_threshold_over_vec_v[stage] <- sum(z_scores_np_v > 1.96 &
                                                       !is.na(z_scores_np_v))
total_entries_adjusted_over_vec_v[stage] <- sum(!is.na(z_scores_np_v))
cat("number of over-represented entries in", stage, "is:",
    num_entries_above_threshold_over_np_v, "\n")
percentage_above_threshold_over_np_v <-
  (num_entries_above_threshold_over_np_v /
     total_entries_adjusted_over_np_v) * 100
cat("percentage of over-represented entries in", stage, "is:",
    percentage_above_threshold_over_np_v, "%\n\n")
##percentage of entries with Z<-1.96##
num_entries_above_threshold_under_np_v <- sum(z_scores_np_v < -1.96 &
                                                !is.na(z_scores_np_v))
total_entries_adjusted_under_np_v <- sum(!is.na(z_scores_np_v))
num_entries_above_threshold_under_vec_v[stage] <- sum(z_scores_np_v < -1.96 &
                                                        !is.na(z_scores_np_v))
total_entries_adjusted_under_vec_v[stage] <- sum(!is.na(z_scores_np_v))
```

```
  cat("number of under-represented entries in", stage, "is:",
      num_entries_above_threshold_under_np_v, "\n")
  percentage_above_threshold_under_np_v <-
    (num_entries_above_threshold_under_np_v /
       total_entries_adjusted_under_np_v) * 100
  cat("percentage of under-represented entries in", stage, "is:",
      percentage_above_threshold_under_np_v, "%\n\n")
  ##most over-represented and under-represented node positions##
  column_means_np_v <- colMeans(z_scores_np_v)
  max_mean_column_np_v <- names(column_means_np_v)[which.max(column_means_np_v)]
  min_mean_column_np_v <- names(column_means_np_v)[which.min(column_means_np_v)]
  cat("the most over-represented node position in", stage, "is",
      max_mean_column_np_v, "\n")
  cat("the most under-represented node position in", stage, "is",
      min_mean_column_np_v, "\n")
}

## number of entries above threshold in S1 is: 68

## percentage of entries above threshold in S1 is: 12.01413 %

## number of over-represented entries in S1 is: 47

## percentage of over-represented entries in S1 is: 8.303887 %

## number of under-represented entries in S1 is: 21

## percentage of under-represented entries in S1 is: 3.710247 %

## the most over-represented node position in S1 is np41

## the most under-represented node position in S1 is np25

## number of entries above threshold in S2 is: 175

## percentage of entries above threshold in S2 is: 16.68255 %

## number of over-represented entries in S2 is: 128

## percentage of over-represented entries in S2 is: 12.2021 %

## number of under-represented entries in S2 is: 47

## percentage of under-represented entries in S2 is: 4.480458 %

## the most over-represented node position in S2 is np34

## the most under-represented node position in S2 is np25

## number of entries above threshold in S3 is: 134

## percentage of entries above threshold in S3 is: 16.68742 %

## number of over-represented entries in S3 is: 106
```

```
## percentage of over-represented entries in S3 is: 13.2005 %

## number of under-represented entries in S3 is: 28

## percentage of under-represented entries in S3 is: 3.486924 %

## the most over-represented node position in S3 is np34

## the most under-represented node position in S3 is np25

## number of entries above threshold in S4 is: 35

## percentage of entries above threshold in S4 is: 9.067358 %

## number of over-represented entries in S4 is: 21

## percentage of over-represented entries in S4 is: 5.440415 %

## number of under-represented entries in S4 is: 14

## percentage of under-represented entries in S4 is: 3.626943 %

## the most over-represented node position in S4 is np34

## the most under-represented node position in S4 is np22

##percentage of entries with |Z|>1.96 across all stages##
num_entries_above_threshold_np_tot_v <- sum(num_entries_above_threshold_vec_v)
total_entries_adjusted_np_tot_v <- sum(total_entries_adjusted_vec_v)
percentage_above_threshold_np_tot_v <- (num_entries_above_threshold_np_tot_v /
total_entries_adjusted_np_tot_v) * 100
cat("number of entries above threshold across all stages: ",
    num_entries_above_threshold_np_tot_v, "\n")

## number of entries above threshold across all stages:  412

cat("percentage of entries above threshold across all stages: ",
    percentage_above_threshold_np_tot_v, "%\n")

## percentage of entries above threshold across all stages:  14.6933 %

##percentage of entries with Z>1.96 across all stages##
num_entries_above_threshold_np_over_tot_v <-
  sum(num_entries_above_threshold_over_vec_v)
total_entries_adjusted_np_over_tot_v <- sum(total_entries_adjusted_over_vec_v)
percentage_above_threshold_np_over_tot_v <-
  (num_entries_above_threshold_np_over_tot_v /
     total_entries_adjusted_np_over_tot_v) * 100
cat("number of over-represented entries across all stages: ",
    num_entries_above_threshold_np_over_tot_v, "\n")

## number of over-represented entries across all stages:  302
```

```r
cat("percentage of over-represented entries across all stages: ",
    percentage_above_threshold_np_over_tot_v, "%\n")

## percentage of over-represented entries across all stages:  10.77033 %

##percentage of entries with Z<-1.96 across all stages##
num_entries_above_threshold_np_under_tot_v <-
  sum(num_entries_above_threshold_under_vec_v)
total_entries_adjusted_np_under_tot_v <- sum(total_entries_adjusted_under_vec_v)
percentage_above_threshold_np_under_tot_v <-
  (num_entries_above_threshold_np_under_tot_v /
     total_entries_adjusted_np_under_tot_v) * 100
cat("number of under-represented entries across all stages: ",
    num_entries_above_threshold_np_under_tot_v, "\n")

## number of under-represented entries across all stages:  110

cat("percentage of under-represented entries across all stages: ",
    percentage_above_threshold_np_under_tot_v, "%\n")

## percentage of under-represented entries across all stages:  3.922967 %
```

After calculating plant species' contributions to node positions in both observed and null model networks, we proceed to perform an NMDS on all possible node positions up to 5 nodes in observed networks (in this case N=23, i.e. all possible positions, and n=133, i.e. plant species), evaluating via PERMANOVA and multivariate homogeneity of groups' dispersions whether different stages, species and interaction types (based on Shannon index of interactions) are significantly different in terms of contribution to node positions.

```r
###ORDINATION###
##prepare dataframe for ordination##
nmds_np_data <- rbind(df_S1, df_S2, df_S3, df_S4)
#add stages and species to a copy of nmds_np_data
stage_names_extended <- c("Stage1", "Stage2", "Stage3", "Stage4")
nmds_np_data_2 <- nmds_np_data
nmds_np_data_2$Stage <- rep(stage_names_extended,
                            times = sapply(list(df_S1, df_S2, df_S3, df_S4),
                                           nrow))
nmds_np_data_2$Species <- c(rownames(df_S1), rownames(df_S2), rownames(df_S3),
                            rownames(df_S4))
#transform NAs into zeros
nmds_np_data <- nmds_np_data %>% mutate_all(~ifelse(is.na(.), 0, .))
nmds_np_data_2 <- nmds_np_data_2 %>% mutate_all(~ifelse(is.na(.), 0, .))
#filter out columns with only zeros (node positions occupied by pollinators)
nmds_np_data <- nmds_np_data %>%
  select_if(~!all(. == 0))
nmds_np_data_2 <- nmds_np_data_2 %>%
  select_if(~!all(. == 0))
#calculate Shannon index of each plant species' interactions and upload values to
nmds_np_data
```

```
bipnet_stages_shannon <- bipnet_stages %>%
  group_by(webID, lower) %>%
  mutate(pij = freq / sum(freq),
         pij_ln_pij = pij * log(pij)) %>%
  ungroup()
bipnet_stages_shannon %>% #calculate -sum(pij*ln(pij))
  group_by(lower, webID) %>%
  summarise(sum_pij_ln_pij = -sum(pij_ln_pij)) %>%
  ungroup() -> summed_values
summed_values %>% #calculate the mean across stages
  group_by(lower) %>%
  summarise(mean_shannon = mean(sum_pij_ln_pij)) %>%
  ungroup() -> mean_values
summed_values <- summed_values %>% #rearrange summed_values to match
nmds_np_data_2
  arrange(webID, lower)
nmds_np_data_2$Shannon <- summed_values$sum_pij_ln_pij #upload Shannon values into
nmds_np_data
nmds_np_data_2 <- nmds_np_data_2 %>% #upload mean Shannon values across stages
into nmds_np_data
  left_join(mean_values, by = c("Species" = "lower"))
colnames(nmds_np_data_2)[ncol(nmds_np_data_2)] <- "Shannon_Mean"
#add interaction diversity labels (maximising the separation between groups)
nmds_np_data_2 <- nmds_np_data_2 %>%
  mutate(
    Shannon_Type = case_when(
      Shannon == 0 ~ "Low Interaction Diversity",
      Shannon > 0 & Shannon < 1.5 ~ "Moderate Interaction Diversity",
      Shannon >= 1.5 ~ "High Interaction Diversity"
    ),
    Shannon_Mean_Type = case_when(
      Shannon_Mean == 0 ~ "Low Interaction Diversity",
      Shannon_Mean > 0 & Shannon_Mean < 1.5 ~ "Moderate Interaction Diversity",
      Shannon_Mean >= 1.5 ~ "High Interaction Diversity"
    )
  )
##NMDS##
set.seed(1)
nmds_np <- metaMDS(nmds_np_data, distance = "bray")

## Run 0 stress 0.03965186
## Run 1 stress 0.05489691
## Run 2 stress 0.04692779
## Run 3 stress 0.05212313
## Run 4 stress 0.04168586
## Run 5 stress 0.04692769
## Run 6 stress 0.05438637
## Run 7 stress 0.04692771
## Run 8 stress 0.03965171
## ... New best solution
## ... Procrustes: rmse 4.377078e-05  max resid 0.0003772863
```

```
## ... Similar to previous best
## Run 9 stress 0.04168589
## Run 10 stress 0.03965167
## ... New best solution
## ... Procrustes: rmse 1.1936e-05  max resid 8.968702e-05
## ... Similar to previous best
## Run 11 stress 0.03965192
## ... Procrustes: rmse 4.367824e-05  max resid 0.0003817908
## ... Similar to previous best
## Run 12 stress 0.05439981
## Run 13 stress 0.05420857
## Run 14 stress 0.03965193
## ... Procrustes: rmse 4.483371e-05  max resid 0.0003822131
## ... Similar to previous best
## Run 15 stress 0.03965185
## ... Procrustes: rmse 4.189541e-05  max resid 0.0003799807
## ... Similar to previous best
## Run 16 stress 0.05058053
## Run 17 stress 0.06137396
## Run 18 stress 0.04168573
## Run 19 stress 0.05058745
## Run 20 stress 0.05211695
## *** Best solution repeated 4 times
```

```r
#diagnostic plots
plot(nmds_np)
```

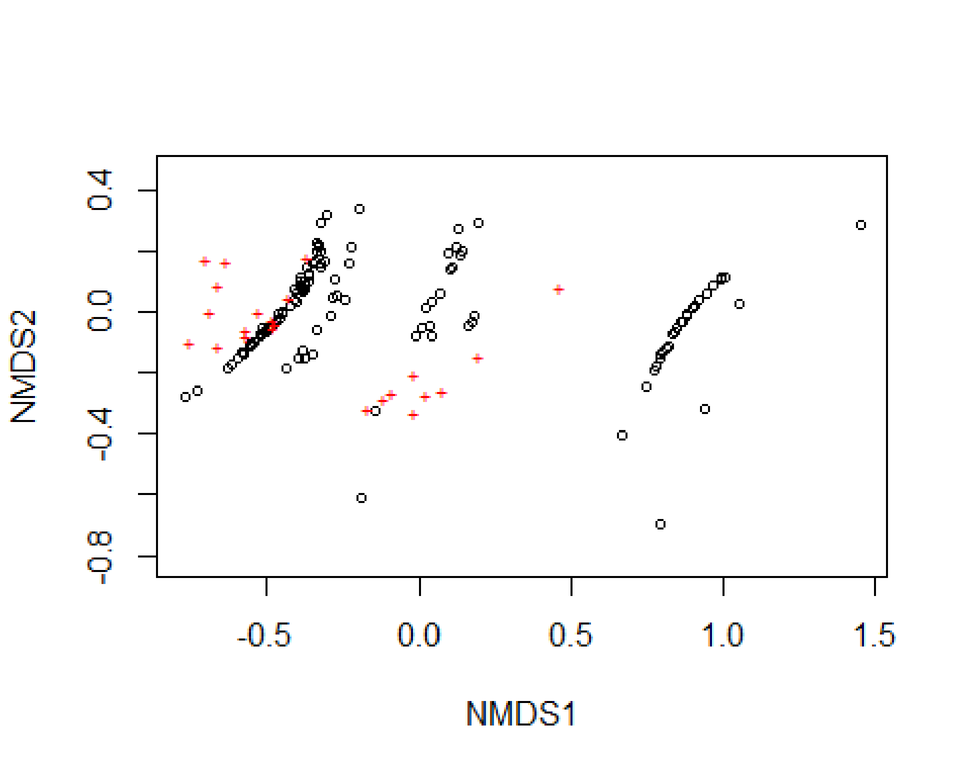

```r
stressplot(nmds_np)
```

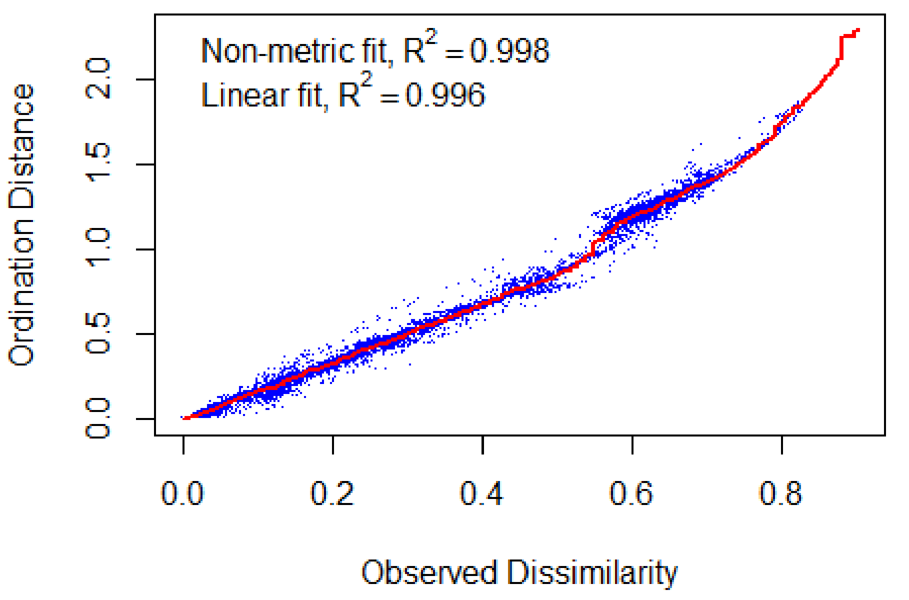

```
#stress value
nmds_np$stress

## [1] 0.03965167

#scores
scores(nmds_np)

## $sites
##                                  NMDS1        NMDS2
## Achillea erba-rotta       -0.351719736 -0.136776389
## Anthyllis vulneraria      -0.520883927 -0.075907738
## Campanula barbata          0.094080185  0.190124029
## Campanula cochleariifolia  1.449284543  0.289123253
## Cerastium alpinum         -0.360120627  0.118976844
## Cerastium arvense         -0.463703718 -0.008638566
## Epilobium fleischeri      -0.595662298 -0.153233616
## Erigeron alpinus           0.993359087  0.105928372
## Hieracium angustifolium    0.848373316 -0.052265412
## Hieracium staticifolium   -0.489421320 -0.052259218
## Leontodon hispidus        -0.365288837  0.128066392
## Leucanthemopsis alpina    -0.405110901  0.036150106
## Leucanthemum adustum       0.897876279  0.013689214
## Leucanthemum vulgare      -0.329976493  0.218499673
## Linaria alpina             0.993358189  0.105928181
## Lotus corniculatus        -0.576119301 -0.131004476
```

```
## Minuartia verna           0.877627763 -0.014976106
## Pedicularis tuberosa      0.838880116 -0.064141061
## Pilosella cymosa         -0.226335376  0.162072045
## Pilosella officinarum     0.165433596 -0.049004376
## Rumex scutatus            1.006044313  0.113099037
## Saxifraga bryoides       -0.489028472 -0.045877295
## Saxifraga paniculata     -0.540496578 -0.098207122
## Sempervivum arachnoideum  -0.143505787 -0.321786142
## Thymus praecox           -0.343480499  0.164007451
## Trifolium ochroleucon     0.197847379  0.289968557
## Trifolium pallescens     -0.324477929  0.200043871
## Achillea erba-rotta1     -0.493810649 -0.052941421
## Adenostyles alpina        0.880184580 -0.006063192
## Alchemilla monticola      0.744182606 -0.247073321
## Antennaria dioica        -0.382628909 -0.123907474
## Anthyllis vulneraria1    -0.406876045  0.070709594
## Bartsia alpina            0.791628704 -0.697543510
## Campanula barbata1        0.010273285 -0.054579623
## Campanula scheuchzeri     0.865730242 -0.034119854
## Carduus defloratus       -0.310547271  0.164473699
## Centaurea nervosa        -0.404802371  0.057785137
## Cerastium arvense1       -0.544871205 -0.100862428
## Chaerophyllum villarsii    0.802973581 -0.129239545
## Epilobium fleischeri1    -0.470967734 -0.030896917
## Erigeron alpinus1         0.880184484 -0.006063201
## Galium anisophyllon      -0.329370522  0.211315614
## Gymnadenia conopsea       0.040274719  0.035158121
## Hieracium angustifolium1   0.136562709  0.185892763
## Hieracium murorum        -0.409188900  0.039031824
## Hieracium staticifolium1  -0.477048794 -0.041641654
## Hippocrepis comosa        0.834356451 -0.074071721
## Leontodon helveticus     -0.410340825  0.042296293
## Leontodon hispidus1      -0.504957017 -0.069051558
## Leucanthemopsis alpina1    0.667094136 -0.407584166
## Leucanthemum vulgare1    -0.522737726 -0.078419020
## Lotus corniculatus1      -0.381475562  0.077039432
## Myosotis alpestris       -0.347848878  0.159148667
## Orchis mascula            0.880184587 -0.006063191
## Phyteuma betonicifolium   -0.425045397  0.018957990
## Pilosella cymosa1         0.859709317 -0.033237803
## Pilosella officinarum1   -0.456623492 -0.018882519
## Polygonum viviparum       0.944092707  0.062709127
## Potentilla aurea         -0.628059683 -0.182874181
## Ranunculus montanus      -0.434908261 -0.186652508
## Ranunculus villarsii     -0.575232192 -0.136203939
## Rhinanthus minor         -0.222902319  0.214424758
## Rhododendron ferrugineum  -0.484118140 -0.048630541
## Rumex scutatus1           0.795258136 -0.140791584
## Saxifraga aizoides       -0.617389990 -0.171240530
## Saxifraga paniculata1    -0.578535686 -0.131430628
## Silene rupestris         -0.291755113 -0.012534669
```

```
## Silene vulgaris            0.100350848  0.137222504
## Solidago virgaurea         0.108150048  0.146542596
## Thymus praecox1           -0.375816243 -0.155330851
## Tofieldia calyculata       0.142869506  0.198469030
## Trifolium badium          -0.338828942  0.198293337
## Trifolium hybridum         0.880184618 -0.006063188
## Trifolium ochroleucon1     0.033734774 -0.044655147
## Trifolium pallescens1     -0.323113907  0.161101977
## Trifolium pratense        -0.280664865  0.044726467
## Veronica fruticans        -0.385216522  0.065320773
## Achillea erba-rotta2       0.024389218  0.013623821
## Achillea millefolium       0.124498148  0.212554548
## Anthyllis vulneraria2     -0.242808126  0.041130441
## Campanula barbata2         0.175371610 -0.033418707
## Campanula scheuchzeri1     0.069967409  0.060539238
## Carduus defloratus1        0.773974705 -0.191651266
## Centaurea nervosa1        -0.397840317 -0.152681291
## Cerastium arvense2        -0.387918612  0.078040815
## Crepis pontana             0.995536425  0.110543540
## Dryas octopetala          -0.511879331 -0.069759138
## Epilobium fleischeri2     -0.376061391  0.102978498
## Gentiana nivalis           0.965877988  0.084461036
## Hieracium murorum1        -0.507527357 -0.068572696
## Hieracium staticifolium2  -0.388557637  0.116237678
## Hippocrepis comosa1        0.179778493 -0.013400079
## Huguenina tanacetifolia   -0.469563901 -0.034848019
## Leontodon hispidus2       -0.496230420 -0.057587655
## Leucanthemum vulgare2     -0.196405371  0.340998389
## Lotus corniculatus2       -0.363468454  0.103084102
## Myosotis alpestris1       -0.373557284  0.084012602
## Peucedanum ostruthium     -0.548550844 -0.105683174
## Phyteuma betonicifolium1  -0.446318513 -0.005550737
## Pilosella officinarum2    -0.556596544 -0.113883257
## Potentilla aurea1         -0.730895297 -0.261385318
## Ranunculus montanus1      -0.381880493  0.068703417
## Ranunculus villarsii1     -0.554620730 -0.102736254
## Rhododendron ferrugineum1 -0.594391733 -0.149293483
## Sempervivum arachnoideum1  0.941943451 -0.315342691
## Silene dioica              0.128138800  0.271499666
## Silene vulgaris1          -0.337739200 -0.056452815
## Thymus praecox2           -0.302739740  0.318244660
## Trifolium hybridum1        0.795076923 -0.150885043
## Trifolium ochroleucon2    -0.368205574  0.148970161
## Trifolium pallescens2     -0.322541723  0.147242681
## Trifolium pratense1       -0.323018315  0.295564168
## Trifolium repens           0.811233912 -0.119438309
## Vaccinium vitis-idaea      0.904185577  0.020208332
## Veronica fruticans1       -0.188542067 -0.613051129
## Achillea erba-rotta3      -0.334430839  0.229112703
## Anthyllis vulneraria3      0.905296435  0.019144013
## Campanula rhomboidalis     0.781597939 -0.176896047
```

```
## Dactylorhiza maculata      0.044605660 -0.078434013
## Hieracium murorum2        -0.276041076  0.106550385
## Leontodon hispidus3       -0.448123006 -0.002983232
## Lotus corniculatus3       -0.386675528  0.101641754
## Peucedanum ostruthium1    -0.515252248 -0.050019946
## Potentilla aurea2         -0.373444874  0.072604029
## Ranunculus montanus2      -0.271131625  0.056439109
## Ranunculus platanifolius   0.922223294  0.038084337
## Ranunculus villarsii2     -0.448270378  0.001509302
## Rhododendron ferrugineum2 -0.770267695 -0.278507641
## Sempervivum montanum       1.449284546  0.289123242
## Vaccinium vitis-idaea1     0.922229501  0.038085350
## Valeriana tripteris       -0.009969862 -0.080316409
## Veronica chamaedrys        0.818347490 -0.110400529
## Vicia sepium               1.048778736  0.028677531
##
## $species
##            NMDS1        NMDS2
## np2   0.46270040  0.078229663
## np4  -0.37167970  0.177017794
## np6   0.19645036 -0.148952062
## np8  -0.02015142 -0.204506730
## np11  0.07737673 -0.257379958
## np12 -0.43166043  0.047210064
## np14 -0.52656788 -0.001187507
## np16 -0.63295733  0.168584864
## np18 -0.11609492 -0.284747237
## np21 -0.09203605 -0.263960610
## np22 -0.47966799 -0.053720916
## np24  0.02399132 -0.270668012
## np25 -0.47953171 -0.027591375
## np28 -0.16775414 -0.316227302
## np29 -0.56542697 -0.056600523
## np31 -0.66234273 -0.112769948
## np34 -0.01746261 -0.332080245
## np35 -0.65893327  0.088414224
## np38 -0.47278183 -0.039461105
## np41 -0.56770017 -0.076708392
## np42 -0.68829175 -0.002481114
## np44 -0.75261593 -0.097764482
## np46 -0.70013976  0.175449331

#create NMDS scores dataframe
nmds_np_scores <- data.frame(scores(nmds_np)$sites)
nmds_np_scores$Stage <- nmds_np_data_2$Stage
nmds_np_scores$Species <- nmds_np_data_2$Species
#calculate centroids
centroids_np_stages <- aggregate(cbind(NMDS1, NMDS2) ~ Stage,
                                 data = nmds_np_scores, FUN = mean)
centroids_np_species <- aggregate(cbind(NMDS1, NMDS2) ~ Species,
```

```r
                              data = nmds_np_scores, FUN = mean)
##test difference between ordination groups##
#adonis (stage)
set.seed(1)
adonis_np_stages <- adonis2(nmds_np_data ~ nmds_np_data_2$Stage,
                            data = nmds_np_data, permutations = 1000,
                            method = "bray")
adonis_np_stages

## Permutation test for adonis under reduced model
## Terms added sequentially (first to last)
## Permutation: free
## Number of permutations: 1000
##
## adonis2(formula = nmds_np_data ~ nmds_np_data_2$Stage, data = nmds_np_data,
permutations = 1000, method = "bray")
##                       Df SumOfSqs      R2      F Pr(>F)
## nmds_np_data_2$Stage   3   0.2602 0.02022 0.8874 0.4545
## Residual             129  12.6103 0.97978
## Total                132  12.8705 1.00000

#adonis (species)
set.seed(1)
adonis_np_species <- adonis2(nmds_np_data ~ nmds_np_data_2$Species,
                             data = nmds_np_data, permutations = 1000,
                             method = "bray")
adonis_np_species

## Permutation test for adonis under reduced model
## Terms added sequentially (first to last)
## Permutation: free
## Number of permutations: 1000
##
## adonis2(formula = nmds_np_data ~ nmds_np_data_2$Species, data = nmds_np_data,
permutations = 1000, method = "bray")
##                         Df SumOfSqs      R2    F   Pr(>F)
## nmds_np_data_2$Species  73  10.8017 0.83926 4.22 0.000999 ***
## Residual                59   2.0688 0.16074
## Total                  132  12.8705 1.00000
## ---
## Signif. codes:  0 '***' 0.001 '**' 0.01 '*' 0.05 '.' 0.1 ' ' 1

#adonis (shannon)
set.seed(1)
adonis_np_shannon <- adonis2(nmds_np_data ~ nmds_np_data_2$Shannon_Type,
                             data = nmds_np_data, permutations = 1000,
                             method = "bray")
adonis_np_shannon

## Permutation test for adonis under reduced model
```

```
## Terms added sequentially (first to last)
## Permutation: free
## Number of permutations: 1000
##
## adonis2(formula = nmds_np_data ~ nmds_np_data_2$Shannon_Type, data =
nmds_np_data, permutations = 1000, method = "bray")
##                              Df SumOfSqs      R2      F   Pr(>F)
## nmds_np_data_2$Shannon_Type   2   9.4924 0.73753 182.65 0.000999 ***
## Residual                    130   3.3781 0.26247
## Total                       132  12.8705 1.00000
## ---
## Signif. codes:  0 '***' 0.001 '**' 0.01 '*' 0.05 '.' 0.1 ' ' 1

#betadisper (stage)
distance_matrix_np <- vegdist(nmds_np_data, method = "bray")
set.seed(1)
betadisper_np_stages <- betadisper(distance_matrix_np, nmds_np_data_2$Stage)
permutest(betadisper_np_stages, pairwise = TRUE)

##
## Permutation test for homogeneity of multivariate dispersions
## Permutation: free
## Number of permutations: 999
##
## Response: Distances
##            Df Sum Sq  Mean Sq      F N.Perm Pr(>F)
## Groups      3 0.1044 0.034807 0.8875    999  0.434
## Residuals 129 5.0593 0.039219
##
## Pairwise comparisons:
## (Observed p-value below diagonal, permuted p-value above diagonal)
##         Stage1  Stage2  Stage3 Stage4
## Stage1         0.43500 0.29100  0.654
## Stage2 0.42689         0.73200  0.213
## Stage3 0.29881 0.74274          0.135
## Stage4 0.65647 0.22631 0.15037

#betadisper (species)
set.seed(1)
betadisper_np_species <- betadisper(distance_matrix_np, nmds_np_data_2$Species)
permutest(betadisper_np_species, pairwise = FALSE)

##
## Permutation test for homogeneity of multivariate dispersions
## Permutation: free
## Number of permutations: 999
##
## Response: Distances
##           Df  Sum Sq   Mean Sq      F N.Perm Pr(>F)
## Groups    73 0.93446 0.0128009 1.4931    999  0.236
## Residuals 59 0.50581 0.0085731
```

```r
#betadisper (shannon)
set.seed(1)
betadisper_np_shannon <- betadisper(distance_matrix_np,
                                    nmds_np_data_2$Shannon_Type)
permutest(betadisper_np_shannon, pairwise = TRUE)

##
## Permutation test for homogeneity of multivariate dispersions
## Permutation: free
## Number of permutations: 999
##
## Response: Distances
##            Df  Sum Sq  Mean Sq      F N.Perm Pr(>F)
## Groups      2 0.33431 0.167153 17.597    999  0.001 ***
## Residuals 130 1.23484 0.009499
## ---
## Signif. codes:  0 '***' 0.001 '**' 0.01 '*' 0.05 '.' 0.1 ' ' 1
##
## Pairwise comparisons:
## (Observed p-value below diagonal, permuted p-value above diagonal)
##                                High Interaction Diversity
## High Interaction Diversity
## Low Interaction Diversity                     5.7874e-01
## Moderate Interaction Diversity                6.8320e-07
##                                Low Interaction Diversity
## High Interaction Diversity                   5.9800e-01
## Low Interaction Diversity
## Moderate Interaction Diversity               7.4748e-05
##                                Moderate Interaction Diversity
## High Interaction Diversity                              0.001
## Low Interaction Diversity                               0.001
## Moderate Interaction Diversity
```

We model Shannon index of interactions (calculated on all species across stages and on species' centroids) in relation to NMDS1 coordinates of node positions' ordination.

```r
###SHANNON~NMDS1(NP)###
##upload Shannon_Mean values and Shannon_Mean_Types to centroids_np_species##
centroids_np_species$Shannon_Mean <- NA
centroids_np_species$Shannon_Mean_Type <- NA
for (species in centroids_np_species$Species) {
  matching_species <- which(nmds_np_data_2$Species == species)
  if (length(matching_species) > 0) {
    matching_species <- matching_species[1]
    centroids_np_species$Shannon_Mean[centroids_np_species$Species == species] <-
nmds_np_data_2$Shannon_Mean[matching_species]
    centroids_np_species$Shannon_Mean_Type[centroids_np_species$Species
                                           == species] <-
nmds_np_data_2$Shannon_Mean_Type[matching_species]
  }
```

```
}
##linear models##
lm_shannon_1 <- lm(Shannon ~ NMDS1, data = nmds_np_data_2)
summary(lm_shannon_1)

##
## Call:
## lm(formula = Shannon ~ NMDS1, data = nmds_np_data_2)
##
## Residuals:
##      Min       1Q   Median       3Q      Max
## -1.26294 -0.43766 -0.03865  0.35825  1.38112
##
## Coefficients:
##             Estimate Std. Error t value Pr(>|t|)
## (Intercept)  1.46571    0.04850   30.22   <2e-16 ***
## NMDS1       -1.67138    0.08328  -20.07   <2e-16 ***
## ---
## Signif. codes:  0 '***' 0.001 '**' 0.01 '*' 0.05 '.' 0.1 ' ' 1
##
## Residual standard error: 0.5593 on 131 degrees of freedom
## Multiple R-squared:  0.7546, Adjusted R-squared:  0.7527
## F-statistic: 402.8 on 1 and 131 DF,  p-value: < 2.2e-16

lm_shannon_2 <- lm(Shannon ~ poly(NMDS1, 2), data = nmds_np_data_2)
summary(lm_shannon_2)

##
## Call:
## lm(formula = Shannon ~ poly(NMDS1, 2), data = nmds_np_data_2)
##
## Residuals:
##      Min       1Q   Median       3Q      Max
## -1.57392 -0.27845 -0.01788  0.26538  1.12981
##
## Coefficients:
##                  Estimate Std. Error t value Pr(>|t|)
## (Intercept)       1.46571    0.03909  37.499  < 2e-16 ***
## poly(NMDS1, 2)1 -11.22509    0.45077 -24.902  < 2e-16 ***
## poly(NMDS1, 2)2   3.81602    0.45077   8.466 4.66e-14 ***
## ---
## Signif. codes:  0 '***' 0.001 '**' 0.01 '*' 0.05 '.' 0.1 ' ' 1
##
## Residual standard error: 0.4508 on 130 degrees of freedom
## Multiple R-squared:  0.8418, Adjusted R-squared:  0.8394
## F-statistic: 345.9 on 2 and 130 DF,  p-value: < 2.2e-16

lm_shannon_3 <- lm(Shannon_Mean ~ NMDS1, data = centroids_np_species)
summary(lm_shannon_3)

##
## Call:
```

```
## lm(formula = Shannon_Mean ~ NMDS1, data = centroids_np_species)
##
## Residuals:
##      Min       1Q   Median       3Q      Max
## -1.01921 -0.30337 -0.05828  0.24922  1.11854
##
## Coefficients:
##             Estimate Std. Error t value Pr(>|t|)
## (Intercept)  1.41078    0.05341   26.42   <2e-16 ***
## NMDS1       -1.50400    0.08698  -17.29   <2e-16 ***
## ---
## Signif. codes:  0 '***' 0.001 '**' 0.01 '*' 0.05 '.' 0.1 ' ' 1
##
## Residual standard error: 0.4462 on 72 degrees of freedom
## Multiple R-squared:  0.8059, Adjusted R-squared:  0.8032
## F-statistic:   299 on 1 and 72 DF,  p-value: < 2.2e-16
```

```
lm_shannon_4 <- lm(Shannon_Mean ~ poly(NMDS1, 2), data = centroids_np_species)
summary(lm_shannon_4)
```

```
##
## Call:
## lm(formula = Shannon_Mean ~ poly(NMDS1, 2), data = centroids_np_species)
##
## Residuals:
##      Min       1Q   Median       3Q      Max
## -1.28025 -0.15647 -0.05892  0.21541  0.72462
##
## Coefficients:
##                 Estimate Std. Error t value Pr(>|t|)
## (Intercept)      1.19080    0.04331  27.492  < 2e-16 ***
## poly(NMDS1, 2)1 -7.71536    0.37260 -20.707  < 2e-16 ***
## poly(NMDS1, 2)2  2.11598    0.37260   5.679 2.77e-07 ***
## ---
## Signif. codes:  0 '***' 0.001 '**' 0.01 '*' 0.05 '.' 0.1 ' ' 1
##
## Residual standard error: 0.3726 on 71 degrees of freedom
## Multiple R-squared:  0.8665, Adjusted R-squared:  0.8628
## F-statistic: 230.5 on 2 and 71 DF,  p-value: < 2.2e-16
```

```
#model selection
aic_lm_shannon <- AIC(lm_shannon_1, lm_shannon_2)
aic_lm_shannon
```

```
##              df      AIC
## lm_shannon_1  3 226.8508
## lm_shannon_2  4 170.4529
```

```
aic_lm_shannon_mean <- AIC(lm_shannon_3, lm_shannon_4)
aic_lm_shannon_mean
```

```
##               df      AIC
## lm_shannon_3  3 94.53973
## lm_shannon_4  4 68.82854

#diagnostic
shapiro.test(lm_shannon_4$residuals)

##
##  Shapiro-Wilk normality test
##
## data:  lm_shannon_4$residuals
## W = 0.93421, p-value = 0.0008171

plot(lm_shannon_4)
```

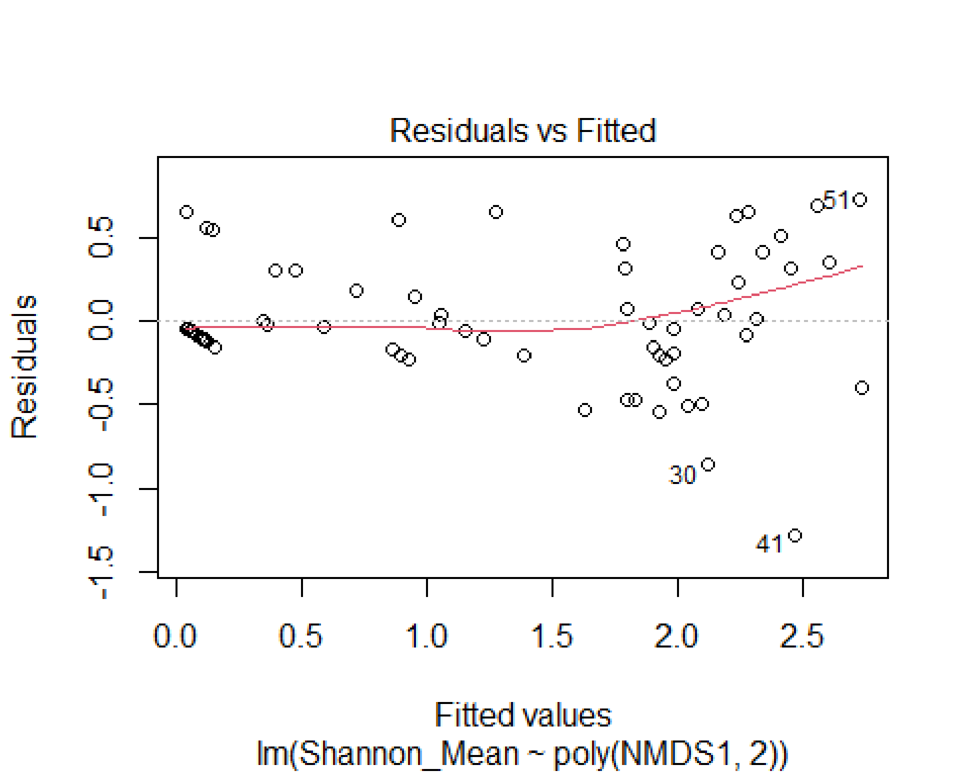

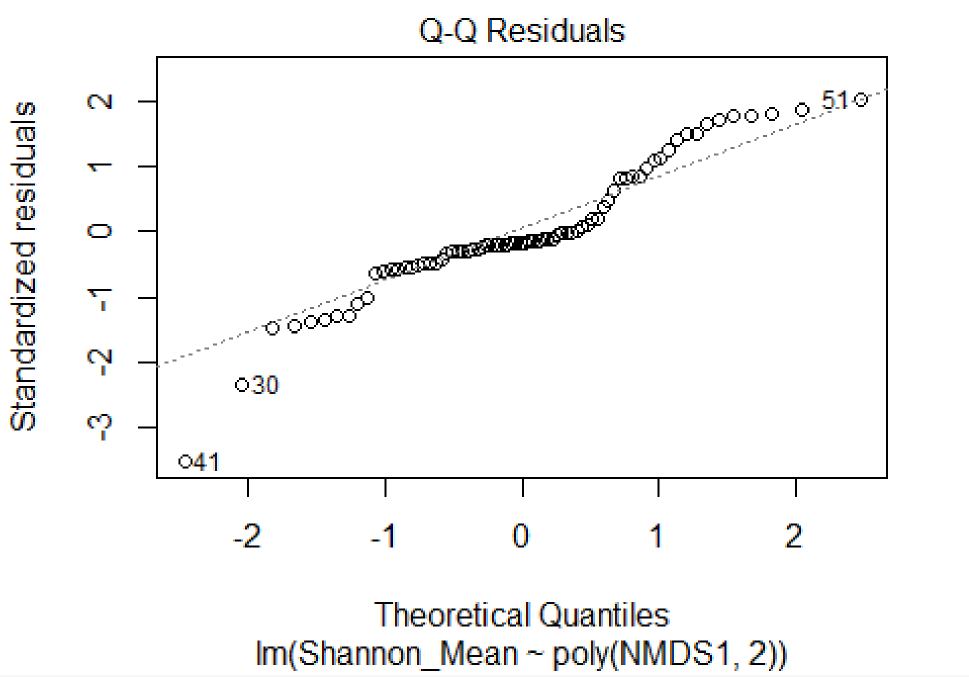
Q-Q Residuals
Standardized residuals
Theoretical Quantiles
lm(Shannon_Mean ~ poly(NMDS1, 2))
51
30
41

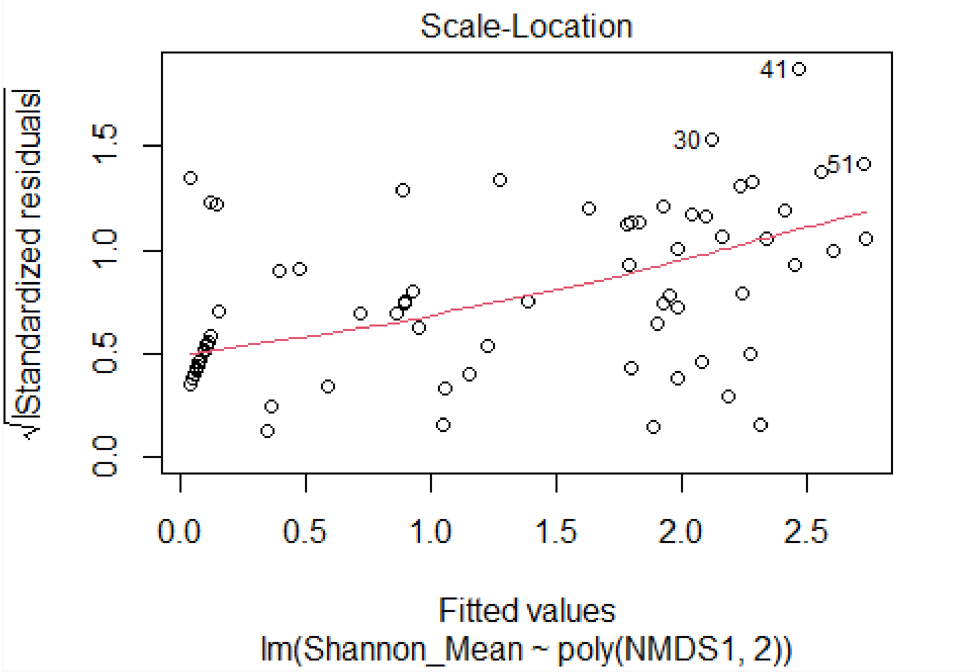
Scale-Location
√|Standardized residuals|
Fitted values
lm(Shannon_Mean ~ poly(NMDS1, 2))
41
30
51

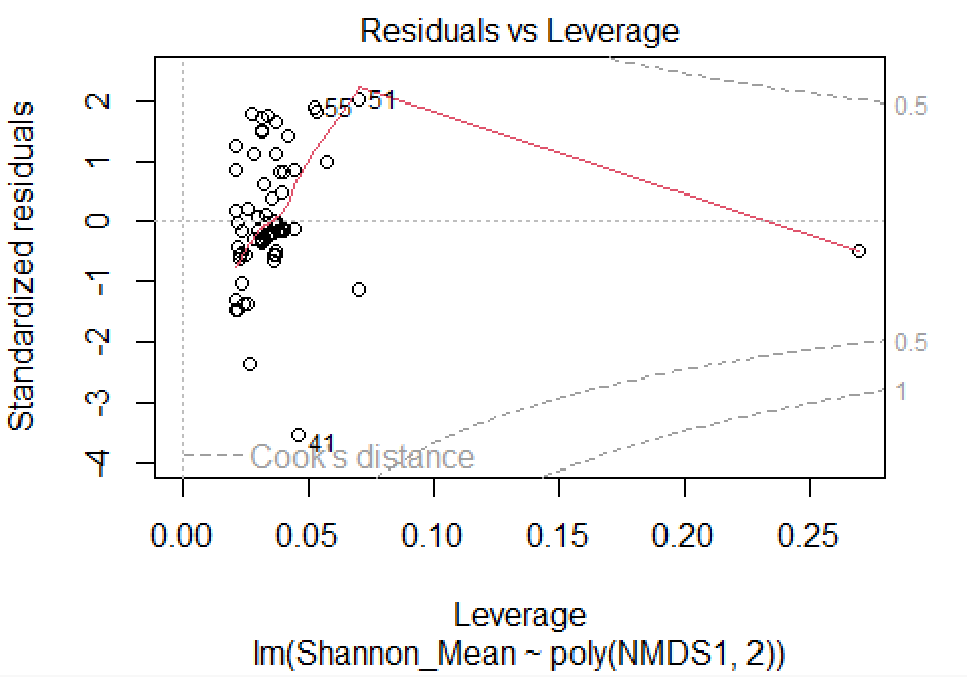

```
##generalised linear models##
#transform Shannon to handle zeros for gamma family
nmds_np_data_2$Shannon_T <- nmds_np_data_2$Shannon + 10^-3
centroids_np_species$Shannon_Mean_T <- centroids_np_species$Shannon_Mean + 10^-3
#models
glm_shannon_1 <- glm(Shannon_T ~ NMDS1, data = nmds_np_data_2,
                     family = Gamma(link = "inverse"))
summary(glm_shannon_1)

##
## Call:
## glm(formula = Shannon_T ~ NMDS1, family = Gamma(link = "inverse"),
##     data = nmds_np_data_2)
##
## Coefficients:
##             Estimate Std. Error t value Pr(>|t|)
## (Intercept)  1.33164    0.07759   17.16   <2e-16 ***
## NMDS1        1.68151    0.10923   15.39   <2e-16 ***
## ---
## Signif. codes:  0 '***' 0.001 '**' 0.01 '*' 0.05 '.' 0.1 ' ' 1
##
## (Dispersion parameter for Gamma family taken to be 0.4172312)
##
##     Null deviance: 421.91  on 132  degrees of freedom
## Residual deviance: 319.68  on 131  degrees of freedom
## AIC: 227.92
```

```
##
## Number of Fisher Scoring iterations: 9
```

```r
glm_shannon_2 <- glm(Shannon_T ~ poly(NMDS1, 2), data = nmds_np_data_2,
                     family = Gamma(link = "inverse"))
summary(glm_shannon_2)
```

```
##
## Call:
## glm(formula = Shannon_T ~ poly(NMDS1, 2), family = Gamma(link = "inverse"),
##     data = nmds_np_data_2)
##
## Coefficients:
##                   Estimate Std. Error t value Pr(>|t|)
## (Intercept)         2.8572     0.4023   7.103 7.18e-11 ***
## poly(NMDS1, 2)1   41.4698     7.0367   5.893 3.06e-08 ***
## poly(NMDS1, 2)2   10.7820     2.3642   4.560 1.17e-05 ***
## ---
## Signif. codes:  0 '***' 0.001 '**' 0.01 '*' 0.05 '.' 0.1 ' ' 1
##
## (Dispersion parameter for Gamma family taken to be 1.16729)
##
##     Null deviance: 421.91  on 132  degrees of freedom
## Residual deviance: 272.05  on 130  degrees of freedom
## AIC: 202.22
##
## Number of Fisher Scoring iterations: 8
```

```r
glm_shannon_3 <- glm(Shannon_Mean_T ~ NMDS1, data = centroids_np_species,
                     family = Gamma(link = "inverse"))
summary(glm_shannon_3)
```

```
##
## Call:
## glm(formula = Shannon_Mean_T ~ NMDS1, family = Gamma(link = "inverse"),
##     data = centroids_np_species)
##
## Coefficients:
##             Estimate Std. Error t value Pr(>|t|)
## (Intercept)   1.5134     0.1434  10.554 2.88e-16 ***
## NMDS1         2.2291     0.2629   8.478 1.97e-12 ***
## ---
## Signif. codes:  0 '***' 0.001 '**' 0.01 '*' 0.05 '.' 0.1 ' ' 1
##
## (Dispersion parameter for Gamma family taken to be 0.5738824)
##
##     Null deviance: 264.66  on 73  degrees of freedom
## Residual deviance: 199.45  on 72  degrees of freedom
## AIC: 80.037
##
## Number of Fisher Scoring iterations: 6
```

```
glm_shannon_4 <- glm(Shannon_Mean_T ~ poly(NMDS1, 2),
                     data = centroids_np_species,
                      family = Gamma(link = "inverse"))
summary(glm_shannon_4)

##
## Call:
## glm(formula = Shannon_Mean_T ~ poly(NMDS1, 2), family = Gamma(link =
"inverse"),
##     data = centroids_np_species)
##
## Coefficients:
##                  Estimate Std. Error t value Pr(>|t|)
## (Intercept)       3.1647     0.5429   5.829 1.51e-07 ***
## poly(NMDS1, 2)1  29.1299     6.0063   4.850 7.06e-06 ***
## poly(NMDS1, 2)2   9.2814     2.5653   3.618 0.000552 ***
## ---
## Signif. codes:  0 '***' 0.001 '**' 0.01 '*' 0.05 '.' 0.1 ' ' 1
##
## (Dispersion parameter for Gamma family taken to be 1.156944)
##
##     Null deviance: 264.66  on 73  degrees of freedom
## Residual deviance: 177.08  on 71  degrees of freedom
## AIC: 70.417
##
## Number of Fisher Scoring iterations: 7

#model selection
aic_glm_shannon <- AIC(glm_shannon_1, glm_shannon_2)
aic_glm_shannon

##                df      AIC
## glm_shannon_1  3 227.9248
## glm_shannon_2  4 202.2212

aic_glm_shannon_mean <- AIC(glm_shannon_3, glm_shannon_4)
aic_glm_shannon_mean

##                df      AIC
## glm_shannon_3  3 80.03678
## glm_shannon_4  4 70.41686
```

We conclude by drawing the ordination plot (grouped by stage, species and interaction diversity) and the relation between Shannon index of interactions and NMDS1 coordinates of node positions' ordination.

```
###GRAPHS###
figure_S4a <- ggplot(nmds_np_scores, aes(x = NMDS1, y = NMDS2, colour = Stage))
                     + geom_point(data = nmds_np_scores,aes(x = NMDS1,y = NMDS2,
                                                           colour = Stage),
                                 size = 2) +
                     geom_encircle(aes(group = Stage), size = 1.8)+
```

```
                    scale_colour_manual(values = c("Stage1" = "blue3",
                                                   "Stage2" = "green3",
                                                   "Stage3" = "yellow3",
                                                   "Stage4" = "red3")) +
                    coord_fixed() +
                    theme_bw() +
                    theme(axis.ticks = element_blank(),
                          axis.title.x = element_text(size = 18),
                          axis.title.y = element_text(size = 18),
                          panel.background = element_blank(),
                          plot.background = element_blank(),
                          panel.grid.major = element_blank(),
                          panel.grid.minor = element_blank()) +
                    labs(title = "A")
figure_S4a
```

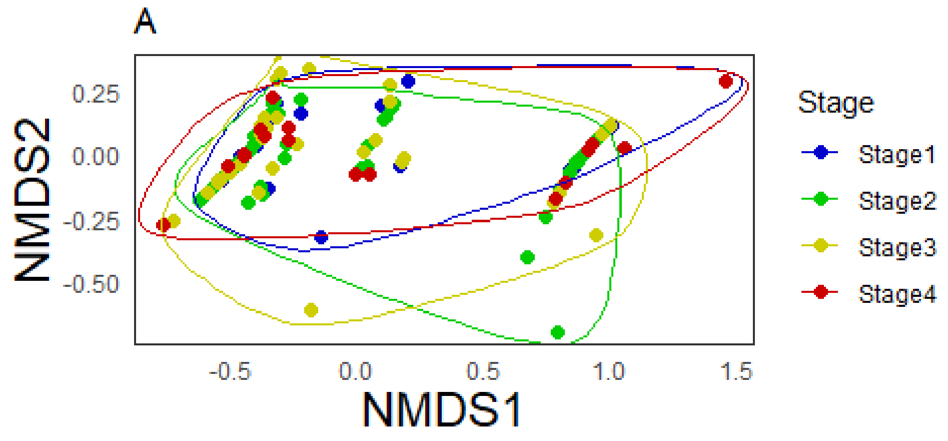

```
figure_S4b <- ggplot(centroids_np_species, aes(x = NMDS1, y = NMDS2,
                                                colour = Species)) +
                    geom_point(data = centroids_np_species, size = 2) +
                    coord_fixed() +
                    theme_bw() +
                    theme(legend.position = "none",
                          panel.grid.major = element_blank(),
                          panel.grid.minor = element_blank()) +
                    labs(title = "B")
figure_S4b
```

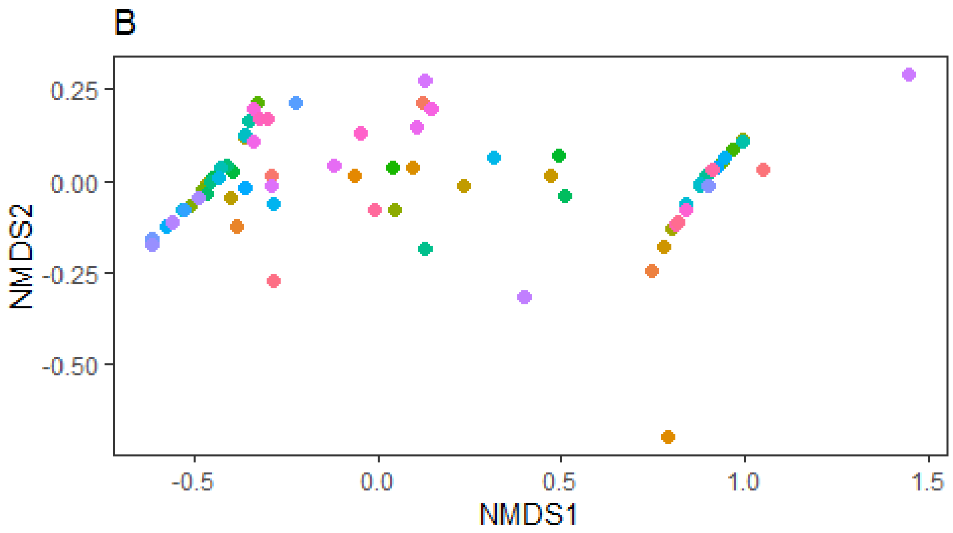

```
figure_5a <- ggplot(centroids_np_species, aes(x = NMDS1, y = NMDS2,
                                               colour = Shannon_Mean_Type)) +
                    geom_point(size = 1.5) +
                    geom_encircle(aes(group = Shannon_Mean_Type), size = 1.8) +
                    scale_colour_manual(values =
                                          c("High Interaction Diversity" =
                                              "navyblue",
                                            "Moderate Interaction Diversity" =
                                              "dodgerblue3",
                                            "Low Interaction Diversity" =
                                              "darkturquoise"),
                                        name = "Plant Species Interaction Type",
                                        breaks =
                                          c("High Interaction Diversity",
                                            "Moderate Interaction Diversity",
                                            "Low Interaction Diversity")) +
                    coord_fixed() +
                    theme_bw() +
                    theme(axis.ticks = element_blank(),
                          axis.title = element_text(size = 10),
                          panel.background = element_blank(),
                          plot.background = element_blank(),
                          panel.grid.major = element_blank(),
                          panel.grid.minor = element_blank(),
                          legend.title = element_text(size = 8),
                          legend.text = element_text(size = 7)) +
                    labs(title = "A")
```

```
figure_5a
```

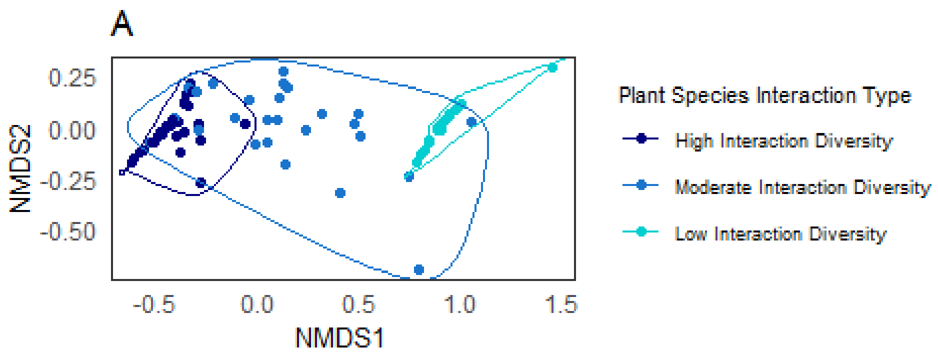

```
figure_5b <- ggplot(data = centroids_np_species,
                    mapping = aes(NMDS1, Shannon_Mean,
                                  color = Shannon_Mean_Type)) +
                    geom_point() +
                    geom_smooth(method = glm, formula = y ~ poly(x, 2),
                                color = "black", linewidth = 0.8) +
                    scale_color_manual(values =
                                         c("High Interaction Diversity" =
                                             "navyblue",
                                           "Moderate Interaction Diversity"
                                           = "dodgerblue3",
                                           "Low Interaction Diversity" =
                                             "darkturquoise"),
                                       name = "Plant Species Interaction Type",
                                       breaks =
                                         c("High Interaction Diversity",
                                           "Moderate Interaction Diversity",
                                           "Low Interaction Diversity")) +
                    theme_bw() +
                    theme(panel.grid.major = element_blank(),
                          panel.grid.minor = element_blank(),
                          legend.title = element_text(size = 8),
                          legend.text = element_text(size = 7),
                          axis.title = element_text(size = 10)) +
                    labs(x = "NMDS1 [x coords of node positions NMDS]",
```

```
                            y = "Shannon Index of interactions [H']",
                            title = "B")
figure_5b
```

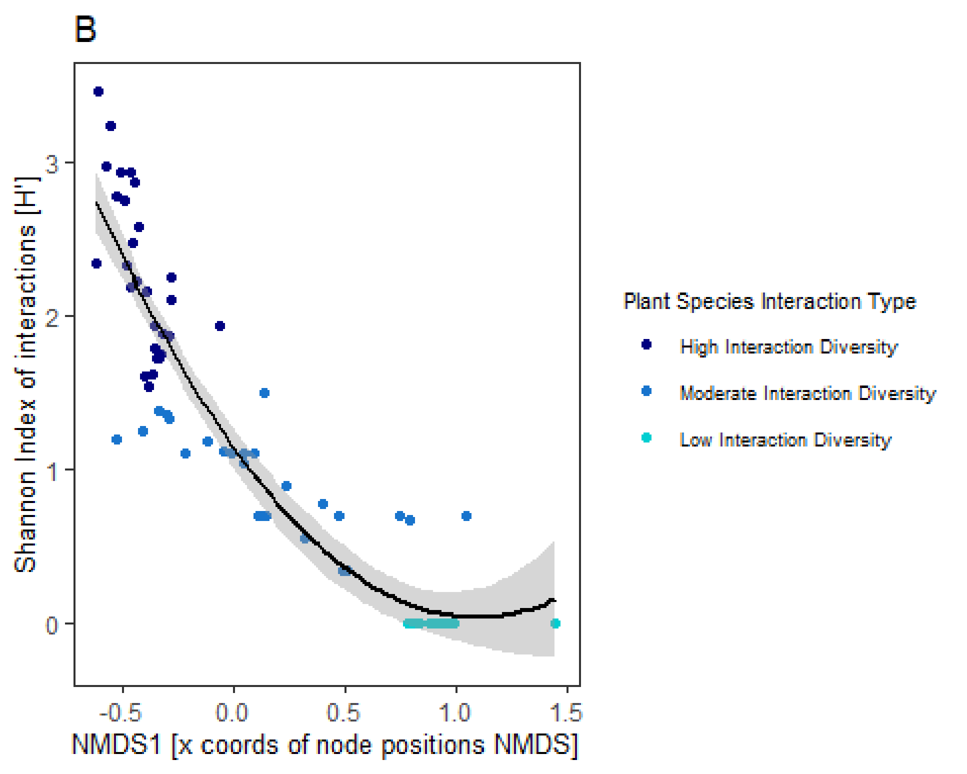
